# Defensive lipid droplets are PUFA reservoirs driving bacterial clearance and inflammation

**DOI:** 10.64898/2026.03.12.711356

**Authors:** Marta Bosch, Gyana-Lipsa Parida, Marina Sánchez-Quijada, Carles Ruiz-Mirapeix, Miguel Sánchez-Álvarez, Laura Pedró-Cos, Alba Fajardo, Harriet P. Lo, Mariano Alonso-Bivou, Rémi Safi, Erika Pineda, James Rae, James E.B. Curson, Bernhard Keller, Jesús Balsinde, Anna M. Planas, Matthew J. Sweet, Albert Herms, Caroline Demangel, Robert G. Parton, Albert Pol

## Abstract

Lipid droplets (LDs) rapidly form in infected cells to participate in the defence against microbes. Here, we investigate the involvement of LD lipids in these immune responses. Comparative shotgun and targeted lipidomics demonstrate that *in vivo* host LDs accumulate polyunsaturated fatty acids (PUFAs). PUFAs arrive at cells from the bloodstream to be further metabolised into complex PUFAs accrued by LD-triglycerides and -phospholipids. Host lipid metabolism is transcriptionally controlled by rapid, transient, and intricate immune programs initiated by pathogen-associated molecular patterns and relayed by cytokines such as interferons (type I and II), interleukins (IL-1β), and tumour necrosis factor. When this lipid and signalling environment is reproduced in cultured macrophages, newly formed LDs accumulate defensive proteins, coordinate the synthesis of complex PUFAs, and become PUFA reservoirs and suppliers. Among LD-PUFAs, the ω-6 arachidonic acid is the most actively metabolised during the initial phases of innate immunity. Released from LDs by adipose triglyceride lipase, arachidonic acid is used by macrophages for prostaglandin synthesis, bacterial phagocytosis, and elimination of microbes.

**Graphical Abstract:** 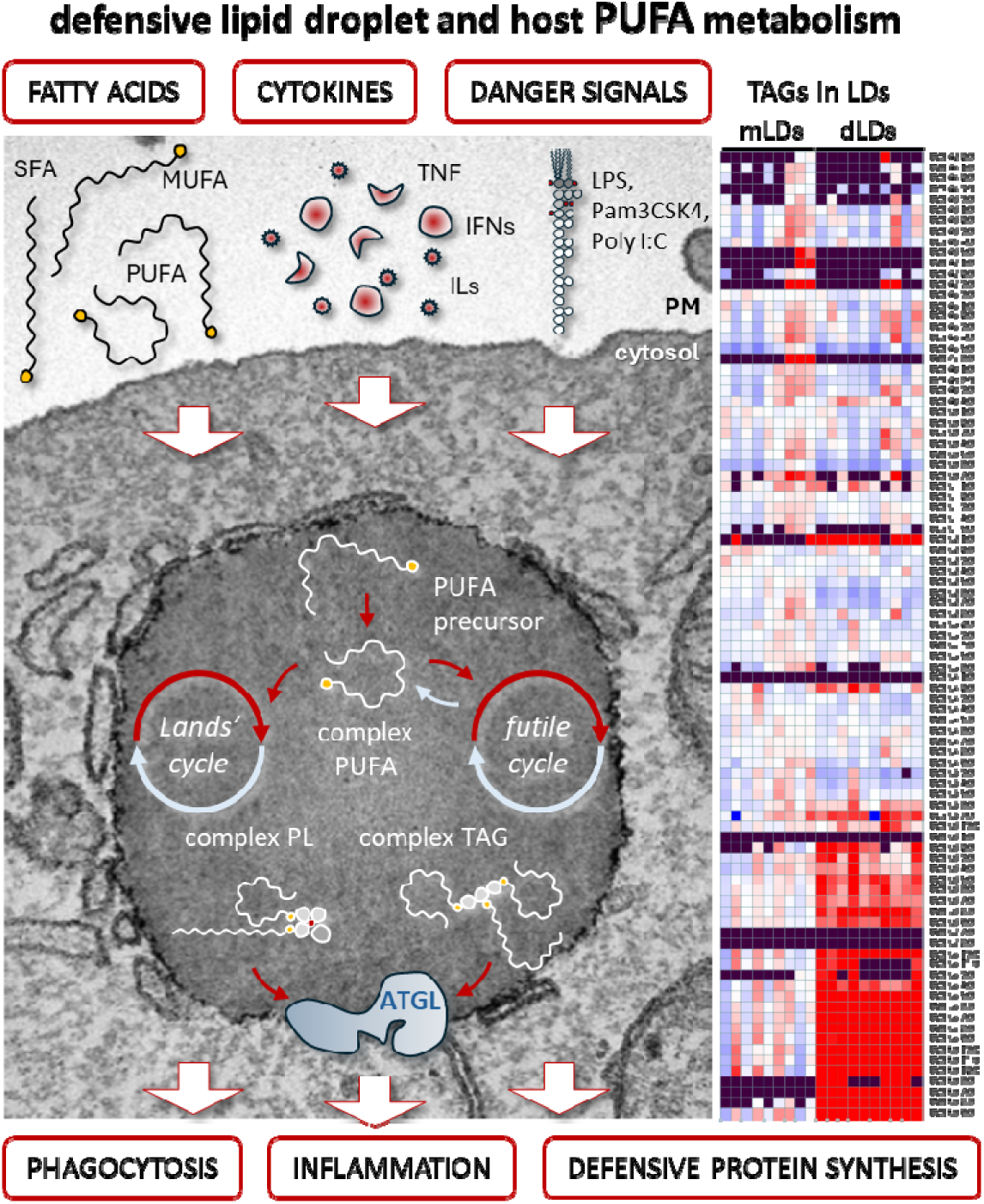

## Introduction

In all eukaryotic cells, lipids are stored and distributed by lipid droplets (LDs) ^1^. Rapidly formed when extracellular nutrients are available, LDs are highly dynamic organelles that sense and adapt to the cellular needs by progressively switching from efficient lipid depots to active lipid suppliers. On demand, LDs provide cells with metabolic energy in the form of fatty acids (FAs) and with substrates to synthesise membranes (e.g. cholesterol), lipid-based compounds (e.g. eicosanoids), and influential signalling molecules (e.g. ceramides) ^2^. Furthermore, reflecting additional roles beyond lipid metabolism, the LD proteome includes proteins not directly related to lipids such as histones, toxic compounds, transcription factors, and immune-related proteins ^3^.

Intriguingly, infected cells rapidly accumulate LDs. In recent years, the list of pathogens known to induce and physically interact with LDs has grown exponentially to include some of the most medically relevant pathogens, including bacteria (e.g. *Mycobacterium tuberculosis*), viruses (e.g. hepatitis C), and parasites (e.g. *Plasmodium falciparum*) ^4^. The widespread explanation for fat-laden infected cells is that pathogens manipulate host lipid metabolism to obtain substrates for growth or long-term survival ^5–8^.

While considered for a long time to be passive lipid inclusions, three decades of intensive research have positioned LDs as key hubs of cell metabolism but also as generic responders to a variety of cellular stresses ^9,10^. Among organelles, LDs can be promptly formed and deeply remodel their proteome to provide cells with transient platforms for organising urgent feedback to stress situations. To survive in the face of an endless number of aggressive invaders, host cells have efficiently coevolved with pathogens by developing a plethora of alarm mechanisms and generic defensive reactions collectively defined as innate immunity. Dependence on host lipids could be a generic weakness of invaders, with LDs playing a crucial role as a strategic location for sensing infection and rapidly organising several innate immunity responses ^11–13^.

So far, the innate immune functions attributed to host LDs encompass microbial killing, immune signalling, and lipid-mediated inflammation ^14^. The proteomic characterisation of hepatic LDs purified from mice treated with lipopolysaccharide (LPS), a potent activator of innate immunity, demonstrated that more than 300 proteins are recruited to LDs in response to danger signals ^15^. LPS-LDs collect antiviral proteins such as viperin (Rsad2), antimicrobial peptides such as cathelicidin (Camp), and a plethora of interferon (IFN)-inducible GTPases such as Igtp, Iigp1, Tgtp1, and Ifi47 ^15–17^. Furthermore, LDs use histones to protect bacterially infected *Drosophila* embryos ^18^. Thus, when compared to the LDs formed in healthy cells, host LDs have a unique proteome. Hereafter, we use the terms “defensive-LDs” (dLDs) and “metabolic-LDs” (mLDs) to emphasise these traits ^4^.

Although mechanistic details are unknown, dLDs mediate the synthesis of IFN in virally infected hepatocytes and pancreatic beta cells ^19–21^ or in *E. coli*-infected macrophages and mice models of sepsis ^22^. Similarly, the synthesis of interleukin-6 (IL-6) is reduced when the assembly of dLDs is impaired, or the activity of the LD-resident and rate-limiting adipose triglyceride lipase (ATGL) is inhibited ^22–24^. Recently, the presence of viral sensors (e.g. Rig-I and Mda5) and downstream transcription factors (e.g. STAT1 and STAT2) on LDs has been proposed ^25^, potentially explaining the role of dLDs in the transcriptional modulation of immune proteins.

Host LDs have also been involved in lipid-mediated inflammation. Early studies proposed that production of prostaglandins, such as PGE2, occurs locally on LDs ^26,27^; a process involving LD polyunsaturated fatty acids (PUFAs), such as arachidonic acid (ARA), and the activities of phospholipase A2 (PLA2) ^28^ and cyclooxygenase-2 (COX-2) ^29^. More recently, several studies have demonstrated that ATGL provides ARA for the synthesis of PGE2 in cell cultures and *in vivo* models ^22–24,30,31^ and thus, positioning LDs as pro-inflammatory organelles.

Therefore, while many mechanistic details are still missing, several studies point towards dLDs occupying a strategic place in the immune responses simultaneously occurring within host cells. This raises several fundamental biological questions. How does innate immunity induce biogenesis of dLDs or remodels “metabolic” LDs into “defensive” LDs? Are there equivalent changes in the dLD lipidome as occurs with their proteome? And if so, what is the role of dLD lipids in antimicrobial defence? Indeed, although much less characterised than the LD proteome ^32^, we anticipate major changes in the dLD lipidome and predict a key role for lipids in innate immunity.

Here, we have performed a comprehensive analysis of the lipid component of dLDs purified from LPS-treated mouse livers. The high purity of these fractions allowed in previous studies the detection of significant changes even in minor dLD proteins ^15^. Now, comparative shotgun and targeted lipidomics provide unprecedented details of the *in vivo* composition of both mLDs and dLDs and identify LPS-induced changes in minor but highly bioactive lipid species. In parallel, we have systematically analysed the immune signalling pathways driving accumulation and metabolism of dLD lipids.

The results demonstrate that dLDs are generic immune responders that, formed by an intense crosstalk between the transcriptional programs of innate immunity and lipid metabolism, use proteins and lipids to participate in a plethora of antimicrobial responses.

## Results

### Lipidomic characterisation of defensive LDs

Lipid droplets (LDs) rapidly form in infected cells, reflecting a link between immunity and lipid metabolism ^4^. Innate immune triggered accumulation of intracellular lipids occurs in both immune “professional” and “non-professional” cells, likely reflecting that dLDs organise generic defences. For instance, dLDs accumulate in THP-1 macrophages infected with *Escherichia coli* (Figs. 1A, 1B, and 1C) but also in hepatocytes of mice treated with LPS (Figs. 1D, 1E, and 1F). When compared with the LDs of untreated or starving cells, the LDs formed during infection (Fig. 1C) or after LPS (Fig.1F) were significantly smaller, suggesting a distinct lipid composition.

**Figure 1.**
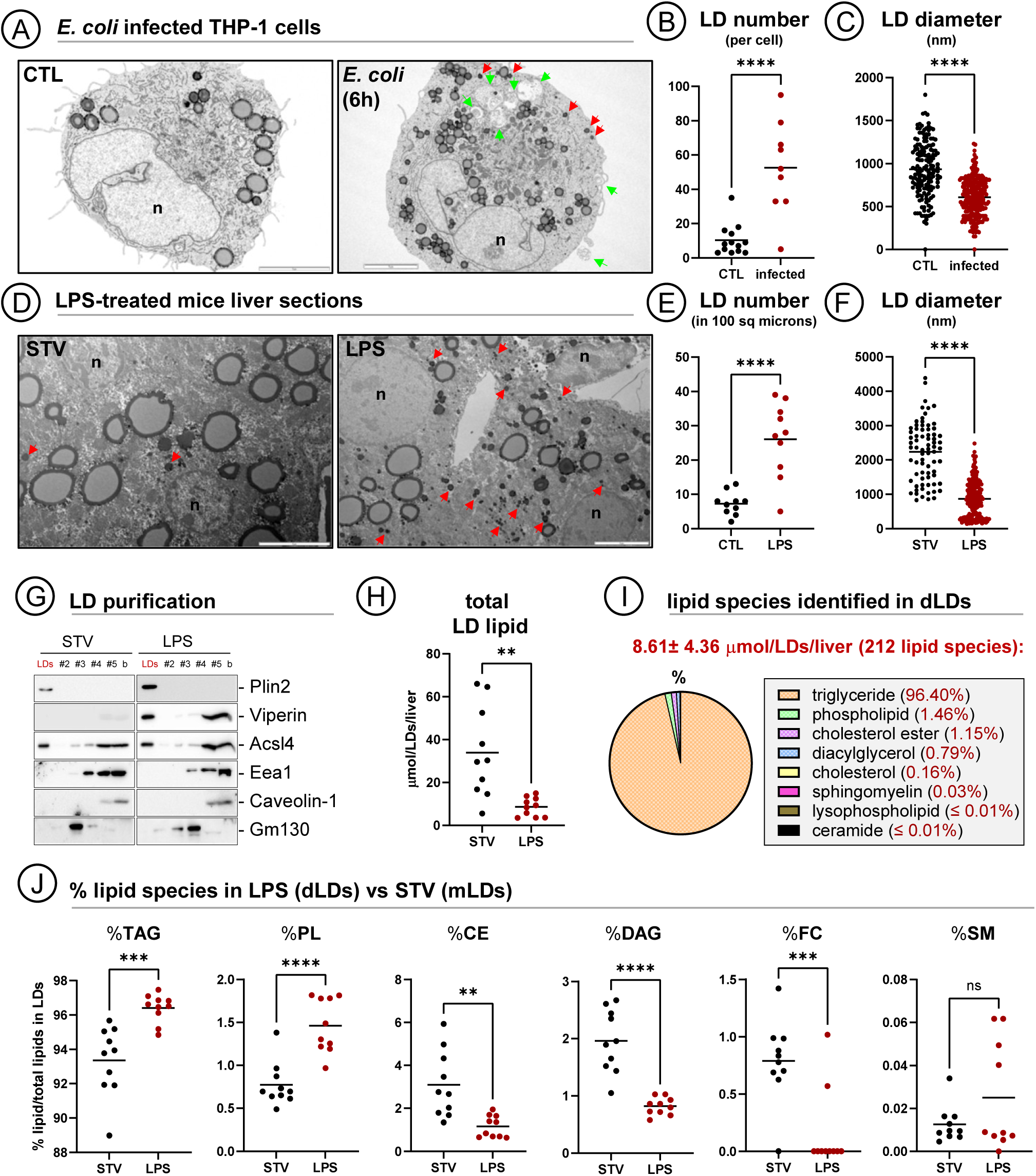
interplay between innate immunity and lipid metabolism. (**A to C**) Representative transmission electron microscopy images of THP-1 macrophages uninfected (CTL) or infected with *E. coli* for 6 hours (A). Red arrows indicate LDs and green arrows bacteria. The number of LDs per cell was counted in 24 CTL- and 24 infected-cells (B) and the apparent diameter of 161 CTL- and 255 infected-cell LDs measured (C). Graphs show means (combined from N=2); ****P < 0.0001 calculated in a two-tailed unpaired t-test. Scale bars are 5μm. **(D to F)** Representative transmission electron microscopy images of liver sections from 16 hours starved mice (STV, mLDs) or mice starved and additionally treated with LPS (LPS, dLDs). Red arrows indicate LDs. The number of LDs per cell was counted in 20 CTL- and 20 LPS-treated hepatocytes (E) and the apparent diameter of 73 STV- and 179 LPS-LDs measured (F). Graphs show means (combined from N=2); ****P < 0.0001 calculated in a two-tailed unpaired t-test. Scale bars are 5μm. **(G)** Livers from mice starved for 16 hours (STV) or from mice additionally treated with LPS (LPS) were fractionated in sucrose density gradients. Distribution of organelle and LD markers (Eea1, endosomes; Caveolin-1, plasma membrane; GM130, Golgi) was analysed by Western blotting (representative of N=2). LD core proteins such as Plin2 or Acsl4 and immune LD proteins such as Viperin floated onto the top fraction (LDs). **(H)** Shotgun lipidomic analysis of the LD fractions purified in (Fig. 1H). The graph shows the mean (μmols) of total lipid quantified in LDs from each individual liver (combined from N=10); **P < 0.01 calculated in a two-tailed unpaired t-test. Raw data in Table S1. **(I)** The dLD fraction (purified from LPS-livers) of each liver contained an average of 8.61 μmol of lipids distributed among 212 different lipid species (see complete description in figures below). The pie chart depicts the average percentage (%) of each lipid species within the total lipid in dLDs (mean of N=10). **(J)** Comparison of the relative abundance in dLDs (purified from LPS-livers) or mLDs (purified from STV-livers) of the lipid species identified in (Fig. 1J). Graphs show means (combined from N=10); ns is not significant, *P < 0.05, **P < 0.01, ***P < 0.001, ****P < 0.0001 in a two-tailed unpaired t-test. Abbreviations: triacylglycerol, TAG; phospholipids, PL; cholesterol esters, CE; diacylglycerol, DAG; free cholesterol, FC; and sphingomyelin, SM.

To determine their lipid composition, mLDs (16 hours starvation) and dLDs (starvation+LPS for 16 hours) were purified from mouse livers (Fig. 1G). Validating our approach, viperin was enriched in dLDs but was absent from mLDs (Fig. 1G). Next, lipid species in purified LDs were identified and quantified by shotgun lipidomics. The total amount of lipid was significantly lower in dLDs than in mLDs (Fig. 1H). The analysis identified 263 different lipid species (Tables S1 and S2). Among them, triacylglycerol (TAG) represented 96% of dLD lipids, followed by phospholipids (PL, 1.5%), cholesterol esters (CE, 1.2%), diacylglycerol (DAG, 0.8%), free cholesterol (FC, 0.2%), and only traces of sphingomyelin (SM), lysophospholipids, and ceramides (less than 0.1%) (Fig. 1I) (Table S1). When compared with mLDs (Fig. 1J), dLDs were enriched in TAGs and PLs but contained less DAGs and very low levels of FCs or CEs. The low levels of CE and FC may reflect the inhibition of cholesterol synthesis in infected cells ^33^.

Thus, the most distinctive trait of dLDs is the relative enrichment of PLs which is consistent with the smaller size of dLDs and likely reflects a larger surface area to volume ratio. To evaluate if dLDs potentially represents a relevant cellular pool of PLs, we estimated the number of dLD-PLs. Quantitative stereological analysis from random electron microscopy (EM) sections of the hepatocytes of LPS treated mice was used to provide an estimate of dLD surface area. In the sampled cells, the total surface area of the LDs per hepatocyte was approximately 2.59x higher than the plasma membrane. By relating the relative LD surface area on random sections to the absolute volume of the nucleus, as measured by light microscopy (mean diameter 9.8µm), an estimate of the total PL surface of dLDs per cell was obtained. Assuming a FA packing density of 1.35 PLs per nm^2^ ^34^, this suggested 3.65×10^9^ total dLD-PLs in one hepatocyte of LPS-treated liver (cf 1.94×10^9^ in starved liver). This estimation illustrates both the magnitude of the lipid changes occurring in LPS-treated cells and the ability of dLDs to control cellular PL balance.

### Defensive LDs accumulate and supply polyunsaturated fatty acids

Next, we analysed the FA species of dLDs by targeted lipidomics (LC/ESI-MS/MS). The analysis identified 26 different species (Fig. 2A) (Table S3). Among FAs, the ω-6 PUFA linoleic acid (LA) was the most abundant at 30.96%, followed by the monounsaturated (MUFA) oleic acid (OA, 26.87%), and the saturated (SFA) palmitic acid (PA, 17.82%) (Fig. 2B).

**Figure 2.**
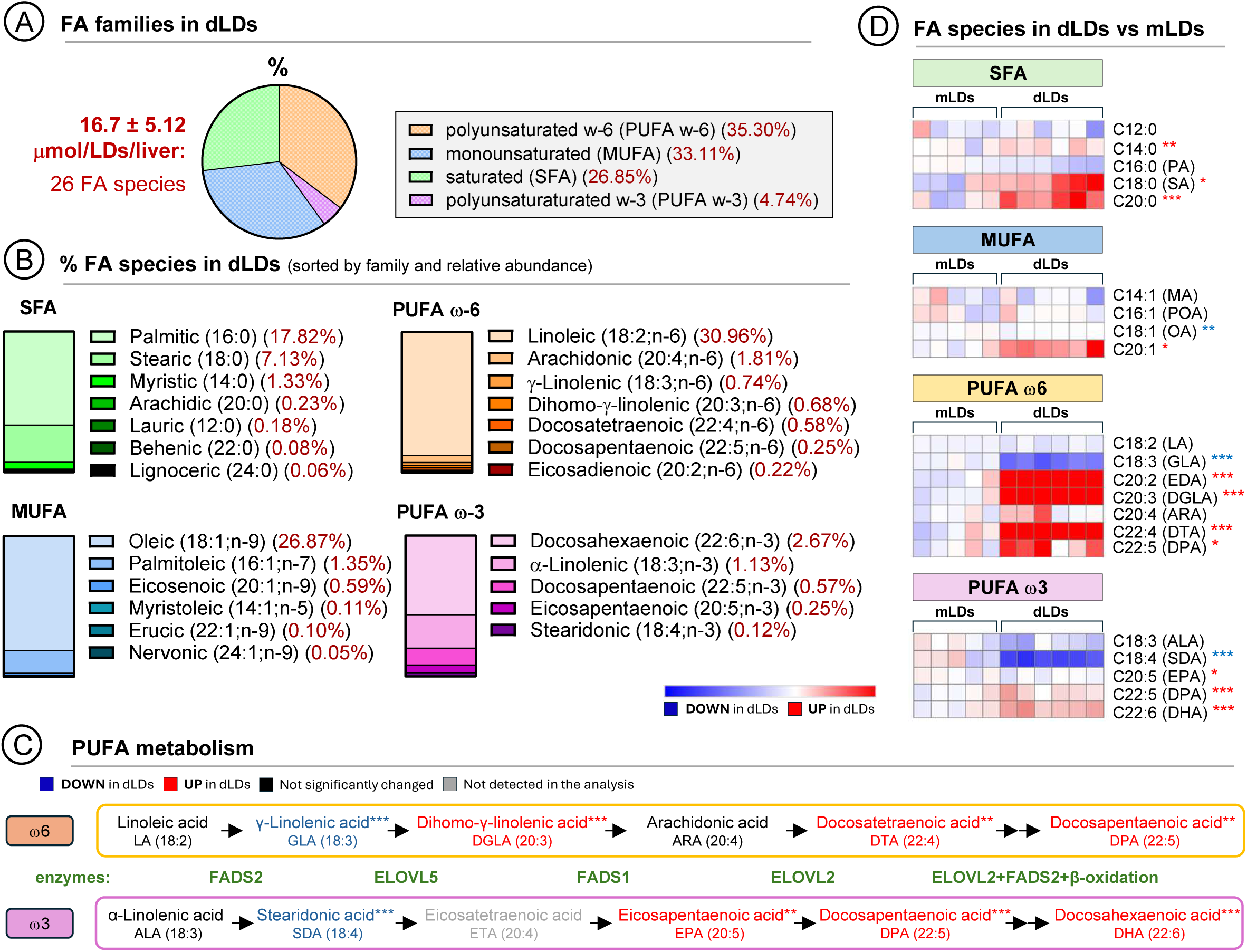
Fatty acid species identified in lipid droplets. **(A)** LDs were purified as in (Fig. 1H) and FAs identified by targeted lipidomics (LC/ESI-MS/MS). The dLDs purified from each individual liver contained an average of 16.7 μmol of FAs distributed among 26 different FA species. The box shows the relative (%) abundance of each FA family (combined from N=6). Raw data in Table S3. **(B)** Relative abundance of each FA when compared to the total amount of dLD FAs (combined from N=6). Raw data in Table S3. **(C)** Schematic representation of PUFA metabolism. Precursor ω-6 (orange box) and ω-3 (purple box) PUFAs are derived from dietary sources to be metabolised into complex PUFAs by the action of elongases and desaturases (green letters). PUFAs significantly enriched or reduced in dLDs when compared with mLDs are respectively indicated with red or blue letters. **(D)** Heatmaps showing the relative abundance (%) of each FA in dLDs compared to mLDs, as normalised to the mLD average for each subspecies (A). Red and blue asterisks indicate respectively that the FA is significantly enriched or reduced in dLD: **P < 0.01, ***P < 0.001 in a two-tailed unpaired t-test. Raw data in Table S3.

As far as we know, this is the first analysis that determines that LA is the major FA of LDs *in vivo* ^32^, an enrichment confirmed by shotgun lipidomics (Table S2). LA is an essential FA obtained through dietary sources and is the precursor of the rest of ω-6 PUFAs (Fig. 2C). Both mLDs and dLDs also contained the ω-3 PUFA α-linolenic acid (ALA, 1.13%), essential FA and the precursor of the rest of ω-3 PUFAs (Fig. 2C). When compared with mLDs, dLDs were significantly enriched in complex ω-6 and ω-3 PUFA (20 and 22 carbons) but depleted in precursors (18 carbons), reflecting an active PUFA metabolism in response to LPS (Fig. 2D). The only complex PUFA that was not significantly accumulated in dLDs was ARA (ω-6; 20:4), suggesting that ARA may be actively consumed in the LDs of LPS-activated cells.

The apparent enrichment in PUFAs suggests that dLDs may be formed from exogenous FAs. To examine this possibility, we analysed the FA composition of mouse serum after the LPS treatment (2 hours). The serum of LPS-treated animals was significantly enriched in FAs when compared with serum from control mice (Fig. 3A and Table S4). The ω-6 PUFAs were the most abundant FAs (40.34%), followed by SFAs (29.82%), MUFAs (24.62%), and ω-3 PUFAs (5.22%). A total of 33 different FA species were identified, with LA being the most abundant (27.30%), PA the most common SFA (21.75%) and OA the major MUFA (18.75%) (Fig. 3B), explaining the enrichment of PUFAs in dLDs. When compared with the serum of control mice, the LPS-serum was enriched in most FAs but especially in some minor ω-6 and ω-3 PUFAs such as dihomo-γ-linolenic acid (DGLA, ω-6; 20:3) and docosapentaenoic acid (DPA, ω-3; 22:5) (Fig. 3C). Indeed, the relative FA composition of dLDs largely reflects that of LPS-serum with the striking exception of ARA (Fig. 3D), suggesting again an active consumption of this FA specifically at the LDs of LPS-activated cells. Differences in the PA/OA ratio likely reflect a differential affinity of diacylglycerol acyltransferase (DGAT) enzymes for esterification of these FAs into TAGs ^35^.

**Figure 3.**
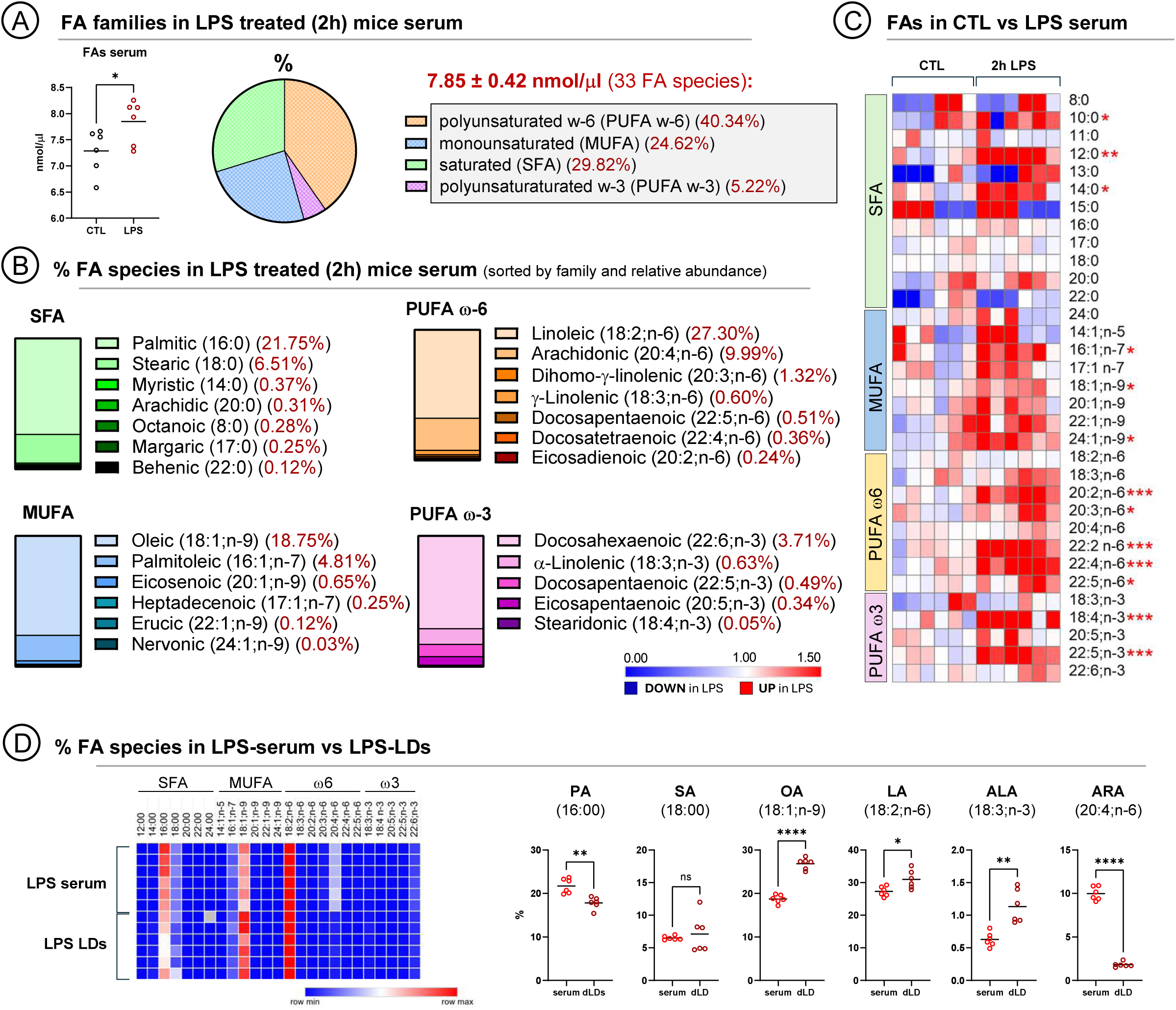
Fatty acid species identified in mice serum. **(A)** Serum was obtained from untreated (CTL) or from mice 2 hours after LPS administration (LPS) and FAs identified by targeted lipidomics (LC/ESI-MS/MS). The average amount of FA/μl in CTL- and LPS-serum is shown in the graph (combined from N=6); *P < 0.05 in an unpaired t-test. Each μl of LPS-serum contained an average of 7.85 nmols of FAs distributed among 33 different species. The pie chart depicts the percentage (%) of each FA family with respect to the total FAs found in LPS-serum (combined from N=6). Raw data in Table S4. **(B)** Relative abundance of each FA when compared to the total FAs in LPS-serum (average of N=6). Raw data in Table S4. **(C)** Heatmaps comparing the concentration (nmols/μl) for each FA species in CTL-versus LPS-serum, as normalised to control average. Red asterisks indicate that the FA is statistically enriched in LPS-serum (combined from N=6); *P < 0.05, **P < 0.01, ***P < 0.001 in a two-tailed unpaired t-test. Raw data in Table S3. **(D)** Heatmap comparing the relative abundance (%) of each FA in LPS-serum (Fig. 3B) and dLDs (Fig. 2B). Most abundant FAs are compared in the graphs. Red and blue asterisks indicate respectively that the FA is significantly enriched or reduced in dLD. Graphs show means (combined from N=6); ns is not significant, *P < 0.05, **P < 0.01, ***P < 0.001, ****P < 0.0001 in an unpaired t-test.

We next determined the dLD lipid species that accumulate and supply PUFAs. The shotgun lipidomics analysis of dLD TAG identified 92 different species (Fig. 4A). The most abundant TAGs contained between 50 and 56 carbons (Fig. 4B). When compared with mLDs, dLDs were significantly enriched in very long (from 56 carbons upwards) and largely unsaturated (up to 12 double bonds) TAGs (Fig. 4C), supporting the notion that dLD TAGs are PUFA reservoirs.

**Figure 4.**
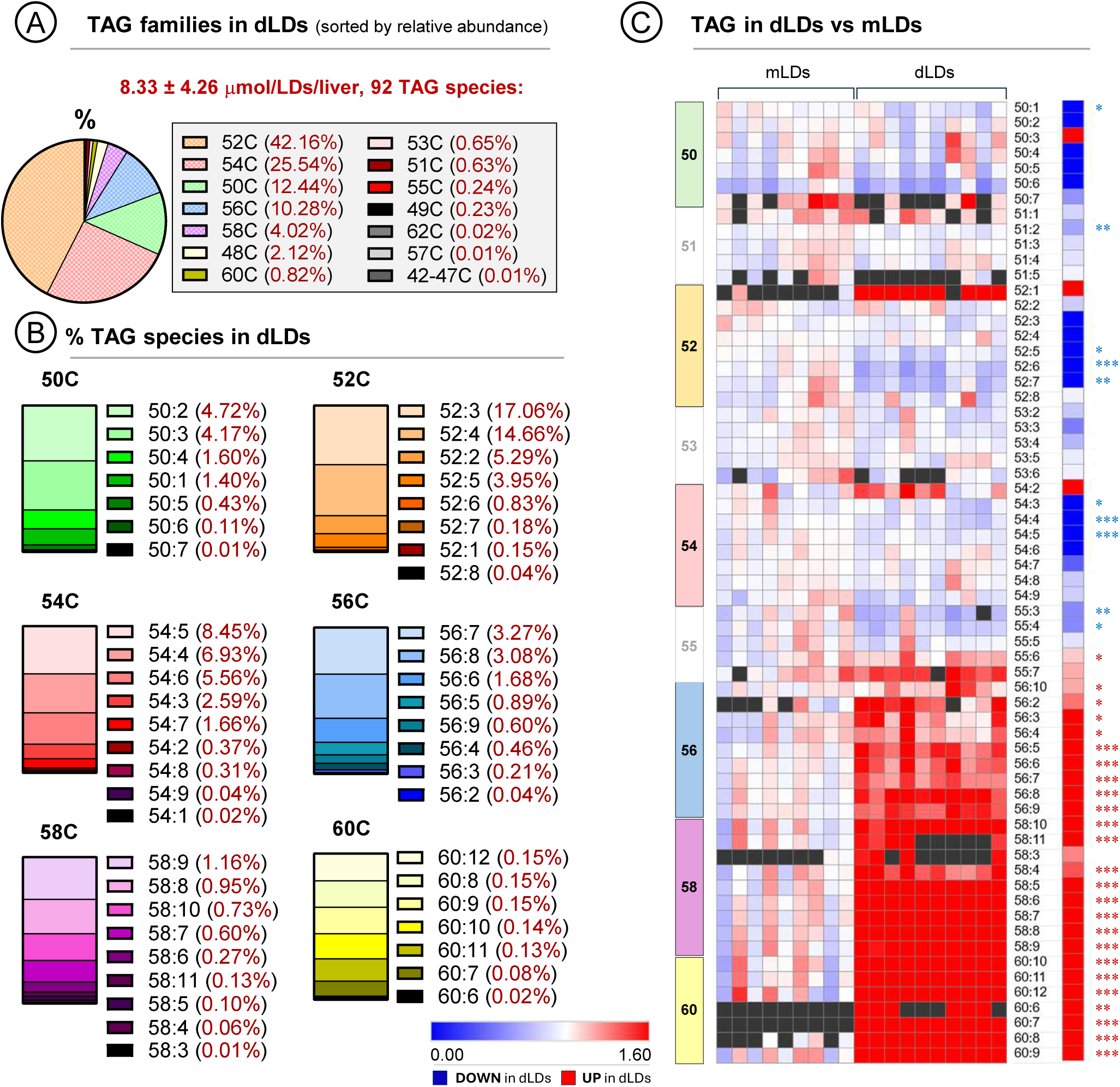
Triglyceride species identified in lipid droplets. **(A)** LDs were purified as in (Fig. 1H) and TAGs identified by shotgun lipidomics. The dLD fraction of each liver contained an average of 8.3 μmol of TAGs distributed among 92 different species (N=10). The box shows the relative abundance of major TAGs classified accordingly the number of carbons. Raw data in Table S1. **(B)** Relative abundance (%) of each TAG species with respect to the total TAGs found in dLDs and classified accordingly to the number of carbons and number of double bounds (combined from N=10). Raw data in Table S1. **(C)** Heatmaps showing the relative abundance (%) of each TAG species in dLDs as compared to mLDs, as normalised to the mLD average of each corresponding species. Red and blue asterisks indicate respectively that the TAG is significantly enriched or reduced in dLDs (combined from N=10); *P < 0.05, **P < 0.01, ***P < 0.001, ****P < 0.0001 in an unpaired t-test. Raw data in Table S1. The additional heatmap column represents the differential value from substracting the averages of each species from each class.

Although only representing 1.5 mol % of the total lipid (Fig. 1I), the lipidomic analysis detected significant changes in LD-PLs. A total of 43 different species of PLs were identified in dLDs (Fig. 5A) (34 species in mLDs), with phosphatidylcholine being the most abundant (PC, 56.67%), followed by phosphatidylethanolamine (PE, 27.78%), phosphatidylinositol (PI, 14.94%), and only traces of phosphatidylserine (PS, 0.24%) (Fig. 5A). The dLD PLs generally contained PA (16:0) or stearic acid (SA, 18:0) and one PUFA, being PC 16:0_18:2 (17.84%) and PI 18:0_20:4 (12.86%) the most abundant (Fig. 5B). When compared with mLDs, dLDs were slightly but significantly enriched in PC but depleted of PE (Fig. S1A). The analysis also detected an enrichment of SA but a reduction of PA and OA in sn-1 position (Fig. S1B), perhaps partially explaining the reduced size of dLDs ^36^.

**Figure 5.**
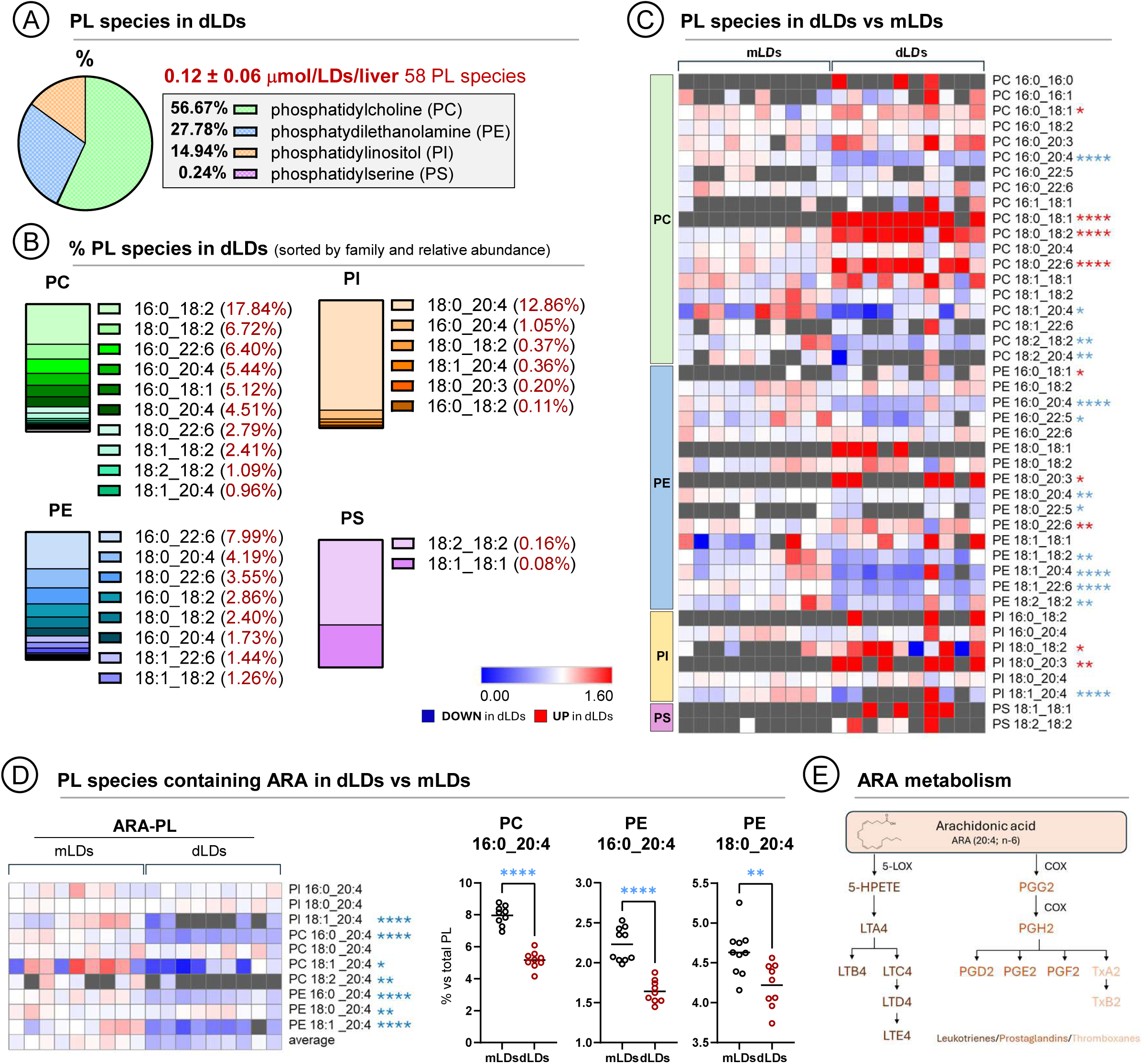
Phospholipid species identified in lipid droplets. **(A)** LDs were purified as in (Fig. 1H) and PLs identified by shotgun lipidomics. The dLD purified fraction of each liver contained an average of 0.12 μmol of PLs distributed among 58 different species. The box shows the relative abundance of each PL species (combined from N=10). Raw data in Table S1, only PLs identified in at least n=3 were included in the analysis. **(B)** Relative abundance of each PL with respect to the total amount of dLD-PLs. The FA species contained in each PL are indicated (combined from N=10). Raw data in Table S1. **(C)** Heatmaps showing the relative abundance (%) of each PL species in dLDs compared to mLDs, as normalised to mLD average. Red and blue asterisks respectively indicate that the PL is significantly enriched or reduced in dLD. Combined from N=10; *P < 0.05, **P < 0.01, ***P < 0.001, ****P < 0.0001 in an unpaired t-test. Raw data in Table S1 (N=10). **(D)** Heatmaps comparing the relative abundance (%) of ARA-containing PLs across mLDs and dLDs samples, as normalised to mLD average. Blue asterisks indicate that the PL is significantly reduced in dLDs. The graphs compare the abundance (%) of major PC and PE species containing ARA. Combined from N=9; *P < 0.05, **P < 0.01, ***P < 0.001, ****P < 0.0001 in an unpaired t-test. Raw data in Table S1. **(E)** Schematic representation of ARA metabolism and its involvement in inflammatory eicosanoid formation.

Interestingly, the dLD PLs were significantly depleted of ARA (Fig. 5D), pinpointing that PLs are the ARA donors previously anticipated (Fig. 3). ARA reduction was especially significant in abundant PLs such as PC 16:0_20:4, PE 16:0_20:4, and PE 18:0_20:4, representing 11.4 mol % of the total PL in dLDs (Fig. 5D). Further, the reduction was specific for ARA and not observed for other bioactive FAs such as docosahexaenoic acid (DHA, ω3; 22:6) (Fig 5C). The depletion of ARA occurs in PC and PE but not in other abundant PLs such as PI 18:0_20:4 (12.86 mol % total PL) or lipid species such as CEs or DAGs (Fig. S2), clearly suggesting that dLD PC and PE are ARA donors. In conclusion, PUFA metabolism is activated in response to LPS and dLDs accumulate and supply these PUFAs. Due to the large surface area occupied by LD-PLs this role could have important implications in the immune response, especially in inflammation since ARA is metabolised into inflammatory eicosanoids (Fig. 5E).

### Innate immune signals driving dLD formation and consumption

Thus, the dLDs formed by innate immunity have a characteristic lipid composition. To infer connections between immune signals and lipid metabolism, we used Ingenuity Pathway Analysis (IPA) to model networks of transcriptional upstream regulators of proteins enriched in dLD when compared to mLDs ^15^. The *in-silico* analysis predicts that dLDs are generated in response to self-reinforcing circuits simultaneously activated by immunity and metabolism programs (Fig. 6A). Immune signals triggered by pathogen associated molecular patterns (PAMPs) and Toll-like receptors (TLR), would be translated into transcription factors (NFκB, IRF or STAT) and transduced by cytokines (TNF, IL1-1β, and IFNs). Both, the dLD “core proteome” (exemplified by Plin2, ATGL, and ACSL4) and the dLD “immune proteome” (exemplified by viperin, Igtp, and Camp) could be simultaneously regulated by key transcription factors of innate immunity (NFκB, IRF or STAT) and lipid metabolism (PPARs and SREBFs). The best example of this intense crosstalk is Plin2, the major dLD protein, which emerges as a key hub integrating multiple immunity and metabolism signals.

**Figure 6.**
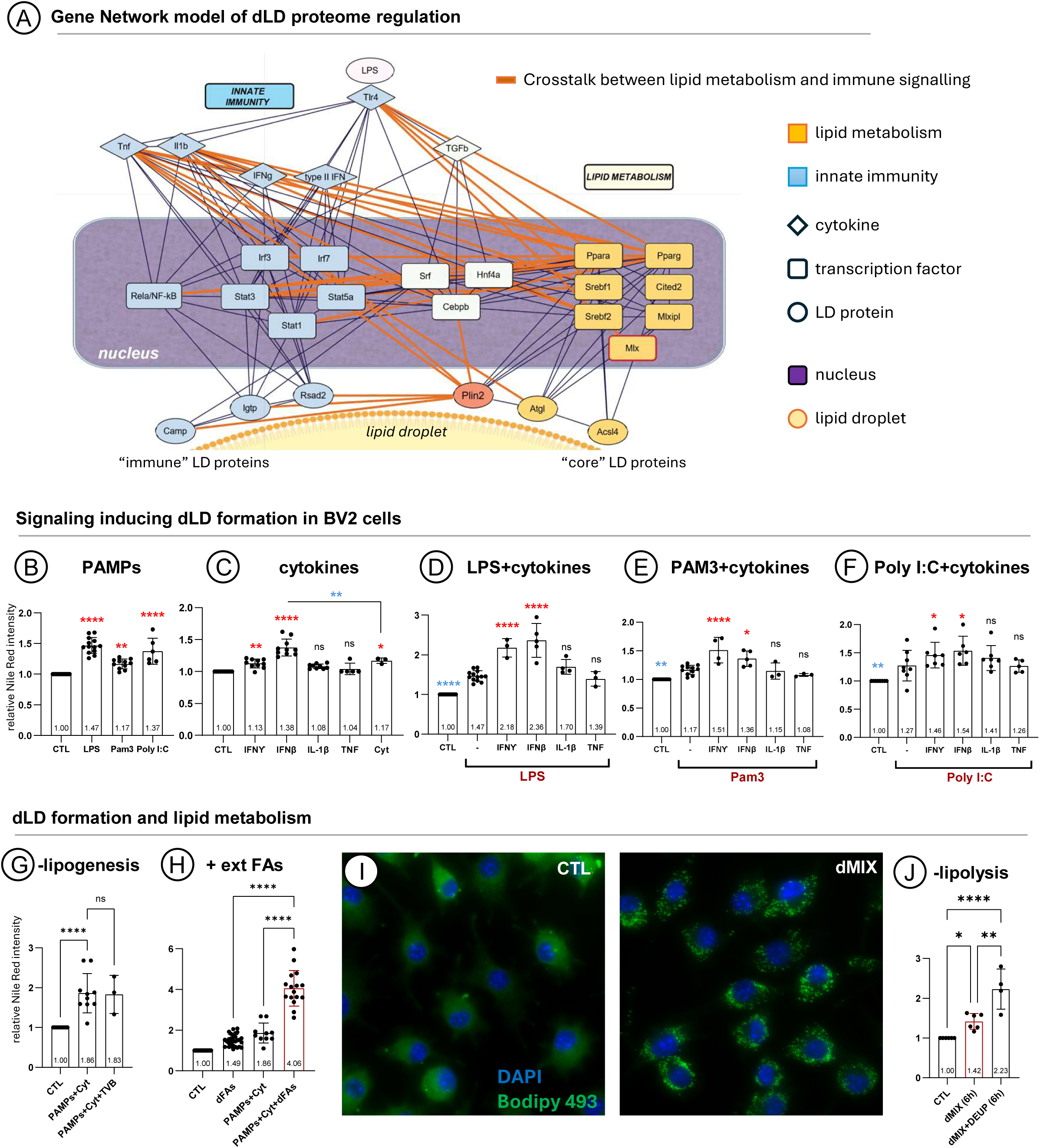
Innate immunity signals regulating defensive lipid droplets. **(A)** Gene network modeling of regulators governing proteins up-regulated in LPS-LDs ^15^ by applying Upstream Regulation Inference analyses from the Ingenuity Pathway Analysis (IPA™) platform; gene relationships are completed from STRING™ database. Cytokines and TLRs are indicated with rhombuses, transcription factors with squares, and proteins with circles. The blue color indicates defensive programs, and the yellow denotes lipid metabolism. Orange lines highlight crosstalk relationships between inflammation and lipid metabolism nodes. **(B to F)** The LD content of BV2 cells was determined by flow cytometry after 16 hours of a treatment with PAMPs, cytokines, or both simultaneously. (B) shows the LD content after LPS, Pam3CSK4, or Poly I:C. (C) shows the LD content in cells treated with IFNγ, IFNβ, IL-1β, TNF, or the combination of the four cytokines (Cyt). (D to F) show the LD content in cells co-treated with the indicated PAMP and cytokine. Graphs show means±SD (combined from at least N=3); ns is not significant, *P < 0.05, **P < 0.01, ***P < 0.001, ****P < 0.0001 in an unpaired t-test vs untreated cells (CTL) for B and C or vs PAMP-treated cells for D, E, and F. Red and blue asterisks indicate respectively significant enrichment or reduction. **(G)** The LD content of BV2 cells untreated (CTL) or treated for 16 hours with a combination of PAMPs (LPS and Pam3CSK4) and cytokines (Cyt: IFNγ, IFNβ, IL-1β, and TNF) was determined in the presence or absence of the lipogenesis inhibitor TVB-2640 (FASN inhibitor). Graph shows means±SD; ns is not significant; ****P < 0.0001 in an ordinary one-way ANOVA. Combined from N=10 for PAMPs+Cyt4 and N=3 for PAMPs+Cyt4+TVB. **(H)** BV2 cells were treated for 16 hours with dFAs (15μm/ml; 40% LA, 35% OA, 23% PA, and 2% ALA), PAMPs and cytokines as in (Fig. 6G), or simultaneously with dFAs, PAMPs, and cytokines (dMIX) and the LD content quantified by flow cytometry. Graph shows means±SD (combined from at least N=10); ns, not significant, ****P < 0.0001 in an ordinary one-way ANOVA. **(I)** Representative confocal images (N=3) of BV2 cells untreated (CTL) or treated with the dMIX for 16 hours and labelled with BODIPY (LDs, green) and DAPI (nucleus, blue). Scale bar is 20µm. **(J)** The LD content of BV2 cells untreated (CTL) or treated for 6 hours with the dMIX was quantified in the presence or absence of DEUP (lipolysis inhibitor). Graph shows means±SD (combined from at least N=4); *P < 0.05, **P < 0.01, ****P < 0.0001 in an ordinary one-way ANOVA.

To experimentally address these predictions, we systematically tested LD formation in cells treated with these factors individually or in combination. For these initial experiments, we selected BV2 cells; a microglial cell line efficiently forming LDs when treated with LPS ^37^. BV2 cells were treated with PAMPs or/and cytokines and LD formation quantified by flow cytometry 16 hours later. Initially, we tested different PAMPs: LPS to activate TLR4, Pam3CSK4 for TLR1/TLR2 and Poly I:C for TLR3. All three PAMPs increased LD accumulation (Fig. 6B), reinforcing the notion that LD formation is a generic immune response. Next, we analysed LD formation in response to cytokines. IFNγ and IFNβ promoted the accumulation of LDs while IL-1β and TNF had little effect (Fig. 6C). When cells were incubated with the four cytokines simultaneously (Fig. 6C), the LD content was lower than in cells only treated with IFN, suggesting that some cytokines promote the net formation of LDs while others may increase their consumption. We next combined PAMPs and cytokines. In all cases, IFNs potentiated the LD formation induced by PAMPs, while IL-1β and TNF had little effect or induced their consumption (Figs. 6D, 6E, and 6F). Thus, multiple innate immune signals coordinate formation and metabolism of dLDs.

According to the lipidomics data (Figs. 2 and 3), dLDs are mostly formed by accumulation of extracellular FAs. To corroborate this hypothesis, BV2 cells were treated with PAMPs (LPS and Pam3CSK4) and cytokines (cyt) to induce LD formation and the *de novo* synthesis of FAs (lipogenesis) was inhibited with the fatty acid synthase (FASN) inhibitor (TVB-2640). The inhibition of lipogenesis did not reduce the formation of LDs (Fig. 6G) and thus, dLD FAs have also an extracellular origin in cultured cells. To reproduce the *in vivo* conditions, BV2 cells were then treated with PAMPs and cytokines and simultaneously incubated with extracellular FAs. We prepared a combination of FAs following the relative proportion of the major FAs found in dLDs: 40% LA, 35% OA, 23% PA, and 2% ALA (hereafter dFAs). The combination of PAMPs, cytokines and dFAs potentiated each other and significantly increased the formation of LDs (Fig. 6H). This cocktail of stimuli, that we propose reproduces the environment of cells in infected individuals, will hereafter be referred to as defensive mix (dMIX). BODIPY staining confirmed the dMIX-induced LD formation in all cells within the culture (Fig. 6I). Lipolysis inhibitors, such as diethylumbelliferyl phosphate (DEUP) ^38^, significantly increased the dMIX-induced accumulation of LD (Fig. 6J), thus demonstrating that dLDs are metabolically active organelles. Further, the dMIX induced dLD formation in all tested mice and human cells, including macrophage-like cell lines BV2 and RAW-264.7, as well as hepatic AML12 and HepG2 cells (Fig. S3A), reflecting that the dMIX generates the predicted generic defence organised by dLDs.

### The defensive MIX generates dLDs in cultured cells

When compared with mLDs, dLDs have a characteristic proteome ^15^ and accumulate/distribute PUFAs (Figs. 2, 4, and 5). Therefore, we next analysed whether the dMIX-induced LDs reproduce these traits. Initially, we characterised the dMIX-induced transcriptional profile in BV2 cells by monitoring two main innate immunity branches: NFκB and IFNs. To do so, we measured expression (by qRT-PCR) of representative down-stream genes such as *Tnf*, *Il-6* and *Ifnβ*. The three cytokines were robustly expressed in dMIX treated cells after 3 hours (red asterisks, Fig. 7A) and progressively downregulated after 16 hours (blue asterisks, Fig. 7A). We expected a similar transcriptional profile of the LD proteome. Indeed, both transcripts encoding LD “core” proteins (represented by *Plin2*, *Ascl4* and *Atgl*) as well as “defensive” LD proteins (represented by *viperin*, *Igtp*, and *cathelicidin*) followed a parallel expression profile to the immune mediators, peaking at 3 hours (red asterisks, Fig.7B) and were gradually down regulated after 16 hours (blue asterisks, Fig.7B). When the intracellular distribution of the defensive LD proteins was analysed by immunofluorescence, they were absent on the LDs of cells treated with OA (commonly used to induce mLDs in cells) but accumulated on dMIX-induced LDs (dLDs) (Igtp in Fig. 7C and viperin in Fig. S4). All the dMIX induced LDs were decorated by Igtp and viperin and thus, innate immunity promotes both formation of new dLDs and conversion of preexisting mLDs into dLDs. We also observed by transmission EM (Fig. S3B) a difference in the size of LDs generated with the dMIX as compared to LDs generated by OA in a HeLa cell model system, consistent with the difference in the size of dLDs vs mLDs observed *in vivo*. Together these data demonstrate that the dMIX generates LDs with the distinct physical (SA:V ratio) and compositional (viperin and Igtp enrichment) properties observed for dLDs *in vivo*.

**Figure 7.**
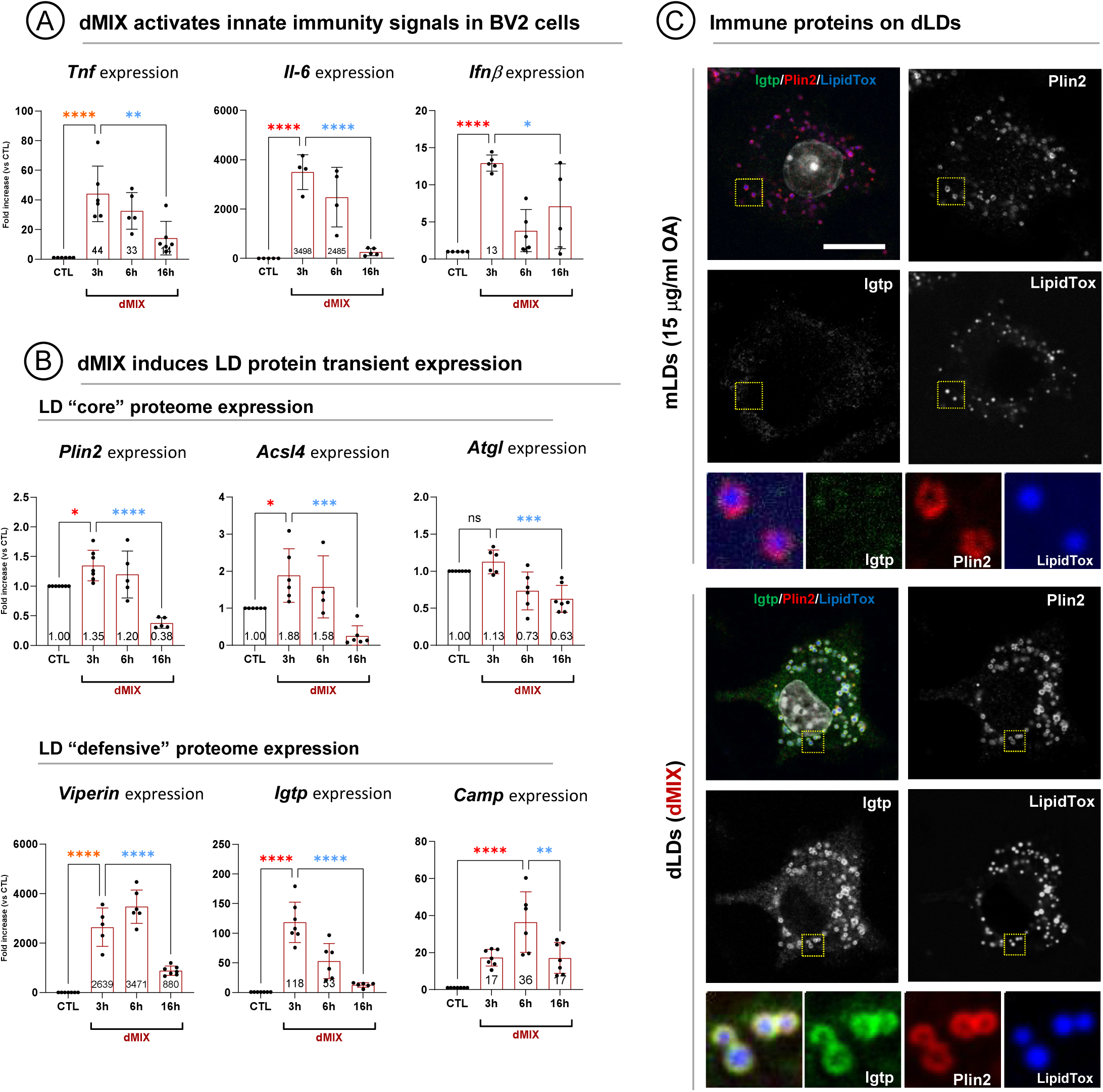
Transcriptional regulation of lipid droplets by innate immunity. **(A)** BV2 cells were activated for 3, 6, or 16 hours with the dMIX and *Tnf*, *Il-6*, and *Ifn*β expression quantified by qRT-PCR and normalised to their expression in untreated cells (CTL). Graphs show means±SD (combined from at least N=4); *P < 0.05, **P < 0.01, ****P < 0.0001 in an ordinary one-way ANOVA. Red and blue asterisks indicate respectively significant enrichment or reduction. **(B)** BV2 cells were stimulated for 3, 6, or 16 hours with dMIX and “core” and “immune” LD proteins expression quantified by qRT-PCR and normalised to their expression in untreated cells (CTL). Graphs show means±SD (combined from at least N=4); ns is not significant, *P < 0.05, **P < 0.01, ****P < 0.0001 in an ordinary one-way ANOVA. Red and blue asterisks indicate respectively significant enrichment or reduction. **(C)** Representative confocal images (N=3) of BV2 cells treated for 16 hours with 15µg/ml OA to induce mLDs or with the dMIX to induce dLDs. Cells were labelled with anti-Igtp (green) and anti-Plin2 (red) antibodies and LipidTox (LD, blue). Scale bar is 20µm. The yellow square marks the region selected for the high magnification panels.

### Defensive LDs regulate cellular PUFA metabolism

The lipidomic analysis of *in vivo* dLDs indicates that host cells activate PUFA metabolism and suggest a key role for dLDs functioning as PUFA depots and suppliers. The dMIX treated cells and the newly formed dLDs are then a convenient cellular model to study this immune response. For these experiments, we utilised RAW-264.7 macrophages, cells with few LDs in control conditions but efficiently forming new dLDs in response to the dMIX (Fig. S3).

We initially characterised the FA composition of RAW-264.7 cells. The macrophages were left untreated or were supplemented with 15 μg/ml of OA or with 15 μg/ml of dFAs. The FA composition of cells across these three conditions was quantified early (6 hours) by targeted lipidomics (LC/ESI-MS/MS). Cells incubated with OA accumulated MUFA (62.50%) but a low concentration of ω-6 PUFAs (7%) (Fig. 8A and Table S5). In contrast, the dFA-treated cells doubled the cellular content of ω-6 PUFAs (15.16%) in just 6 hours (Fig. 8B). Because these cells were only supplied with ω-6 and ω-3 precursors, the accumulation of complex PUFAs demonstrates an active PUFA metabolism by RAW-264.7 macrophages. The enrichment of ARA was also significant and thus, non-activated macrophages do not seem to actively consume this FA.

**Figure 8.**
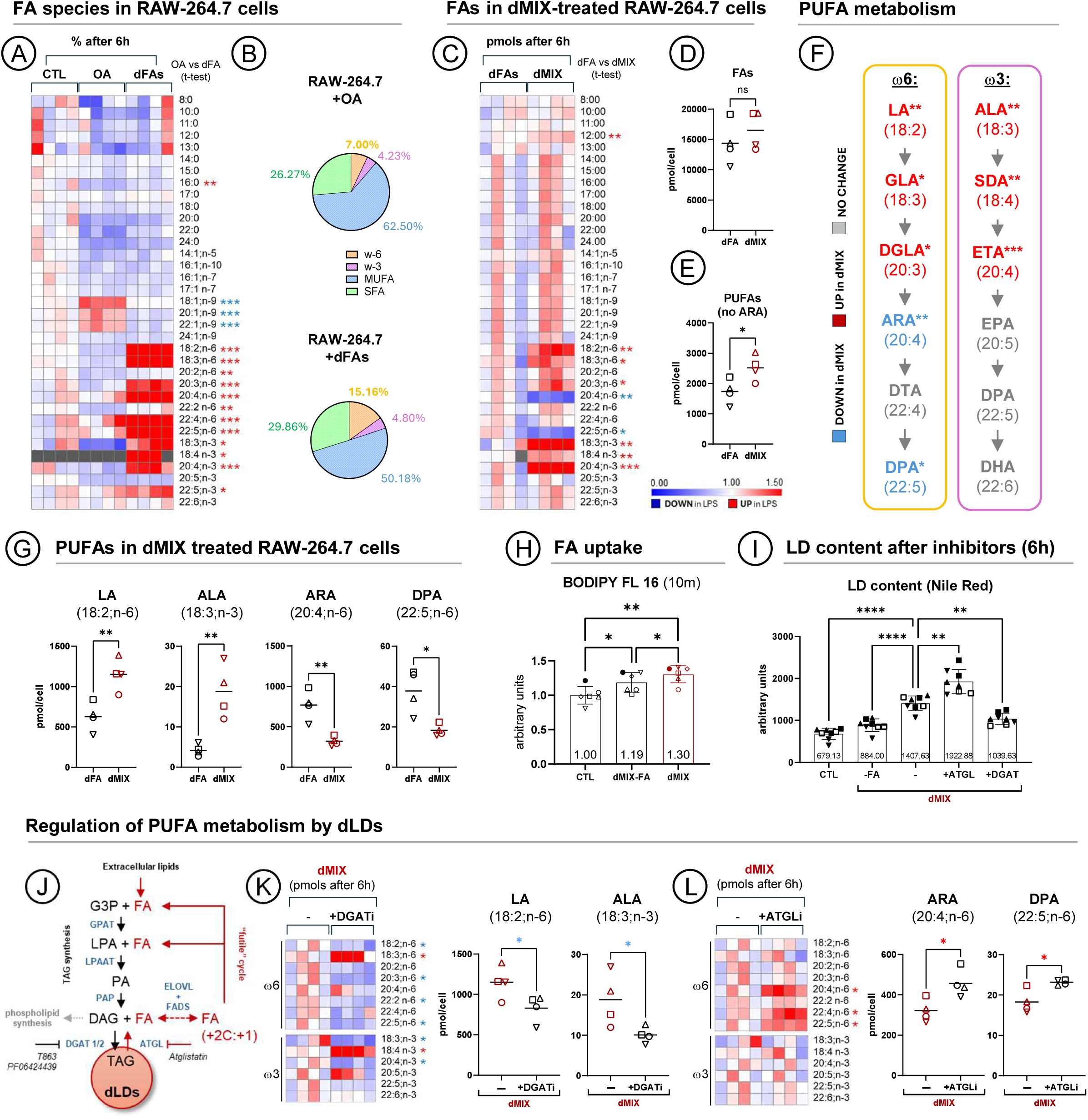
Defensive lipid droplets regulate PUFA metabolism in macrophages. **(A)** RAW-264.7 cells were treated for 6 hours with 15 µg/ml OA or dFAs (as in 6H). Untreated (CTL) and treated cells were analysed by targeted lipidomics to identify FAs. The heatmap compares the relative abundance (%) of each FA in CTL- and treated cells, as normalized to the average of control samples. Red and blue asterisks indicate respectively that the FAs is significant enriched or reduced when FA-treated cells are compared with OA-treated cells. Combined from N=4; *P < 0.05, **P < 0.01, ***P < 0.001, ****P < 0.0001 in an unpaired t-test. Raw data in Table S5. **(B)** The circular diagrams compare the percentage (%) of FA species in RAW-264.7 cells treated with OA or dFAs (Fig. 8A). **(C)** RAW-264.7 cells were treated for 6 hours with 15µg/ml of dFAs (as in 6H) or with the dMIX (also containing 15µg/ml of dFAs). Treated cells were analysed by targeted lipidomics to identify FA species. The heatmap compares the relative abundance (%) of each FA between dFAs- and dMIX-treated cells, as normalized to the average of dFA-treated samples for each species. Red and blue asterisks indicate respectively that the FAs is significantly enriched or reduced in dMIX-treated macrophages. Combined from N=4; *P < 0.05, **P < 0.01, ***P < 0.001, ****P < 0.0001 in an unpaired t-test. Raw data in Table S4. **(D and E)** The graphs show the mean (pmols) of total FAs (D) or PUFAs (E) in dFA-and dMIX-treated cells. In (E), ARA is not included in the calculations due to its high consumption. Each symbol indicates an independent experiment (N=4); ns is not significant and *P < 0.05 in an unpaired t-test. Raw data in Table S5. **(F)** Schematic representation of PUFA metabolism. Red and blue letters respectively indicate if the PUFA is significantly enriched or reduced in dMIX-treated cells when compared with dFA-treated cells (% as in 8C). PUFAs in grey are not significantly modified. Raw data in Table S5. **(G)** The graphs show the average concentration (pmols) of the most abundant PUFAs in dFA- and dMIX-treated macrophages (as in Fig. 8C). Each symbol indicates an independent experiment. Mean combined from N=4; *P < 0.05, **P < 0.01 in an unpaired t-test. Raw data in Table S5. **(H)** RAW-264.7 cells treated for 4 hours as in (Fig. 8C) and then incubated for 10 minutes with a fluorescent FA-BODIPY. FA uptake was determined by flow cytometry. Graph shows mean±SD (combined from at least N=3); *P < 0.05, **P < 0.01 in an unpaired t-test. **(I)** RAW-264.7 cells treated for 6 hours as in (Fig. 8C) and the LD content determined by flow cytometry (Nile Red staining). Some dMIX treated cells were simultaneously incubated with DGAT-1 and -2 inhibitors (T863 and PF06424439; DGATi) or with an ATGL inhibitor (Atglistatin; ATGLi). Each symbol indicates an independent experiment. Graph shows mean±SD (combined from N=4); **P < 0.01, ****P < 0.0001 in an unpaired t-test. **(J)** Schematic representation of the enzymatic reactions (grey letters) driving LD formation (TAG synthesis). Lipid intermediates are indicated with black letters, enzymes with blue letters, and FAs indicated in red. The reactions blocked by the inhibitors used in (Fig. 8I) are indicated. The futile cycle driven by elongases and desaturases to convert PUFA precursors in complex PUFAs is included as a potential mechanism to enrich dLDs in complex TAGs and PLs. **(K)** RAW-264.7 cells treated for 6 hours with dMIX in the presence or absence of DGATi (as in Fig. 8I). Treated cells were analysed by targeted lipidomics to identify FAs. The heatmap compares the relative abundance (%) of each FA between dMIX- and dMIX+DGATi-treated cells, as normalized to the average of dMIX-treated samples for each species. Red and blue asterisks indicate respectively the FAs significantly enriched or reduced in dMIX+DGATi-treated cells. The graphs show the significant reduction of ω6 and ω3 precursors in DGATi-dMIX-treated cells. Mean (combined from N=4); *P < 0.05 in an unpaired t-test. Each symbol indicates an individual experiment. Raw data in Table S5. **(L)** RAW-264.7 cells treated and analysed as in (Fig. 8K) but in the presence or absence of ATGLi. The graphs show that ARA and DPA consumption is significantly reduced in ATGLi-dMIX-treated cells. Mean (combined from N=4); *P < 0.05 in an unpaired t-test. Each symbol indicates an individual experiment. Raw data in Table S5.

Next, we determined whether the activation of innate immunity enhances PUFA metabolism. RAW-264.7 cells were treated for 6 hours with just dFAs or the dMIX and cellular changes in FAs quantified (Fig. 8C). At this early time-point, the dMIX-treated macrophages started to accumulate more FAs than non-activated cells (not statistically significant, Fig 8D) but already exhibited a significant accumulation of PUFAs (red asterisks, Figs. 8C, 8E, and S5). The accumulation of PUFAs was especially significant for ω-6 and ω-3 precursors (18 and 20 carbons) (Figs. 8F and 8G). As observed *in vivo* in LPS-treated animals, ARA was significantly consumed in dMIX-treated cells (Figs. 8C, 8F, and 8G). A significant reduction of DPA (ω6; 22:5) was also evident (Figs. 8F and 8G), a consumption also observed in some *in vivo* dLDs (Fig. 2D). The accumulation of FAs in dMIX treated cells when compared with non-activated macrophages could be explained, at least partially, by the increased capacity of dMIX-treated cells to internalise/retain extracellular FAs (Fig. 8H). Thus, as occurring *in vivo*, cultured macrophages activated with the dMIX accumulate and consume PUFAs.

Next, we determined the involvement of dLDs in the enhanced host PUFA metabolism. Thus, RAW-264.7 cells were activated for 6 hours with the dMIX and additionally treated with LD inhibitors. To reduce LD formation, we used T863 and PF06424439 to inhibit DGAT1 and DGAT2, the last enzymes in TAG synthesis. To reduce LD metabolism, we inhibited ATGL, the first and rate-limiting lipase hydrolysing TAGs, with used Atglistatin (Fig. 8J). In dMIX treated cells, the effect of the compounds was observed after just 6 hours; the inhibition of DGATs (DGATi) partially decreased LD formation and the ATGL inhibitor (ATGLi) increased the accumulation of LDs (Figs. 8I and 8J). Importantly, cells simultaneously treated with dMIX and DGATi, accumulated less PUFAs (blue asterisk, Fig. 8K), which was especially significant with a striking reduction of ω-6 and ω-3 precursors (graphs, Fig. 8K). Further, cells simultaneously treated with dMIX and ATGLi, significantly reduced the consumption of ARA and DPA (Fig. 8L). Therefore, dLDs coordinate immune PUFA metabolism and accumulate and supply these PUFAs.

### Defensive LDs regulate inflammation, signalling, and bacterial clearance

Finally, we analysed the role of dLD-PUFAs in innate immunity. Our previous IPA analysis of the enriched proteins on LPS-LDs ^15^ predicted that dLDs could be simultaneously involved in immune signalling, inflammation, and bacterial killing. The rapid formation of dLDs and dynamic PUFA metabolism in dMIX-activated RAW-264.7 macrophages offers a useful experimental tool to examine all these functions in a single cell model.

To analyse if dLDs modulate immune signalling, RAW-264.7 cells were treated for 3 hours with the dMIX but in the presence of LD inhibitors and expression of representative down-stream genes of the NFκB and IFNs pathways such as *Tnf* and *Il-6* or *Ifnβ* and *viperin* analysed by qRT-PCR. As expected, the dMIX promoted a robust activation of these pathways in RAW-264.7 macrophages and in BV2 microglia, and AML12 hepatocytes (Fig S6). Surprisingly, when cells were activated in the presence of LD inhibitors we observed a dual effect. While DGATi reduced the expression of *Il-6* and *Ifnβ*, the inhibitor increased expression of *Tnf* and had a little effect on *viperin* (Fig. 9A). ATGLi reduced the expression of *Il-6* but significantly increased *viperin* expression without affecting *Ifnβ* or *Tnf* (Fig. 9A). The FA composition of the dMIX did not significantly modify the expression of the immune mediators (Fig. S6). Thus, dLDs are indeed regulatory signalling platforms that differentially modulate the expression of different immune mediators, a process apparently not involving PUFAs but other ATGL-provided FAs.

**Figure 9.**
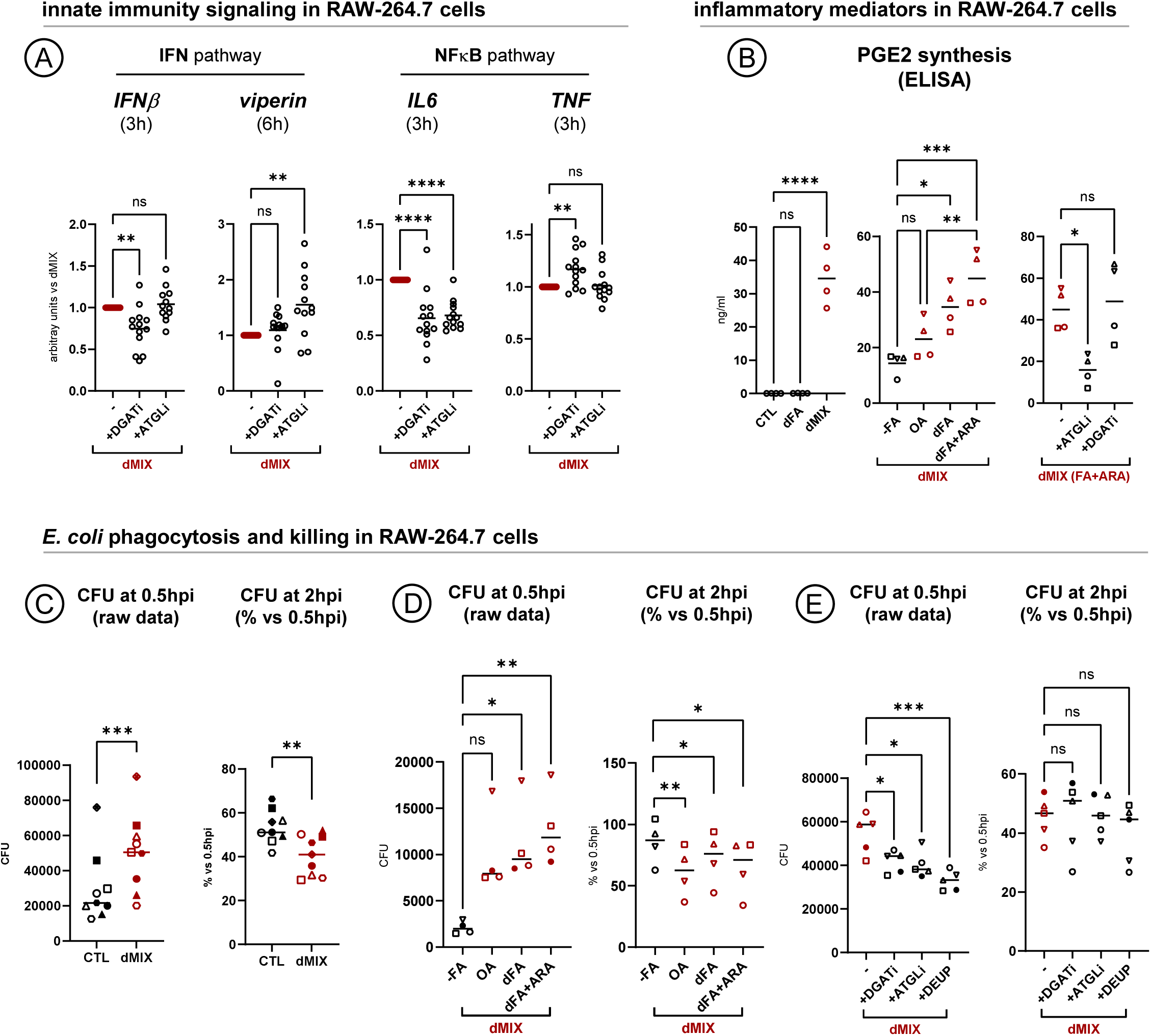
Role of defensive lipid droplets in innate immunity. **(A)** RAW-264.7 cells were treated for 3 or 6 hours with the dMIX in the absence or presence of DGATi or ATGLi. The mRNA expression of *IFN*β, *Viperin*, *IL-6*, and *TNF* was analysed by qRT-PCR. The graphs show the mean (combined from at least N=12); ns is not significant, **P < 0.01, ***P < 0.001, ****P < 0.0001 in an ordinary one-way ANOVA. **(B)** RAW-264.7 cells were treated for 6 hours with 15μg/ml of dFAs or the dMIX (containing 15μg/ml of dFAs) and the concentration of PGE2 secreted to the media determined by ELISA (left graph). In some experiments cells were treated with the dMIX but modifying its lipid composition: without FAs (-FAs), 15μg/ml of only OA, 15μg/ml of dFAs, or 15μg/ml of dFAs additionally containing 5% ARA (central graph). In other experiments, cells were treated with the dMIX supplemented with ARA (dFA+ARA) and simultaneously incubated with DGATi or ATGLi (right graph). The graphs show the mean (combined from N=4); ns is not significant, *P < 0.05, **P < 0.01, ***P < 0.001, ****P < 0.0001 in an ordinary one-way ANOVA. Each symbol indicates an independent experiment. **(C)** RAW-264.7 cells were left untreated or treated for 4 hours with the dMIX (containing 15μg/ml dFAs) and then infected with *E. coli*. Bacterial loads were quantified early after the infection reflecting phagocytosis (0.5 hours post-infection, 0.5hpi) and after 2 hours (2hpi). The left graph shows bacterial units (CFU) and the right graph the percentage (%) of CFU remaining after 2hpi relative to the initial CFU (0.5hpi). Each symbol represents an independent experiment. Graphs show the mean (combined from N=9); **P < 0.01 and ***P < 0.001 in a paired t-test. **(D)** RAW-264.7 cells were treated overnight with the dMIX supplemented with different lipids: without FAs (-FAs), with 15μg/ml of OA (OA), 15μg/ml of dFAs, or with 15μg/ml dFAs additionally supplemented with 5% ARA (dFA+ARA). Cells were then infected with *E. coli*. Bacterial loads were quantified early after the infection reflecting phagocytosis (0.5 hours post-infection, 0.5hpi) and after 2 hours (2hpi). Each symbol indicates an independent experiment. The graphs show the mean (combined from at least N=4); ns is not significant, *P < 0.05, and **P < 0.01 in an ordinary one-way ANOVA (left graph) or in a paired t-test (right graph). **(E)** RAW-264.7 cells were treated for 4 hours with the dMIX supplemented with 15μg/ml of dFAs and additionally incubated with the DGATi, ATGLi, or DEUP. Cells were then infected with *E. coli*. Bacterial loads were quantified early after the infection reflecting phagocytosis (0.5 hours post-infection, 0.5hpi) and after 2 hours (2hpi). Each symbol indicates an independent experiment. The graphs show the mean (combined from at least N=5); ns is not significant, *P < 0.05, ***P < 0.001 in an ordinary one-way ANOVA (left graph) or in a paired t-test (right graph).

Next, we analysed the involvement of dLD-PUFAs in the synthesis of pro-inflammatory prostaglandins. RAW-264.7 cells were treated with the dMIX for 6 hours and secretion of PGE2 measured by ELISA. Control cells or cells just treated with dFAs did not produce PGE2 but the treatment with the dMIX robustly increased levels of secreted PGE2 (Fig. 9B). Under these conditions the FA composition of the dMIX tightly determined PGE2 synthesis (Fig. 9B). When the dMIX contained OA slightly increased PGE2 production while when containing dFAs significantly increased the synthesis of PGE2. Further, when the dMIX was additionally complemented with 5% of ARA, RAW-264.7 cells display the highest capacity for PGE2 synthesis (Fig. 9B). The ARA used to produce PGE2 is largely provided by dLDs because ATGLi dramatically reduced the synthesis of PGE2 in activated macrophages, even in the presence of additional ARA (Fig. 9B). DGATi did not significantly affect PGE2 synthesis in agreement with some reports ^31^ but not with others ^23^, perhaps indicating cell type specific effects or that DGATi only partially inhibited dLD formation in RAW-264.7 cells (Fig. 8I).

Finally, we investigated the role of dLD-PUFAs in bacterial clearance. Initially, for these experiments RAW-264.7 cells were pretreated for 4 hours with the dMIX, then infected with *E. Coli*, and phagocytosis (30 minutes post-infection, 0.5hpi) and bacterial killing (2 hpi) estimated as viable colony forming units (CFU). While we did not observe significant changes in cells treated with dFAs (Fig. S7A), the macrophages pretreated with the dMIX exhibited an enhanced capacity for phagocytosis (Fig. 9C left) and killing (Fig. 9C right). Similar results were obtained when human monocyte-derived macrophages (HMDM) obtained from healthy donors were preincubated with the dMIX and then infected with *E. coli* (K12) (Fig. S7B). These processes strictly depend on FAs, as when the dMIX was not supplemented with FAs (-FAs, Fig. 9D), the phagocytic uptake and the antibacterial capability of macrophages was not enhanced (black symbols, Fig. 9D). When compared with OA, the addition of dFAs and specially the supplementation with ARA improved bacterial uptake (Fig. 9D left), suggesting the involving of PUFAs in these early events. In contrast, the composition of FAs did not affect the killing capacity of dMIX-treated cells (Fig. 9D right), likely reflecting that bacterial killing is a process mediated by the protein component of dLDs ^15^. The direct involvement of dLDs in these processes was tested with LD inhibitors. Both DGATi and ATGLi reduced the phagocytic capacity of dMIX-pretreated macrophages (Fig. 9E left). The involvement of dLD lipids in phagocytosis was further confirmed with the general lipase inhibitor DEUP, that efficiently reduced bacterial loads (Fig. 9E left). However, DGATi did not significantly reduce the percentage of bacterial killing (Fig. 9E right), probably because the inhibitor only partially reduces dLD formation (Fig. 8I) and because the processes of phagocytosis and bacterial killing are coupled. That is, antagonising LD-mediated bacterial uptake likely also compromises LD-mediated bacterial killing. Neither ATGLi nor DEUP reduced the killing capacity of cells, suggesting again that dLD lipids are not directly involved in killing but function as protein platforms. In conclusion, these results deliniate a crucial antibacterial function of dLDs, particularly during the initial steps of bacterial phagocytosis and demonstrate the ability of dLD-PUFAs to potentiate the capacity of macrophages for bacterial clearance.

## Discussion

The accumulation of LDs is a common trait of infected cells. Rather than passive bystanders of infection, progressive understanding of the organelle’s cell biology supports the view that host LDs organise generic innate immune defences. In contrast to the LDs of healthy cells (mLDs) regulated by energetic programs, dLDs are formed and activated in response to an intricate crosstalk between innate immune programs and lipid metabolism. *In vivo*, dLDs show reduced size and differential protein ^15^ and lipid composition with respect to mLDs. To gain mechanistic details of the unique biogenesis and functioning of dLDs, we designed the dMIX for reproducing in cell models the complex systemic environment observed in LPS-treated mouse livers. The dMIX combines PAMPs and cytokines predicted *in-silico* with the PUFA-enriched combination of FAs determined by lipidomics. The LDs induced by the dMIX in cells resemble the dLDs formed *in vivo* by i) rapidly forming in immune “professional” and non-professional” cells ii) accumulating defensive LD proteins, iii) having an increased PL/total lipid ratio and net accessible surface, iv) coordinating PUFA metabolism, and v) simultaneously mediating a plethora of innate immunity responses such as cytokine expression, lipid-mediated inflammation, and bacterial clearance. Specific LD inhibitors prove that these roles are conducted by proteins, lipids, or both and thus, that dLDs are platforms for organising defensive proteins and suppliers of defensive lipids.

The high purity of the LD fractions from the liver of LPS-treated mice analysed here provided an unprecedented resolution in the detection of LDs lipids allowing both to identify major LD lipids *in vivo* and to detect significant changes in otherwise undetectable minor lipid species (i.e. ARA-PC and ARA-PE, 0.1mol % total lipid). When compared to mLDs, dLDs are significantly enriched in complex PUFAs. The FA composition of LDs largely reflects the FA composition of LPS-serum. Endotoxins like LPS and cytokines, such as IL-6 and TNF, stimulate lipolysis in the adipose tissue of humans and mice ^39–42^, suggesting that adipocytes are a likely source of dLD PUFAs. In parallel, dMIX-stimulated macrophages demonstrate an enhanced capacity for FA uptake ^31^, an ability that explains the synergistic effect of cytokines and FAs on dLD formation. In cultured macrophages, arriving PUFA precursors are rapidly transformed into complex PUFAs, a process robustly activated by the dMIX. The dLD lipidomics demonstrate that PUFAs are accrued by dLD-TAGs and -PLs. Such an active metabolism may indicate an innate immune-stimulated TAG futile substrate cycle mediating progressive FA elongation and desaturation ^43^. The TAG futile cycle is likely coupled to an active lysophospholipid-PL Lands’ cycle because dLD-PLs are also enriched in complex PUFAs.

PUFAs have multiple roles in cell biology ^44^ and innate immunity ^45^, including pathogen recognition, signalling modulation, and control of gene expression. Here, we demonstrate a key role of dLDs in these processes by coordinating host PUFA metabolism and by supplying PUFAs for the defence. In our cell systems, dLD-PUFAs mediate inflammation and phagocytosis but are not apparently involved in the transcriptional regulation of cytokines or in bacterial killing.

Among PUFAs, ARA is the most actively metabolised during the initial phases of innate immunity. ARA is consumed by dMIX-activated macrophages but not by quiescent macrophages, indicating that it feeds innate immunity processes. Indeed, ARA is released from dLDs by ATGL to mediate in the synthesis of PGE2 and to enhance the phagocytic capacity of macrophages. ATGL hydrolyses TAG species at the sn-2 and sn-1 position but also possesses phospholipase A2 and transacylase activities ^46^ and thus, both dLD-TAGs and -PLs could be hydrolysed by ATGL. The lipidomic analysis suggest that PC and PE, but not PI, are major ARA donors. In LPS-activated hepatocytes the surface occupied by dLD-PLs can be higher than that of the plasma membrane clearly reflecting the extraordinary potential of dLD-PLs to function as PUFA donors. Furthermore, ATGL mediates PUFA-independent processes including transcriptional regulation of defensive cytokines such as IL-6 and defensive proteins such as viperin.

The reduction of macrophage’s phagocytic capacity when dLD formation or ATGL are inhibited is an intriguing phenotype also observed in other model systems ^23^. ATGL deficient macrophages exhibit impaired phagocytosis, a deficit attributed to reduced PPAR signalling ^47^ or defective small Rho GTPase activation ^48^. As occurring for eicosanoid synthesis, an appealing possibility is that dLDs provide lipids to generate and mature phagolysosomes. The exchange of lipids between LDs and late endosomal compartments is well documented ^49,50^. A recent multi-spectral organelle imaging approach demonstrated that, in activated macrophages, dLDs are highly dynamic organelles forming three- and four-way interactions with other organelles ^31^. Such a dynamic behaviour could facilitate the exchange of defensive PUFAs, other FAs, and even PLs between dLDs and host organelles.

In conclusion, overall LPS promotes changes in approximately 15 mol% of the dLD lipidome, mainly involving complex PUFA accumulation. Clarify the defensive roles of these complex PUFA species deserves further investigation. Because PUFAs are highly bioactive molecules, relatively small compositional variations could represent a robust biological response but also a security limit to avoid excessive inflammatory feedback or to reduce the PUFA-associated risk of lipotoxicity and ferroptosis ^44,51^. Indeed, the innate immune transcriptional programs studied here need to be rapidly switched off once the insult has been terminated. Chronic innate immunity and sustained inflammation are strongly associated with diseases characterised by excessive and long-term accumulation of LDs such as obesity and diabetes ^52^. Obese individuals with type 2 diabetes display a significant enrichment in PUFA-containing TAG species ^53^. Drugs targeting pro-inflammatory cytokine pathways have been approved for clinical use but have important side effects such as weight gain and increased susceptibility to infections ^52^. Therefore, dLDs and particularly ATGL emerge as promising therapeutic targets for treating these conditions. In this context, it will be interesting to explore both the likely role of dLD-PUFAs in the lipid-mediated termination of inflammation and how the inflammatory circuits controlled by dLDs are dysregulated in disease or aging. Indeed, understanding the cell biology of dLDs will undoubtably lead the search for urgently needed novel strategies to fight against resistant intracellular pathogens and to battle obesity-related inflammation and cardiometabolic disease.

## Abbreviations

ACSL: acyl-CoA synthetase long-chain
ALA: α-linolenic acid
ARA: arachidonic acid
ATGL: adipose triglyceride lipase
ATGLi: ATGL inhibitor (atglistatin)
CE: cholesterol ester
COX-2: cyclooxygenase-2
DAG: diacylglycerol
DEUP: diethylumbelliferyl phosphate
dFAs: defensive FAs (combination of 40% LA, 35% OA, 23% PA, and 2% ALA)
DGAT: diacylglycerol acyltransferase
DGATi: DGAT inhibitors (T863 and PF06424439)
DGLA: dihomo-γ-linolenic acid
dLD: defensive-LD
dMIX: defensive mix (combination of PAMPs, cytokines, and dFAs)
DPA: docosapentaenoic acid
EM: transmission electron microscopy
FA: fatty acid
FASN: fatty acid synthase
FC: free cholesterol
hpi: hours post infection
IFN: interferon
Igtp: interferon gamma-induced GTPase
IL: interleukin
IPA: Ingenuity Pathway Analysis
LA: linoleic acid
LC/ESI-MS/MS: liquid chromatography-electrospray ionization-tandem mass spectrometry
LDs: lipid droplet
mLD: metabolic-LD
MUFA: monounsaturated fatty acid
NF-κB: nuclear factor kappa-light-chain-enhancer of activated B cells
OA: oleic acid
PA: palmitic acid
PAMP: pathogen associated molecular pattern
PC: phosphatidylcholine
PE: phosphatidylethanolamine
PGE: prostaglandin
PI: phosphatidylinositol
PL: phospholipid
PLA2: phospholipase A2
Plin2: perilipin-2: PS, phosphatidylserine
PUFA: polyunsaturated fatty acid
qRT-PCR: quantitative reverse transcription polymerase chain reaction
SA: stearic acid
SFA: saturated fatty acid
SM: sphingomyelin
TAG: triacylglycerol
TLR: toll-like receptor
TNF: tumour necrosis factor

## Acknowledgements

AP, CD, and RGP are supported by ERC Synergy Grant (ERC-2022-SYG, 101071784). AP and MB are granted by Ministerio de Ciencia, Innovación y Universidades (MICINN, PID2021-127043OB-I00). AP is supported by H2020-MSCA-ITN-2018 (953489). AP and AMP are granted by CAIXA RESEARCH HEALTH (HR23-00560). RGP was supported by an Australian Research Council (ARC) Laureate Fellowship (FL210100107). MJS is supported by a National Health and Medical Research Council (NHMRC) Investigator Grant (APP1194406). RS is recipient of the Marie Skłodowska-Curie Actions Postdoctoral Fellowship (MSCA-PF) granted by the European commission. MS-A was supported by contracts from Ramón y Cajal and Consolidación Investigadora 2023 programmes (RYC2020-029690 and CNS2023-144831 and funded by MCINN/AEI/ 10.13039/501100011033 and “European Union NextGenerationEU/PRTR). BK was supported by an Early Postdoc Mobility fellowship from the Swiss National Science Foundation (P2ZHP3_184024). AH is a recipient of a Beatriu de Pinós contract from the Generalitat de Catalunya (2022 BP 00043). Authors acknowledge the CERCA Programme/Generalitat de Catalunya and the Cytomics and Image Platforms of IDIBAPS for access to equipment, the Radiological Protection Technology Unit, and the Advanced Optical Microscopy facility at the Scientific and Technological Centres of the University of Barcelona (CCiTUB) for technical help. Authors are grateful for the use of the Microscopy Australia Research Facility at the Centre for Microscopy and Microanalysis (CMM) at The University of Queensland (UQ). Views and opinions expressed are however those of the author(s) only and do not necessarily reflect those of the European Union or the European Research Council. Neither the European Union nor the granting authority can be held responsible for them.

## Figure preparation

Figures were created using Microsoft Excel and PowerPoint (Microsoft 365 MSO). Images were edited with Adobe Photoshop CS3 software (Adobe Systems Inc.). GraphPad Prism 7 (GraphPad Software) was used to create graphs and calculate statistical significances. Heatmaps were created using Morpheus (https://software.broadinstitute.org/morpheus).

## Figure legends

**Figure S1.**
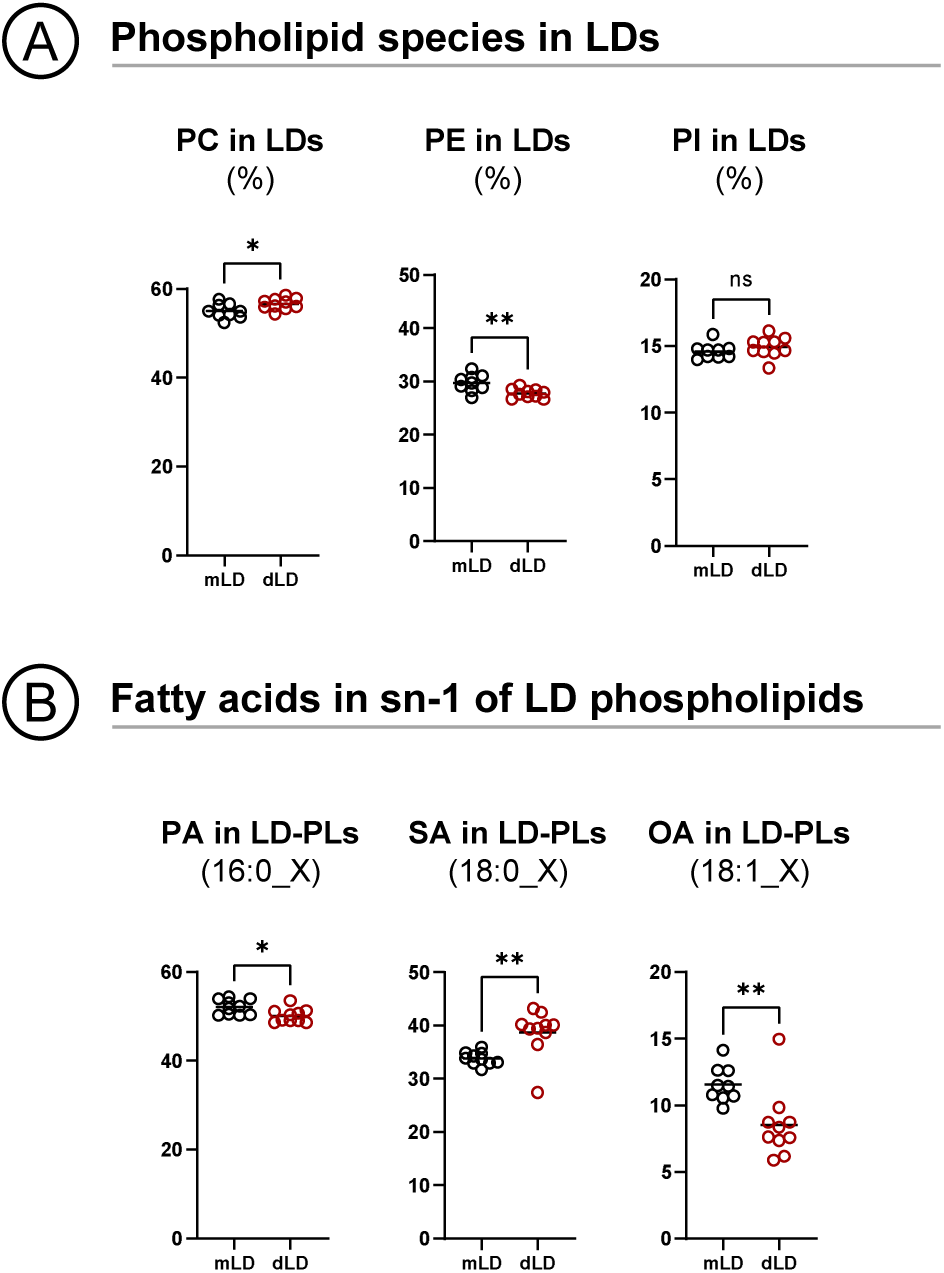
Phospholipids in dLDs. **(A)** LDs were purified as in (Fig. 1H) and PL identified by shotgun lipidomics. The graphs show the % of each family versus the total amount of phospholipid in mLDs and dLDs (N=9); ns is not significant, *P < 0.05, **P < 0.01, ***P < 0.001 in an unpaired t-test. Raw data in Table S1. **(B)** LDs were purified as in (Fig. 1H) and PL identified by shotgun lipidomics. The graphs show the % of PLs containing PA (palmitic), OA (oleic), or SA (stearic) in position sn-1 in mLDs and dLDs (N=9); *P < 0.05 and **P < 0.01 in an unpaired t-test. Raw data in Table S1.

**Figure S2.**
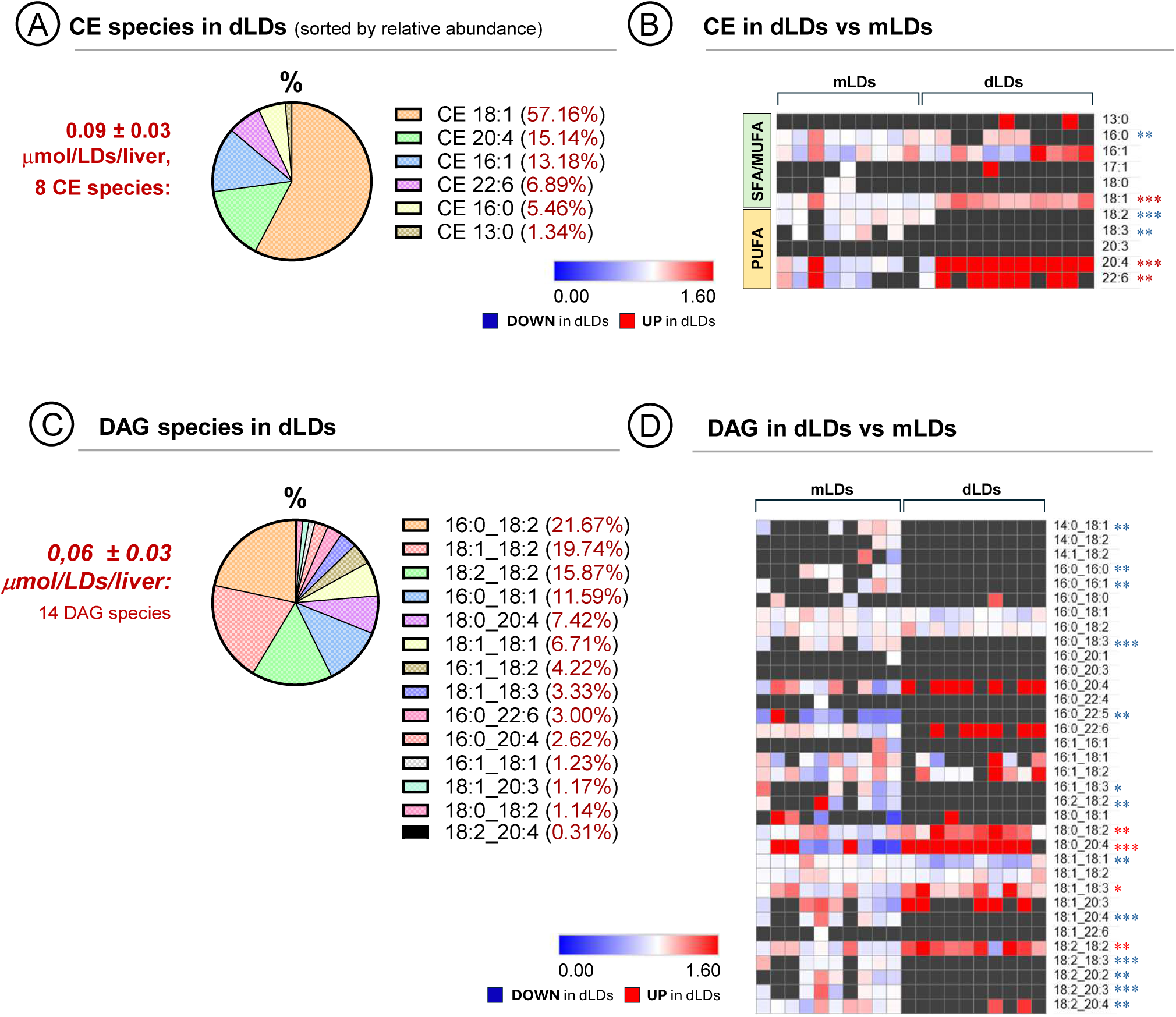
Cholesterol esters and diacylglycerol in defensive lipid droplets. **(A)** LDs were purified as in (Fig. 1H) and CE identified by shotgun lipidomics. The dLD fraction of each liver contained an average of 0.09 μmol of CE distributed among 8 different species (N=10). The box shows the relative abundance of CE species classified accordingly the associated FA. Raw data in Table S1. **(B)** Heatmaps showing the relative abundance (%) of each CE species in dLDs compared to mLDs, as normalised to mLD average. Red and blue asterisks indicate respectively that the CE is significantly enriched or reduced in dLDs (combined from N=10); **P < 0.01, ***P < 0.001 in an unpaired t-test. Raw data in Table S1. **(C)** LDs were purified as in (Fig. 1H) and DAG identified by shotgun lipidomics. The dLD fraction of each liver contained an average of 0.06 μmol of DAG distributed among 14 different species (N=10). The box shows the relative abundance of DAG species classified accordingly the associated FAs. Raw data in Table S1. **(D)** Heatmaps showing the relative abundance (%) of each DAG in dLDs compared to mLDs, as normalised to mLD average. Red and blue asterisks indicate respectively that the DAG is significantly enriched or reduced in dLDs (combined from N=10); *P < 0.05, **P < 0.01, ***P < 0.001, ****P < 0.0001 in an unpaired t-test. Raw data in Table S1.

**Figure S3.**
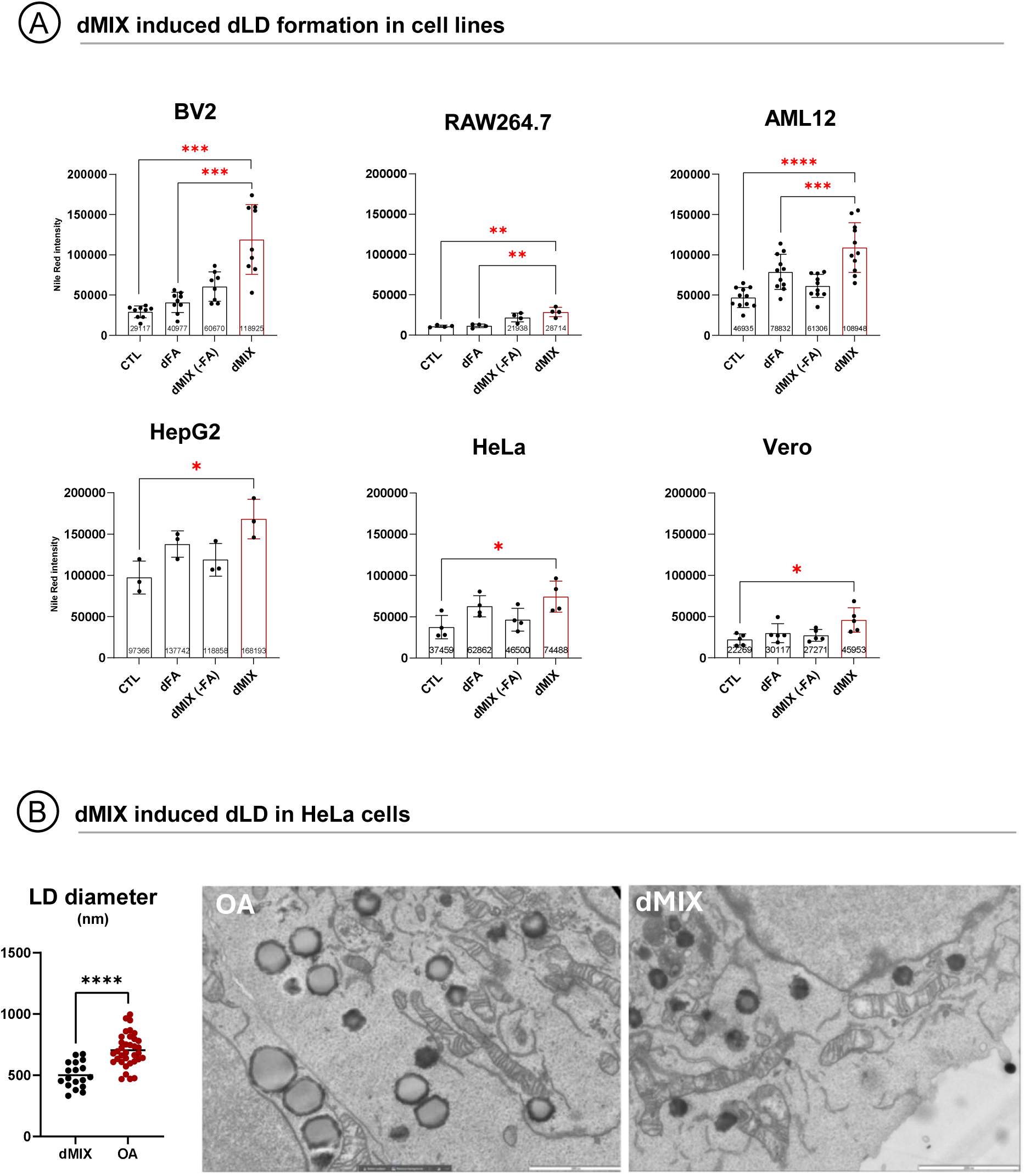
The dMIX induces dLDs in myeloid, hepatic, and fibroblast cell lines. **(A)** The indicated cell types were treated for 16 hours with a combination of dFAs (as in Fig. 6G; dFAs), a combination of PAMPs and cytokines (as in Fig. 6F; dMIX-FAs), or with both simultaneously (dMIX) and the LD content measured by flow cytometry. Graphs show means±SD (combined from at least N=3); *P < 0.05, **P < 0.01, ***P < 0.001, ****P < 0.0001 in a paired t-test. **(B)** HeLa cells were treated with OA or dMIX for 16 hours and then observed by TEM. The apparent diameter of at least n=18 LDs was measured in random sections (graph). Graph show mean±SD (N=1); ****P < 0.0001 in an unpaired t-test. The images show representative sections of cells treated with OA or the dMIX. Scale bar is 200nm.

**Figure S4.**
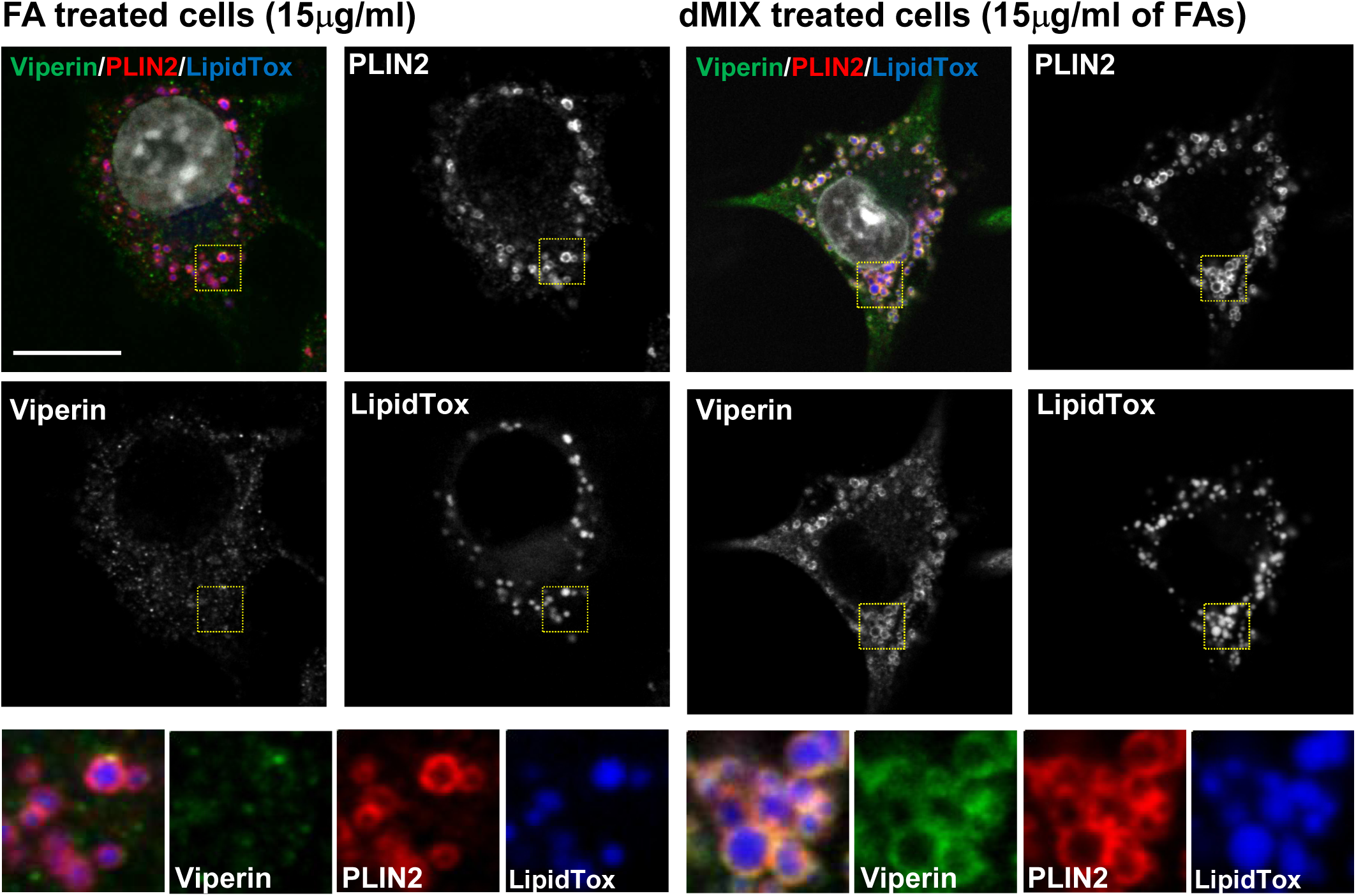
Endogenous Viperin on defensive lipid droplets. **(A)** Representative confocal images (N=3) of BV2 cells treated with OA (15µg/ml) to induce mLDs or dMIX (containing 15µg/ml of FAs: 40% LA, 35% OA, 23% PA, and 2% ALA). Cells were labelled with anti-Viperin (green) and anti-Plin2 (red) antibodies and LipidTox (LD, blue). Scale bar is 20µm. The yellow square marks the region selected for the high magnification panels.

**Figure S5.**
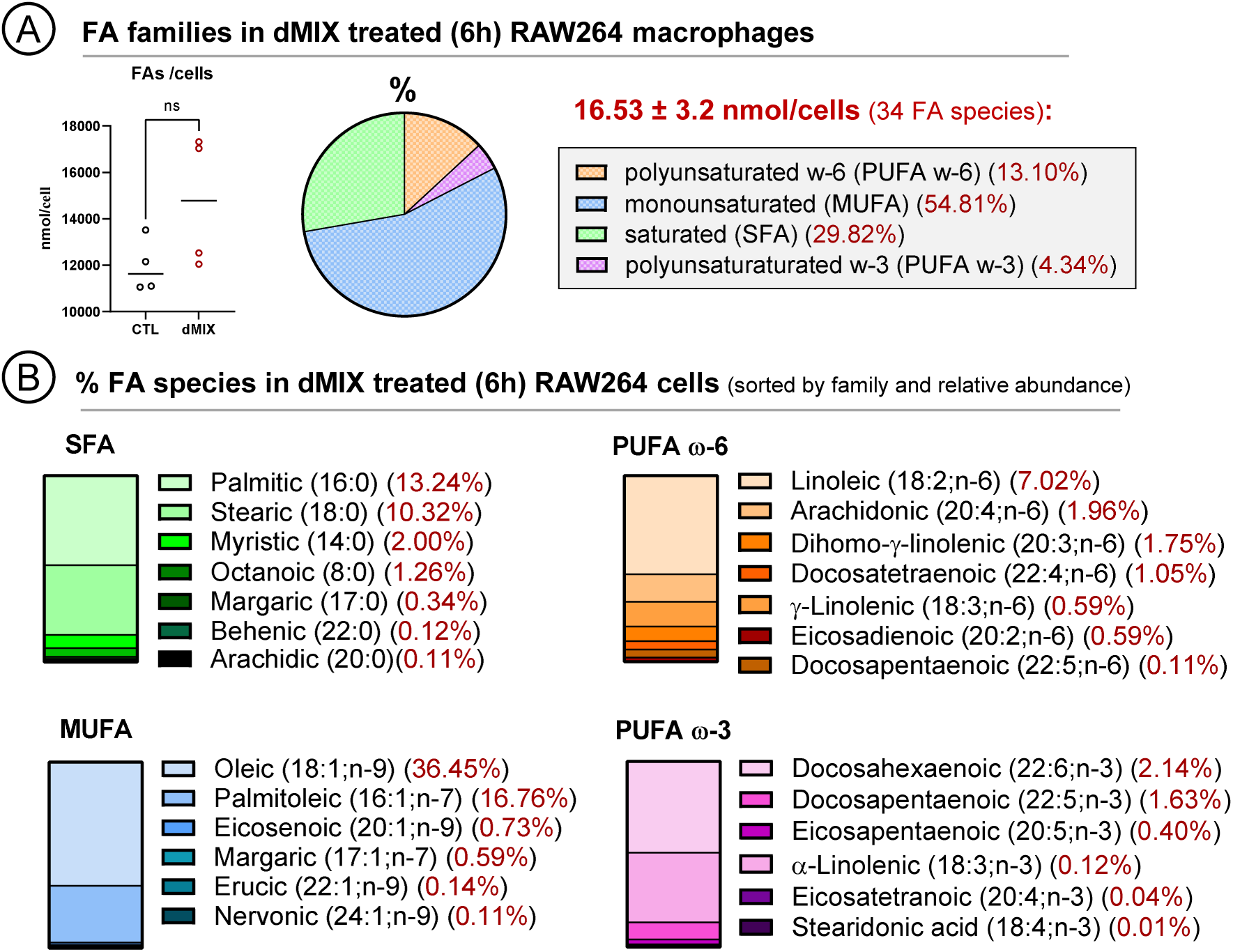
Fatty acid species identified in dMIX treated macrophages. **(A)** RAW-264.7 cells were treated for 6 hours with the dMIX (containing 15µg/ml of dFAs). Treated cells were analysed by targeted lipidomics (LC/ESI-MS/MS) to identify FA species. The graph shows the nmols of FAs identified in untreated (CTL) and dMIX-treated cells (N=4). RAW-264.7 cells treated with the dMIX contained an average of 16.53 nmol of FAs distributed among 34 different species. The box shows the relative (%) abundance of each FA species (combined from N=4). Raw data in Table S4. **(B)** Relative abundance of each FA when compared to the total amount of dLD FAs (combined from N=4). Raw data in Table S4.

**Figure S6.**
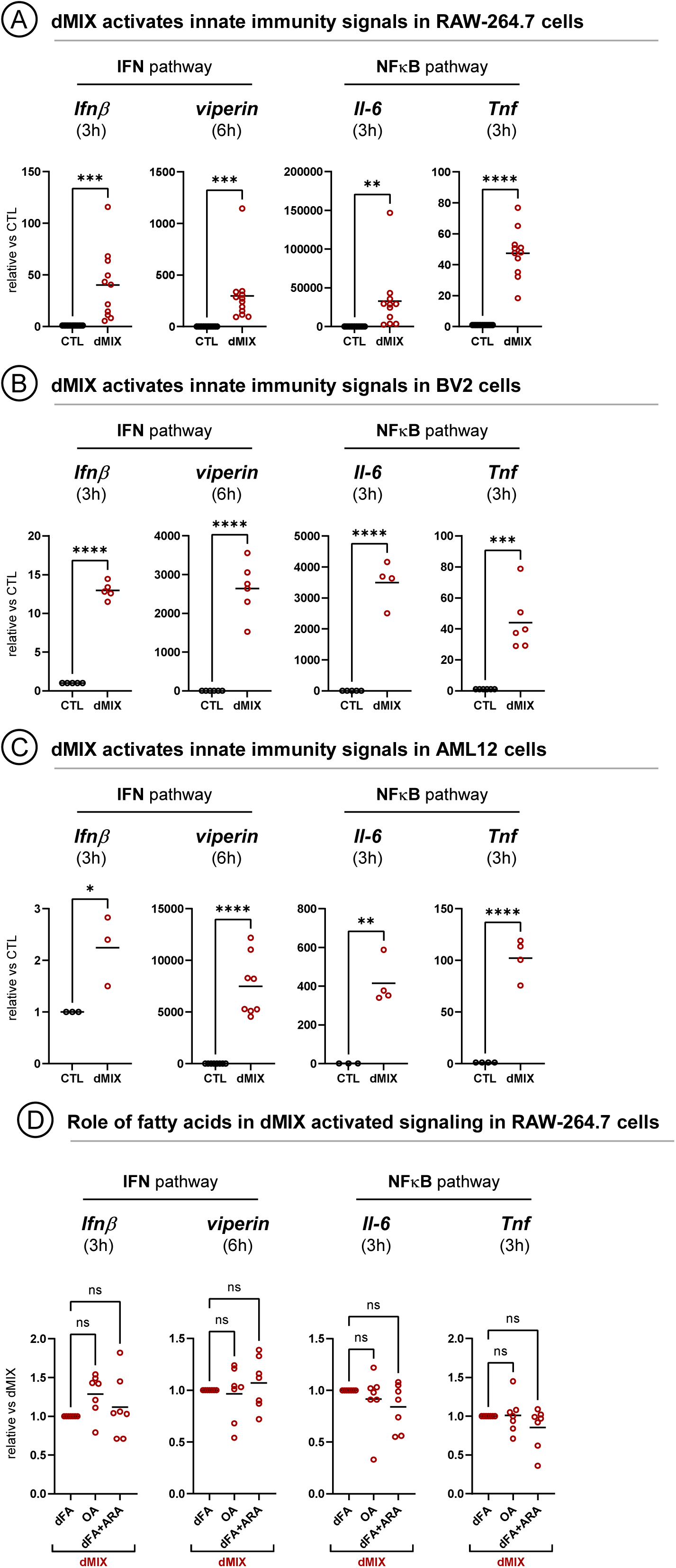
Defensive MIX and immune signalling. **(A)** RAW-264.7 cells were left untreated or treated for 3 or 6 hours with the dMIX supplemented with 15μg/ml of dFAs. The mRNA expression of *Ifn*β, *viperin*, *Il-6*, and *Tnf* was analysed by qRT-PCR. Combined from at least N=3; *P < 0.05, **P < 0.01, ***P < 0.001, ****P < 0.0001 in an unpaired t-test. **(B)** BV2 cells were treated and analysed as in Fig. S5A. **(C)** AML12 cells were treated and analysed as in Fig. S5A. **(D)** RAW-264.7 cells were treated for 3 or 6 hours with the dMIX supplemented with 15μg/ml of dFAs, 15μg/ml of only OA, or 15μg/ml of dFAs additionally supplemented with 5% of ARA. The mRNA expression of *Ifn*β, *viperin*, *Il-6*, and *Tnf* was analysed by qRT-PCR. The graphs show the mean (combined from at least N=7); ns is not significant in an ordinary one-way ANOVA.

**Figure S7.**
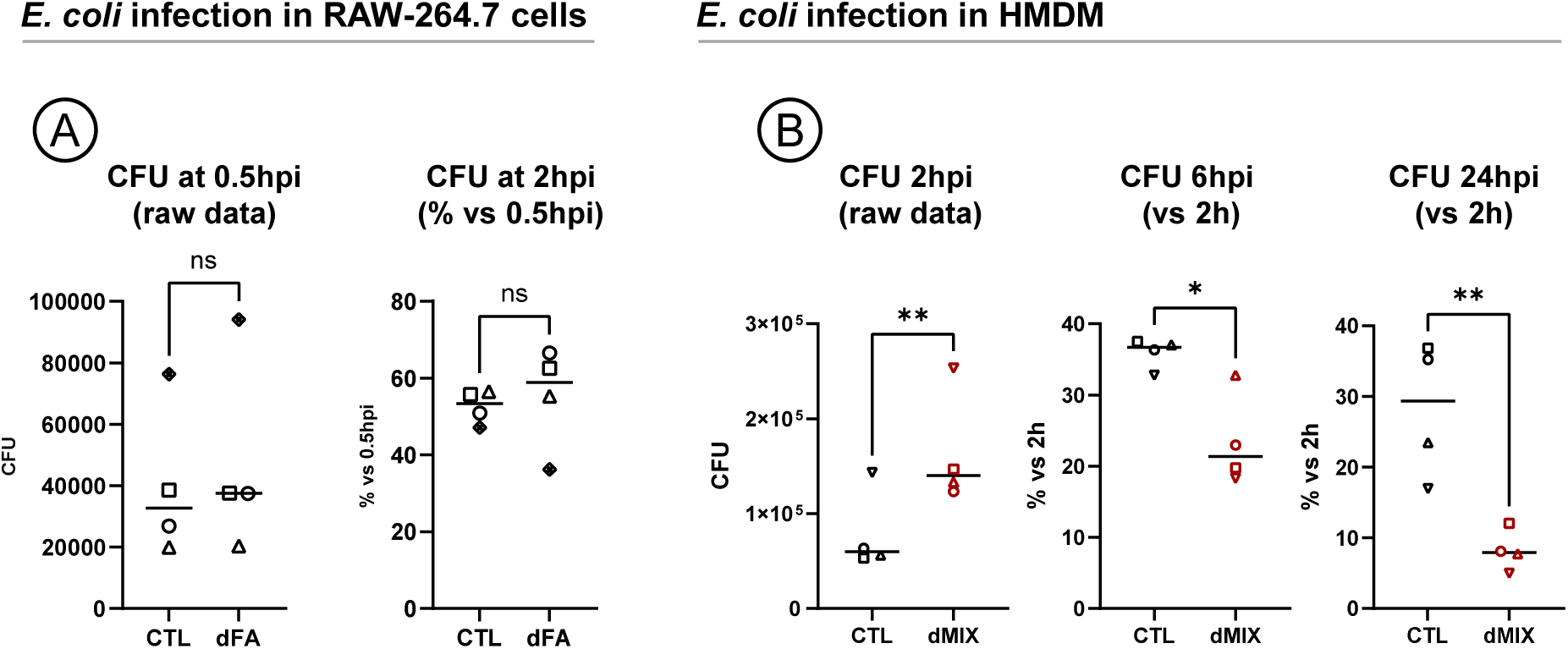
The dMIX increases the killing capacity of human macrophages. **(A)** RAW-264.7 cells were left untreated or treated for 4 hours with15μg/ml dFAs and then infected with *E. coli*. Bacterial loads were quantified early after the infection reflecting phagocytosis (0.5 hours post-infection, 0.5hpi) and after 2 hours (2hpi). The left graph shows bacterial units (CFU) and the right graph the percentage (%) of CFU remaining after 2hpi relative to the initial CFU (0.5hpi). Each symbol represents an independent experiment. Graphs show the mean (combined from N=9); **P < 0.01 and ***P < 0.001 in a paired t-test. **(B)** Human monocyte-derived macrophages (HMDM) from healthy donors were incubated for 16 hours with the dMIX (containing 15µg/ml of dFAs). Cells were then infected with *E. coli*. Bacterial loads in cells from (Fig. S4A) were quantified 2, 6, and 24 hours after infection. Each symbol represents an independent experiment. The left graph shows total bacterial units and the central and right graph the percentage (%) of bacterial units remaining at each time when compared to the initial loads (2hpi). Graphs show the mean (combined from N=4); *P < 0.05, **P < 0.01 in an unpaired t-test.

**Table S1. Shotgun lipidomics of lipid droplet lipid species**

The table includes the raw data corresponding to Figs. 1, 4, 5 and S1. The table contains the pmols of lipid species identified in LDs purified (N=10) from each liver as detailed in Fig.1.

**Table S2. Shotgun lipidomics of lipid droplet FAs**

The table contains the nmols of FAs identified in LDs purified (N=10) from each liver.

**Table S3. Targeted lipidomics of lipid droplet FAs**

The table includes the raw data corresponding to Fig. 2. The table contains the pmols of FAs identified in LDs purified from each liver as detailed in Fig.3.

**Table S4. Targeted lipidomics of mice serum FAs**

The table includes the raw data corresponding to Fig.3. The table contains the nmols of FAs identified in 1μl of mice serum obtained as detailed in Fig.3.

**Table S5. Targeted lipidomics of FAs in lipid-treated macrophages**

The table includes the raw data corresponding to Figs. 8 and S4. The table contains the pmols of FAs identified in macrophages cultured as detailed in Fig.8.

## Methods

### Reagents

Nile red (N3013), DAPI (D1306), BODIPY-FLC16 (D3821), BODIPY 493/503 (D3922) and LipidTox Deep Red (H34477) were from Thermo Fisher Scientific (Waltham, MA, USA). Trypsin/EDTA was from Life Technologies. FA-free bovine serum albumin (BSA; A8806), oleic acid (O1008), α-linolenic acid (L1376), palmitic acid (P5585), IL-1β (GF331), diethylumbelliferyl phosphate (DEUP; D7692), LPS (L2137), T863 (SML0539; DGAT1 inhibitor), Atglistatin (SML1075; ATGL inhibitor), Gentamycin (G1914), Triton X-100, Tween-20, Mowiol, were purchased from Sigma-Aldrich (St. Louis, MO, USA). TNF (300-01A), IL-1β (211-11B) and IFNC (315-05) were from Preprotech, Thermo Fisher Scientific. IFNβ (8234-MB) was from R&D Systems (Minneapolis, MN, USA). Pam3CSK4 and Poly(I:C) were from InvivoGen (San Diego, California, USA). PF-06424439 (HY-108341; DGAT2 inhibitor) was from MedChemExpress (New Jersey, NJ, USA). TVB-2640 was from SelleckChem (Cologne, Germany).

### Treatments

All fatty acids were conjugated to FA-free bovine serum albumin (BSA) at a molar ratio of 6:1 in medium. For the combination of FAs, the proportion used was 40% LA, 35% OA, 23% PA, and 2% ALA, at 1.77mM final concentration conjugated to albumin. The defensive mix (dMIX) is composed of IFNγ (1ng/ml), IFNβ (1ng/ml), IL-1β (1ng/ml), TNF (20ng/ml), LPS (5 µg/ml) and Pam3-CSK4 (1 µg/ml) and the combination of FAs at 15µg/ml unless it is specified. To quantify the size of dLDs, HeLa cells were treated for 6h with either 50µg/ml OA or dMIX (15µg/ml FA concentration).

### Cell culture

The human monocytic cell line THP-1 was obtained from ATCC (Rockville, Maryland) and cultured in RPMI 1640 medium (Gibco, ThermoFisher Scientific) supplemented with 10% heat inactivated FBS (Bovogen Biologicals, Melbourne, VI, Australia), 5 mM sodium pyruvate and 10mM HEPES (Gibco), 50 U/ml penicillin (Invitrogen, Carlsbad, California) and 50 mg/ml streptomycin (Invitrogen). Infection media are similar to complete media; except they do not contain penicillin-streptomycin. Human monocyte-derived macrophages (HMDMs) were obtained by differentiating CD14+ monocytes as previously described ^15^. BV-2 cells were kindly provided by Dr. Anna Planas. AML12 (ATCC, CRL-2254) were cultured in 1:1 mixture of Dulbecco’s modified Eagle’s medium (DMEM, Biological Industries, Cromwell, Connecticut) and Ham’s F12 medium with 0.005mg/ml insulin, 0.005mg/ml transferrin, 5ng/ml selenium, 40ng/ml dexamethasone (from Sigma-Aldrich’s ITS Media Supplement), supplemented with 10% v/v Fetal Bovine Serum (FBS, Biological Industries). HepG2 (ATCC HB8065), HeLa (ATCC CCL2), RAW-264.7 (ATCC TIB71), Vero (ATCC CCL81) cells were cultured in DMEM supplemented with 10% FBS (Biological Industries), 4mM L-glutamine, 1mM pyruvate (Sigma-Aldrich) and 50U/ml Penicillin, 50 mg/ml Streptomycin and non-essential amino acids (Biological Industries).

### Animals and treatments

C57BL/6J male mice (8 to 10 weeks old) were purchased from Charles River Laboratories (Wilmington, Massachusetts). All animal experiments were approved and carried out according to the Animal Care Committee of the University of Barcelona and received humane care in compliance with institutional guidelines regulated by the European Community. All mice were housed in a room with controlled humidity and light-dark cycles of 12 hours. To purify LDs animals were fasted overnight (16 h) and intraperitoneally injected with 200μl of LPS (6 mg/kg final dose) (L2639, Sigma-Aldrich, St Louis, Missouri) or in the case of the control animals with the same volume of saline buffer. To obtain serum, mice were intraperitoneally injected with 200μl of LPS (6 mg/kg final dose) (at 9:00 am) or saline buffer and blood was extracted two hours later by cardiac punction.

### Modelling of dLD upstream gene networks

To model gene networks potentially governing the dynamics of the dLD proteome and their interrelationship, we applied Upstream Regulation Inference analyses from the Ingenuity Pathway Analysis (IPA™) platform (v. 01-23), of the differential dLD proteomes as previously described^15^. From a list of candidate upstream regulators, we selected transcription factors with high prediction score (log (p value of overlap) > 5) that bound directly to regulatory regions of more than 5 dLD proteome members according to the TFEA.ChIP database ^54^. A gene network is then modelled from these factors and proximal regulators using the STRING v.12.0 allowing only for 5 additional connectors; three bona fide innate immunity LD effectors and three LD metabolic regulators from the dLD proteome were included in the network according to the relationships indicated by the IPA analyses. The network was rendered, highlighting edges connecting innate immunity and metabolic regulators, on the Cytoscape 3.9.1. platform.

### Liver fractionation

Livers were perfused with 0.9% NaCl, 0,1% EDTA solution, chopped with a scalpel for two min and transferred into a Dounce tissue grinder at a ratio of 1 gr liver tissue per 3ml homogenization buffer (25 mmol/L Tris-HCl pH 7.5, 100 mmol/L NaCl, 1 mmol/L EDTA, 5 mmol/L EGTA) and homogenized using a loose and tight pestle (3 strokes each). Homogenates were centrifuged at 500g at 4°C for 10 min and 2.5 ml of the supernatant were mixed with 2.5 ml of 2.5mol/L sucrose and placed at the bottom of the ultracentrifuge tube. 1.4 ml of 25%, 15%, 10%, and 5% (w/v) sucrose solution with homogenization buffer were carefully added on the top, with a final upper phase of 25 mmol/L Tris-HCl pH 7.5, 1 mmol/L EDTA, 5 mmol/L EGTA. After centrifugation at 12.000g, at 4°C for 1 hour (SW-41Ti rotor, Beckman

Coulter, Pasadena, California), six fractions were collected and stored at −80°C for western blotting analysis.

### Lipidomic studies

Mass spectrometry-based lipid analysis was performed by Lipotype GmbH (Dresden, Germany) as previouly described ^55^.

#### Sample Collection

For LD isolation, the upper LD phase obtained after liver fractionation was collected and re-centrifuged at 16.000g at 4°C for 10min. Concentrated floating LDs were obtained by removing the lower phase buffer with a syringe and were brought to a final volume of 1 ml with saline solution (0.9% NaCl) with 8% sucrose. Samples were stored at −80°C until sent to Lipotype for the lipidomic analysis. For mice serum studies, blood samples were collected via cardiac puncture, and serum was obtained after centrifugation at 6,000g for 15 min at 4°C in serum heparin separator tubes (Becton Dickinson, Franklin Lakes, New Jersey). Isolated serum was immediately frozen and stored at −80°C until lipid extraction. For RAW-264.7 cell analysis, 2×10^6^ cells were seeded in 6-well plates, and allowed to adhere for 16 h before treatments. Cells were pre-incubated for 2h with LD inhibitors (Atglistatin 70µM or a cocktail including T863 50µM and PF06424439 50µM) or left untreated, followed by 6h incubation with dMIX of different compositions or with fatty acids alone in RAW medium. LD inhibitors were included during dMIX treatment. Then, cells were washed with PBS, scraped from the plate, transferred into a 1,5 ml tube and centrifuged at 800 g for 5 min. Pelleted cells were stored at −80°C until lipid extraction.

#### Lipid extraction for mass spectrometry lipidomics

Lipids were extracted using a chloroform/methanol procedure ^56^. Samples were spiked with internal lipid standard mixture containing: cardiolipin 14:0/14:0/14:0/14:0 (CL), ceramide 18:1;2/17:0 (Cer), diacylglycerol 17:0/17:0 (DAG), hexosylceramide 18:1;2/12:0 (HexCer), lyso-phosphatidate 17:0 (LPA), lyso-phosphatidylcholine 12:0 (LPC), lyso-phosphatidylethanolamine 17:1 (LPE), lyso-phosphatidylglycerol 17:1 (LPG), lyso-phosphatidylinositol 17:1 (LPI), lyso-phosphatidylserine 17:1 (LPS), phosphatidate 17:0/17:0 (PA), phosphatidylcholine 15:0/18:1 D7 (PC), phosphatidylethanolamine 17:0/17:0 (PE), phosphatidylglycerol 17:0/17:0 (PG), phosphatidylinositol 16:0/16:0 (PI), phosphatidylserine 17:0/17:0 (PS), cholesterol ester 16:0 D7 (CE), sphingomyelin 18:1;2/12:0;0 (SM), triacylglycerol 17:0/17:0/17:0 (TAG) and cholesterol D6 (Chol). After extraction, the organic phase was transferred to an infusion plate and dried in a speed vacuum concentrator. The dry extract was re-suspended in 7.5 mM ammonium formiate in chloroform/methanol/propanol (1:2:4; V:V:V). All liquid handling steps were performed using Hamilton Robotics STARlet robotic platform with the Anti Droplet Control feature for organic solvents pipetting.

#### MS data acquisition

Samples were analysed by direct infusion on a QExactive mass spectrometer (Thermo Scientific) equipped with a TriVersa NanoMate ion source (Advion Biosciences). Samples were analysed in both positive and negative ion modes with a resolution of Rm/z=200=280000 for MS and Rm/z=200= Ejsing Ejsing for MSMS experiments, in a single acquisition. MSMS was triggered by an inclusion list encompassing corresponding MS mass ranges scanned in 1 Da increments (Surma et al. 2015). Both MS and MSMS data were combined to monitor CE, Chol, DAG and TAG ions as ammonium adducts; LPC, LPC O-, PC and PC O- as formiate adducts; and CL, LPS, PA, PE, PE O-, PG, PI and PS as deprotonated anions. MS only was used to monitor LPA, LPE, LPE O-, LPG and LPI as deprotonated anions, and Cer, HexCer and SM as formiate adducts.

#### Data analysis and post-processing

Data were analysed with in-house developed lipid identification software based on LipidXplorer ^57,58^. Data post-processing and normalization were performed using an in-house developed data management system. Only lipid identifications with a signal-to-noise ratio >5, and a signal intensity 5-fold higher than in corresponding blank samples were considered for further data analysis.

#### Lipidomics analyses

To represent as heatmaps the relative changes in lipid species composition between mLDs and dLDs, the relative proportions of each lipid species was computed separately for each biological replica. Then, for each lipid species, the average value obtained from mLDs replicas was used to normalise all mLD and dLD values of that specific lipid species, minimizing differences stemming from large variations in relative presence between different lipid species. The MORPHEUS analysis platform (https://software.broadinstitute.org/morpheus) was then used to generate heatmap graphs for each lipid class.

### Electron microscopy

Liver samples and cultured cells in 3-cm dishes were fixed in 2.5% glutaraldehyde, washed in PBS and post-fixed in 2% osmium tetroxide (OsO_4_) with 1.5% potassium ferricyanide. Samples were washed in water, incubated in 1% (w/v) thiocarbohydrazide solution and post-fixed again in 2% OsO4, washed in water and stained with 1% uranyl acetate. Subsequently, samples were further stained with a lead aspartate solution (20mM lead nitrate, 30mM aspartic acid, pH 5.5), dehydrated through a series of acetone solutions, infiltrated with Durcupan resin and polymerised. Ultrathin sections of 60 nm were obtained with an ultramicrotome (EM U26, Leica, Germany), collected on copper grids and imaged using a Hitachi 7700 (Tokyo, Japan) or JEOL 1011 TEM. Quantification of LD size and diameter from cultured cells and liver sections, was done using FIJI-Image software (WayneRasband, NIH) ^59,60^. To gain estimates of the LD surface area in control, fasted, and fasted/LPS-treated mice, small pieces of liver were fixed for electron microscopy (EM) as in previous studies ^15^. After EM processing, thin sections were prepared and imaged in a JEOL1011 TEM. Images of hepatocytes were captured systematically across the sections and then analysed using standard stereological techniques using intersection and point counting to measure the surface to volume ratio, Sv, of lipid droplets and hepatocyte plasma membrane (PM). Intersections with the compartments of interest (LD surface or PM) were related to points over the nucleus, which acted as a reference space of defined volume (mean diameter as measured by light microscopy = 9.8µm). Sv was calculated by measuring intersections (I) in one direction of a grid and points over the reference space (P) using the formula Sv = 2I/P.d where d (the distance between lines of the lattice grid) =1.2µm.

### Immunofluorescence

Cells grown in polylisine coated coverslips (72294-2; Electron Microscopy sciences, Hatfield, Pennsylvania) and treated as specified were fixed for 1 hour in 4% paraformaldehyde (Electron Microscopy Science, Hatfield, Pennsylvania) in PBS, permeabilized in 0.15% Triton X-100 in PBS for 5 min and blocked with 1% BSA (A7906, Sigma-Aldrich) 0,1% Tween-20 in PBS for 10 min at room temperature. Primary antibodies were diluted in blocking solution: mouse monoclonal anti-Viperin (1:100, ab107359, Abcam), mouse monoclonal anti-Igtp (1:100, sc-136317, Santa Cruz Biotechnology), rabbit polyclonal anti-Plin2 (1:200, 15294-1-AP, Proteintech). After 1 hour incubation and three PBS washes, cells were incubated with secondary antibodies diluted 1:300 with blocking solution: donkey anti-mouse IgG Alexa Fluor 488 (A21202), and donkey anti-rabbit IgG Alexa Fluor 555 (A321094), from ThermoFisher Scientific. Finally, cells were labelled with DAPI for 5 min (1:10.000) and LDs were stained with Bodipy 493/503 (1µg/ml) for 10 min, washed twice with PBS and coverslips were mounted with Mowiol. Alternatively, LDs were stained with LipidTox Deep Red at 1:50 dilution in Mowiol.

### Fluorescence microscopy

Images were taken in Leica TCS SP5 laser scanning confocal spectral microscope. Images corresponding to single confocal sections were taken using 405nm laser for DAPI detection, 488nm laser for Viperin and Igtp detection, 514nm laser for Plin2 detection and 633nm laser for LipidTox Deep Red detection, with a x63 oil immersion objective lens with a numerical aperture of 1,4 and a pinhole of 1.5. a.u. Images were analysed using Image J (NIH).

### Measurement of PGE2 concentration

PGE2 concentrations were measured in the supernatant of RAW-264.7 cells determined by a specific enzymatic immunoassay (EIA) using a commercial kit (Cayman Chemicals; 514010) and following the manufacturers protocol. Briefly, RAW-264.7 cells were pre-incubated for 2h with LD inhibitors (Atglistatin, 70uM; T863, 50uM; PF06424439 50uM) followed by 6h incubation with dMIX of different fatty acid compositions in RAW medium. Cell-free supernatants and PGE2 standards were diluted in assay medium, plated into the PGE2 EIA assay plate in duplicate and incubated for 18h at 4°C in the presence of PGE2 AChE tracer and PGE2 mAb. PGE2 EIA was developed with Ellman’s reagent, reconstituted immediately before use, and absorbance measured at a wavelength of 420 nm. Concentration was inferred from the measured absorbance using the standard curve.

### Western blots

Equal volumes of liver fractions were subjected to SDS-polyacrylamide gel electrophoresis (Mini-Protean, Bio-Rad, Berkeley, California) and proteins were transferred to nitrocellulose membranes (162-0115, Bio-Rad). After blocking for 1 hour in 5% non-fat milk Tween-TBS, membranes were incubated for 1hour with the following primary antibodies: rabbit polyclonal anti-Plin2 (1:5000; ab78920, Abcam), rabbit polyclonal anti-EEA1 (1:200; ab2900, Abcam), mouse monoclonal anti-Viperin (1:1000; ab107359, Abcam), mouse monoclonal anti-G M130 (1:2000; Labs 810822, BD-Biosciences San Jose, California), rabbit monoclonal anti-ACSL4 (1:5000; ab155282, Abcam), rabbit monoclonal anti-Caveolin-1 (1:2000; ab192869, Abcam). Membranes were washed and incubated with peroxidase-conjugated secondary antibodies (1:3000) goat anti-rabbit IgG (H+L)-HRP conjugate (1706515, Bio-Rad), goat anti-mouse IgG (H+L)-HRP conjugate (1706516, Bio-Rad), detected with ECL (Biological Industries) and visualised using ImageQuant LAS4000 (GE Healthcare, Chicago, Illinois).

### Quantitative real-time PCR

RNA extraction was performed using the using the EZ-10 DNAaway RNA Miniprep kit (BioBasic, Toronto, Canada) according to the manufacturer’s protocol. Complementary DNA (cDNA) was synthesized from 1 µg RNA with the High-Capacity cDNA Reverse Transcription Kit (Applied Bioscience, ThermoFisher Scientific) according to the manufacturer’s instructions. qRT-PCR was set up using Brilliant SYBR Green qPCR Master Mix (# 600548, Agilent Technologies, Santa Clara, California) and run onMx3000P QPCR System (Agilent Technologies). The sequences of the primers used were as follows: *IL-6*: forward 5’-TGGTACTCCAGAAGACCAGAGG-3’ and reverse 5’-AACGATGATGCACTTGCAGA-3’. *TNF*: forward 5’-CATCTTCTCAAAATTCGAGTGACAA-3’ and reverse 5’-TGGGAGTAGACAAGGTACAACCC-3’. *IFNβ*: forward 5’-TACACTGCCTTTGCCATCCA-3’ and reverse 5’- GAGGACATCTCCCACGTCAA-3’. *Plin2*: forward 5’- GCCCTGCCCATCATCCA-3’ and reverse 5’-GCAGTCTTTCCTCCATCCTGTC-3’. *ACSL4*: forward 5’-CCAGCAAAATCAAGAAGGGAAGC-3’ and reverse 5’-GCGCCGCCAGACAGCATCA-3’. *ATGL*: forward 5’-CACCATCCGCTTGTTGGAGT-3’ and reverse 5’- TCCCCCAGTGAGAGGTTGTT-3’. *Viperin*: forward 5’- CTTCAACGTGGACGAAGACA-3’ and reverse 5’-GACGCTCCAAGAATGTTTCA-3’. *Igtp*: forward 5’-AGTCTGCGACAAGTGCATCA-3’ and reverse 5’- TGTACTCCGAGCTACCTGCT-3’. *Cathelicidin*: forward 5’- CTGTGGCGGTCACTATCACT-3’ and reverse 5’-GTTCCTTGAAGGCACATTGC-3’.

### Flow cytometry

BV2, AML12, HepG2 (160.000 cells/plate), RAW264.7 (400,000 cells/plate), HeLa, and Vero (140,000 cells/plate) cells were seeded in 12-well plate and medium was replaced the following day with the corresponding treatments for 16 hours. To analyse LD content, cells were harvested and collected in microcentrifuge tubes, centrifuged at 700g at room temperature for 3 min, washed with PBS and fixed with 4% paraformaldehyde for 15 min. After a PBS wash, cells were stained in Nile Red (5 µg/ml) in PBS for 15 min, washed three times in PBS and analysed. To analyse FA uptake, cells were incubated with 1μM BODIPY®FLC16 for 10min in their culture medium and then harvested by trypsinization and resuspended in PBS and analysed using Cytoflex, BH15085 Flow Cytometer (Beckman Coulter).

### Bacterial Infection Assays

The bacterial strains used were *E. coli* K-12 MG1655 and *E. coli*-GFP (ATCC-25922). Bacteria were grown overnight in Luria-Bertani (LB) medium under shaking conditions. For HMDM infection, 400.000 cells were plated into 24 well plates, and 2h later media was replaced with either vehicle or dMIX. Cells were left overnight and 1h prior infection media was replaced with antibiotic free media. The overnight *E. coli* MG1655 was washed twice, resuspended in antibiotic-free cell culture medium and then used to infect HMDMs at multiplicity of infection (MOI) of 100. After 1h, extracellular bacteria were removed by replacing the media with fresh media containing 200 µg/ml Gentamycin for 1h, followed by incubation with fresh media containing 20 µg/ml Gentamycin for 2h, 6h, and 24h post-infection. Cells were lysed with 0.1% Triton X-100 in PBS, plated onto LB agar plates and the number of live bacteria was estimated by counting CFU after overnight incubation of the plates at 37°C. For RAW-264.7 infection, cells were seeded at 500.000 cells/plate in 12-well plate and next day before infection, medium was replaced with antibiotic-free culture medium. Cells were infected with *E. coli* at MOI of 10 for 30 min at 37°C and washed three times with culture media with high dose of gentamycin (200 µg/ml) to exclude any extracellular bacteria. To evaluate phagocytic activity, cells were collected immediately after washing (0.5 hours post infection, hpi). To determine bacterial killing activity, cell media was replaced with culture media with low dose of gentamycin (20 µg/ml) and incubated for 90 min at 37°C (2 hpi). Samples were collected after three PBS washes and incubation for 5 min at 37°C with 0.1% Triton X-100 in PBS.

#### Quantification of bacterial cells in RAW-264.7 infected cells

Serial dilutions of RAW-264.7 infected cell lysates (two biological replicates) in LB medium were cultured in a 96-well plate (three technical replicates) at 37°C and the absorbance at 600nm was measured every 20 min after a 60 sec orbital shake (amplitude of 6mm) in a Tecan microplate reader (TECAN, Switzerland) in a method modified from ^61^. Bacterial growth curves represented by the OD_600_ plotted as a function of time were obtained, and data was processed using R software environment. The R package Growthcurver ^62^ was used to extrapolate the time when the bacterial culture reach a threshold of OD_600_ of 0.1 (T). Considering that T will be proportional to the initial bacterial content and given that bacteria show an exponential growth, we apply the qRT-PCR methodology to obtain the bacterial loads relative to CTL. ΔΔT values referred to CTL were calculated for each sample (ΔΔT_sample 1_ =T_sample 1_-T_CTL_) and fold change was calculated as 2^ΔΔTsample^ ^61^.

#### CFU calculations

Serial dilutions of bacterial culture were simultaneously (i) plated on LB agar plates, incubated overnight at 37°C and CFU counted and (ii) cultured in a 96-well plate and measured the OD_600_ over time using a microplate reader. Plots of T values versus CFUs gave a linear correlation between both parameters allowing to obtain the concentration of bacteria for a given T value.

