## Supplemental tables for "Defensive lipid droplets are PUFA reservoirs driving bacterial clearance and inflammation"

pmols of lipids identified in purified LDs from one liver

| | mLDs | | | | | | | | | | dLDs | | | | | | | | | |
|---|---|---|---|---|---|---|---|---|---|---|---|---|---|---|---|---|---|---|---|---|
| lipid species | #1 | #2 | #3 | #4 | #5 | #6 | #7 | #8 | #9 | #10 | #1 | #2 | #3 | #4 | #5 | #6 | #7 | #8 | #9 | #10 |
| CE 13:0;0 | 0 | 0 | 0 | 0 | 0 | 0 | 0 | 0 | 0 | 0 | 3270,87148 | 0 | 0 | 0 | 5212,0124 | 0 | 0 | 0 | 5244,94957 | 0 |
| CE 16:0;0 | 104138,441 | 68672,977 | 69884,3447 | 75911,9479 | 35499,1735 | 105914,656 | 63750,3719 | 95106,678 | 105870,934 | 146603,88 | 11598,2374 | 0 | 0 | 8327,36969 | 16324,2319 | 7584,15794 | 0 | 0 | 0 | 0 |
| CE 16:1;0 | 123297,024 | 68820,3176 | 81336,1549 | 73692,8495 | 28300,2904 | 144938,736 | 71046,7529 | 131119,08 | 139376,535 | 151025,066 | 10104,2639 | 11282,1402 | 13873,5326 | 5732,77283 | 12919,6368 | 4980,15118 | 5224,62006 | 16512,1047 | 19645,931 | 17803,2655 |
| CE 17:1;0 | 0 | 0 | 0 | 0 | 3234,52123 | 0 | 0 | 0 | 0 | 0 | 0 | 0 | 0 | 2483,02062 | 0 | 0 | 0 | 0 | 0 | 0 |
| CE 18:0;0 | 0 | 0 | 0 | 8405,82132 | 4008,28743 | 0 | 0 | 0 | 0 | 0 | 0 | 0 | 0 | 0 | 0 | 0 | 0 | 0 | 0 | 0 |
| CE 18:1;0 | 417078,819 | 347744,581 | 309332,555 | 344460,017 | 142630,157 | 461763,611 | 261714,878 | 459529,076 | 374505,539 | 584432,526 | 51547,287 | 44776,9274 | 61835,7658 | 38899,9863 | 68964,1698 | 32649,5132 | 13055,9006 | 58474,6106 | 61810,8413 | 59657,1409 |
| CE 18:2;0 | 251096,439 | 205382,707 | 0 | 191217,575 | 96035,0342 | 281942,266 | 180496,807 | 293463,612 | 266396,713 | 376642,342 | 0 | 0 | 0 | 0 | 0 | 0 | 0 | 0 | 0 | 0 |
| CE 18:3;0 | 0 | 12071,2757 | 0 | 10465,0346 | 5368,95943 | 19818,8496 | 0 | 17393,1229 | 0 | 0 | 0 | 0 | 0 | 0 | 0 | 0 | 0 | 0 | 0 | 0 |
| CE 20:3;0 | 0 | 0 | 0 | 0 | 0 | 0 | 0 | 0 | 0 | 0 | 0 | 0 | 0 | 0 | 0 | 2396,48008 | 0 | 0 | 0 | 0 |
| CE 20:4;0 | 24534,0729 | 14683,4109 | 19158,4826 | 12806,9353 | 6528,91986 | 21025,9199 | 13643,3578 | 20276,9589 | 0 | 25684,5284 | 12576,2126 | 14035,402 | 16294,028 | 9785,56506 | 16871,9196 | 8386,37257 | 4086,16066 | 16594,5133 | 14547,0002 | 16641,2618 |
| CE 22:6;0 | 24627,4567 | 11554,949 | 18714,2595 | 11702,4835 | 6501,18128 | 16988,136 | 0 | 0 | 0 | 28226,8095 | 6028,12131 | 0 | 12144,9232 | 4258,69401 | 9303,13322 | 5272,54581 | 0 | 11812,3531 | 12114,4384 | 0 |
| Cer 40:1;2 | 901,859999 | 0 | 0 | 0 | 0 | 0 | 0 | 0 | 980,139971 | 896,034986 | 0 | 0 | 361,983329 | 0 | 0 | 0 | 0 | 0 | 0 | 0 |
| Cer 42:1;2 | 1166,27254 | 0 | 0 | 378,515 | 180,664659 | 0 | 610,475019 | 928,69997 | 1312,51499 | 1034,96499 | 168,241998 | 0 | 606,918335 | 0 | 298,024449 | 0 | 0 | 474,003355 | 0 | 0 |
| Cer 42:2;2 | 1198,56998 | 0 | 0 | 389,646664 | 212,137341 | 741,299987 | 0 | 1063,49751 | 1499,56495 | 1403,28005 | 152,275999 | 0 | 0 | 0 | 286,488897 | 0 | 0 | 364,649991 | 0 | 0 |
| Chol | 746016,083 | 100919,325 | 260515,722 | 140602,926 | 54483,785 | 212551,359 | 0 | 376243,439 | 412651,825 | 501433,83 | 58901,5299 | 0 | 71320,2222 | 0 | 0 | 0 | 0 | 0 | 0 | 0 |
| DAG 14:0;0_18:1;0 | 2569,76992 | 0 | 0 | 0 | 0 | 2505,77331 | 0 | 2811,35499 | 6390,84518 | 6250,93222 | 0 | 0 | 0 | 0 | 0 | 0 | 0 | 0 | 0 | 0 |
| DAG 14:0;0_18:2;0 | 0 | 0 | 0 | 0 | 0 | 0 | 0 | 0 | 12523,0122 | 12103,045 | 0 | 0 | 0 | 0 | 0 | 0 | 0 | 0 | 0 | 0 |
| DAG 14:1;0_18:2;0 | 0 | 0 | 0 | 0 | 0 | 0 | 0 | 63648,5195 | 0 | 77804,594 | 0 | 0 | 0 | 0 | 0 | 0 | 0 | 0 | 0 | 0 |
| DAG 16:0;0_16:0;0 | 0 | 0 | 0 | 4986,90327 | 1989,35397 | 9943,24366 | 0 | 0 | 20491,302 | 21926,1527 | 0 | 0 | 0 | 0 | 0 | 0 | 0 | 0 | 0 | 0 |
| DAG 16:0;0_16:1;0 | 11561,563 | 0 | 0 | 0 | 0 | 10559,1035 | 0 | 8749,58277 | 25418,7322 | 21055,6197 | 0 | 0 | 0 | 0 | 0 | 0 | 0 | 0 | 0 | 0 |
| DAG 16:0;0_18:0;0 | 0 | 2541,05118 | 0 | 0 | 0 | 0 | 2718,4701 | 0 | 0 | 0 | 0 | 0 | 0 | 0 | 0 | 0 | 488,558006 | 0 | 0 | 0 |
| DAG 16:0;0_18:1;0 | 111851,034 | 25849,9781 | 38817,2722 | 46705,265 | 16859,8366 | 100061,582 | 39287,7555 | 76980,7005 | 191938,629 | 193661,976 | 4629,86724 | 9090,81868 | 10784,3685 | 3738,48216 | 7913,89783 | 3692,20757 | 4195,10078 | 9400,30495 | 6915,16596 | 12643,8772 |
| DAG 16:0;0_18:2;0 | 194270,916 | 40908,8347 | 74032,313 | 70083,9233 | 29057,9783 | 163647,257 | 67810,0586 | 138428,497 | 344558,449 | 344947,891 | 9332,43688 | 14062,6853 | 25551,3668 | 6355,35049 | 16153,5284 | 6492,99049 | 7199,57047 | 18805,6024 | 15074,8455 | 19893,099 |
| DAG 16:0;0_18:3;0 | 7086,89749 | 0 | 0 | 3182,92658 | 1194,07066 | 7435,58963 | 0 | 5557,6849 | 14908,3054 | 16488,4698 | 0 | 0 | 0 | 0 | 0 | 0 | 0 | 0 | 0 | 0 |
| DAG 16:0;0_20:1;0 | 0 | 0 | 0 | 0 | 0 | 0 | 0 | 0 | 0 | 2803,44486 | 0 | 0 | 0 | 0 | 0 | 0 | 0 | 0 | 0 | 0 |
| DAG 16:0;0_20:3;0 | 0 | 0 | 0 | 0 | 0 | 0 | 0 | 0 | 0 | 0 | 0 | 0 | 0 | 0 | 0 | 0 | 126,894796 | 0 | 0 | 0 |
| DAG 16:0;0_20:4;0 | 8994,24732 | 3972,93117 | 4974,13317 | 4080,53994 | 1615,05063 | 12250,5601 | 0 | 10579,5372 | 11324,2328 | 17557,7581 | 1151,58534 | 0 | 4550,77847 | 1138,11533 | 1890,86888 | 0 | 2418,50243 | 0 | 2482,82236 | 2035,68147 |
| DAG 16:0;0_22:4;0 | 0 | 0 | 0 | 0 | 530,446649 | 0 | 0 | 0 | 0 | 0 | 0 | 0 | 0 | 0 | 0 | 0 | 0 | 0 | 0 | 0 |
| DAG 16:0;0_22:5;0 | 3266,72494 | 5957,34225 | 0 | 1109,75504 | 709,545294 | 3011,41659 | 0 | 2457,82495 | 4963,65011 | 5635,9148 | 0 | 0 | 0 | 0 | 0 | 0 | 0 | 0 | 0 | 0 |
| DAG 16:0;0_22:6;0 | 23181,9558 | 5900,24418 | 8678,53324 | 8965,97147 | 3592,99056 | 18233,0529 | 8499,01021 | 15667,2204 | 29873,4355 | 42085,7048 | 0 | 0 | 7115,50474 | 0 | 3415,75676 | 1490,12601 | 1653,91235 | 0 | 2989,36591 | 4658,04467 |
| DAG 16:1;0_16:1;0 | 0 | 0 | 0 | 0 | 0 | 0 | 0 | 0 | 4859,1876 | 2995,40758 | 0 | 0 | 0 | 0 | 0 | 0 | 0 | 0 | 0 | 0 |
| DAG 16:1;0_18:1;0 | 22449,8391 | 3621,42457 | 0 | 7347,62351 | 2284,09672 | 20200,0173 | 5944,64779 | 17088,6302 | 46215,3482 | 42290,6923 | 0 | 1778,21721 | 1548,10667 | 0 | 0 | 0 | 1267,52558 | 1663,13837 | 0 | 2821,75082 |
| DAG 16:1;0_18:2;0 | 41105,1607 | 8920,20014 | 15640,6768 | 10830,4548 | 3880,874 | 31673,6666 | 15824,548 | 30190,537 | 80618,7534 | 72524,5667 | 0 | 4413,07 | 4558,33673 | 1470,96062 | 2983,3444 | 0 | 2433,24642 | 4471,14507 | 3087,46366 | 6460,75395 |
| DAG 16:1;0_18:3;0 | 5778,66495 | 0 | 0 | 0 | 0 | 3968,58017 | 0 | 4010,44488 | 4969,74498 | 7265,48016 | 0 | 0 | 0 | 0 | 0 | 0 | 0 | 0 | 0 | 0 |
| DAG 16:2;0_18:2;0 | 8910,53975 | 0 | 0 | 0 | 2553,33726 | 5097,73652 | 0 | 6567,08479 | 10089,798 | 14243,0472 | 0 | 0 | 0 | 0 | 0 | 0 | 0 | 0 | 0 | 0 |
| DAG 18:0;0_18:1;0 | 0 | 10033,1487 | 10151,9903 | 0 | 1898,29206 | 0 | 0 | 0 | 0 | 12605,5503 | 0 | 0 | 0 | 3379,29408 | 0 | 0 | 0 | 0 | 0 | 0 |
| DAG 18:0;0_18:2;0 | 6103,57225 | 1741,92442 | 2340,65334 | 3184,93505 | 1457,17732 | 5139,84005 | 2250,21496 | 4440,13745 | 10953,2881 | 15465,9772 | 362,623334 | 635,432856 | 2170,64341 | 368,440668 | 745,864444 | 341,08901 | 461,996794 | 934,168299 | 726,75179 | 770,62616 |
| DAG 18:0;0_20:4;0 | 26234,1595 | 14340,3753 | 21469,9427 | 6178,30833 | 2251,34261 | 15897,5331 | 18268,9178 | 14177,7027 | 12823,2098 | 16029,588 | 2926,92725 | 6389,66288 | 9600,4796 | 3167,8133 | 6335,96685 | 2065,61661 | 1972,26887 | 9122,5783 | 4800,45038 | 0 |
| DAG 18:1;0_18:1;0 | 70386,9581 | 17795,6751 | 25459,5868 | 34093,5262 | 12435,908 | 61124,1913 | 26325,0184 | 53167,82 | 127670,584 | 132602,787 | 2858,37523 | 5081,11273 | 5555,66986 | 2182,86928 | 4435,68892 | 2332,22508 | 2193,28003 | 4615,48686 | 3996,66702 | 8795,63845 |
| DAG 18:1;0_18:2;0 | 167478,924 | 42742,8309 | 55207,2779 | 72213,3446 | 31970,6853 | 143493,614 | 60082,8743 | 136930,637 | 321350,632 | 322349,701 | 6990,45563 | 14294,8709 | 19650,6802 | 6866,50658 | 14453,409 | 6603,47652 | 5902,94151 | 15240,6184 | 13457,2321 | 22543,6533 |
| DAG 18:1;0_18:3;0 | 22622,1132 | 6986,88454 | 10763,9337 | 7803,00697 | 3144,61263 | 17423,8237 | 10133,7898 | 16911,2873 | 33634,4886 | 34272,5158 | 1238,17603 | 3692,80713 | 3253,69 | 983,18998 | 2253,40552 | 1108,23202 | 764,32519 | 3465,4967 | 2113,66948 | 2718,11394 |
| DAG 18:1;0_20:3;0 | 3689,36747 | 0 | 0 | 2146,58161 | 1006,48602 | 4363,6167 | 0 | 3214,14739 | 5737,63013 | 5032,19008 | 927,71999 | 2434,75999 | 0 | 0 | 0 | 662,784517 | 617,934418 | 0 | 1355,66585 | 0 |
| DAG 18:1;0_20:4;0 | 8469,18762 | 0 | 0 | 3399,95344 | 2077,5733 | 7396,04314 | 0 | 8468,92238 | 14723,7623 | 17868,1028 | 0 | 0 | 0 | 0 | 0 | 0 | 0 | 0 | 0 | 0 |
| DAG 18:1;0_22:6;0 | 0 | 0 | 0 | 0 | 1432,57141 | 0 | 0 | 0 | 0 | 0 | 0 | 0 | 0 | 0 | 0 | 0 | 0 | 0 | 0 | 0 |
| DAG 18:2;0_18:2;0 | 92394,8097 | 29384,5495 | 41037,0445 | 33506,7622 | 16940,6904 | 71736,7045 | 47272,9683 | 73968,0767 | 160064,259 | 170301,952 | 5878,19799 | 13912,7132 | 18211,1629 | 5269,94928 | 11078,9214 | 5244,68994 | 2657,02724 | 17015,357 | 11683,0837 | 13268,7129 |
| DAG 18:2;0_18:3;0 | 19729,923 | 0 | 0 | 5995,00179 | 2712,60007 | 13762,366 | 0 | 13601,7478 | 31091,2895 | 30675,6473 | 0 | 0 | 0 | 0 | 0 | 0 | 0 | 0 | 0 | 0 |
| DAG 18:2;0_20:2;0 | 0 | 0 | 0 | 1198,30837 | 689,876 | 2942,54343 | 0 | 2931,25987 | 4605,40265 | 5385,83279 | 0 | 0 | 0 | 0 | 0 | 0 | 0 | 0 | 0 | 0 |
| DAG 18:2;0_20:3;0 | 2138,99255 | 0 | 0 | 850,589971 | 536,115328 | 1527,55668 | 0 | 2006,35999 | 4019,93006 | 4604,16764 | 0 | 0 | 0 | 0 | 0 | 0 | 0 | 0 | 0 | 0 |
| DAG 18:2;0_20:4;0 | 8460,36494 | 2327,31554 | 2994,40006 | 2538,63176 | 1858,53338 | 7325,90357 | 3835,00755 | 8744,99261 | 14989,63 | 17998,0969 | 0 | 0 | 0 | 0 | 0 | 0 | 515,709209 | 0 | 1081,88587 | 0 |
| DAG 18:2;0_22:5;0 | 0 | 0 | 0 | 0 | 582,575321 | 0 | 0 | 0 | 0 | 0 | 0 | 0 | 0 | 0 | 0 | 0 | 0 | 0 | 0 | 0 |
| DAG 18:2;0_22:6;0 | 0 | 0 | 0 | 0 | 2732,38468 | 0 | 0 | 0 | 0 | 0 | 0 | 0 | 0 | 0 | 0 | 0 | 0 | 0 | 0 | 0 |
| LPC 16:0;0 | 2120,49499 | 827,357769 | 1013,28999 | 1085,16167 | 398,266673 | 1236,76668 | 0 | 1331,60993 | 1965,87756 | 2114,90497 | 320,441326 | 382,722872 | 472,626686 | 188,944658 | 374,188887 | 149,382994 | 210,855198 | 1361,43833 | 0 | 400,495392 |
| LPC 18:2;0 | 0 | 0 | 0 | 0 | 203,396002 | 0 | 0 | 0 | 0 | 0 | 0 | 0 | 0 | 0 | 0 | 0 | 0 | 1092,06329 | 0 | 0 |
| LPE 16:0;0 | 413,774997 | 0 | 0 | 147,023338 | 91,4366643 | 0 | 0 | 308,155008 | 378,072485 | 376,180001 | 80,4866652 | 0 | 0 | 0 | 0 | 0 | 0 | 183,155003 | 0 | 0 |
| LPE 18:0;0 | 550,935008 | 0 | 0 | 176,59833 | 89,0806715 | 0 | 0 | 0 | 331,884995 | 406,609997 | 106,707335 | 0 | 0 | 0 | 118,492221 | 0 | 64,6340013 | 182,701672 | 0 | 0 |
| LPE O-16:1;0 | 0 | 0 | 0 | 0 | 0 | 0 | 240,800008 | 0 | 0 | 0 | 0 | 0 | 0 | 0 | 0 | 0 | 0 | 0 | 0 | 0 |
| LPG 16:0;0 | 0 | 0 | 0 | 0 | 0 | 0 | 0 | 0 | 0 | 0 | 0 | 0 | 0 | 47,3146687 | 0 | 0 | 0 | 0 | 0 | 0 |
| PC 16:0;0_16:0;0 | 0 | 0 | 0 | 0 | 0 | 0 | 0 | 0 | 0 | 0 | 527,17332 | 0 | 0 | 0 | 680,214432 | 0 | 275,796008 | 0 | 0 | 0 |
| PC 16:0;0_16:1;0 | 1432,22243 | 0 | 0 | 523,131688 | 0 | 850,646695 | 0 | 0 | 0 | 967,179984 | 326,202671 | 733,907138 | 338,351677 | 175,319334 | 366,277794 | 0 | 325,153995 | 639,553318 | 0 | 914,395406 |
| PC 16:0;0_18:1;0 | 14654,5279 | 6235,27315 | 9649,63675 | 7626,43655 | 3315,59467 | 10775,76 | 6563,45487 | 6263,24236 | 13555,7675 | 17222,6429 | 5808,10483 | 10584,9028 | 6445,93835 | 3375,7019 | 5511,24997 | 2390,2154 | 3356,92444 | 8781,6747 | 3762,24714 | 11475,8183 |
| PC 16:0;0_18:2;0 | 55773,5491 | 23970,9685 | 33584,0162 | 30200,4623 | 13637,989 | 39149,8343 | 27870,8482 | 47042,4032 | 60411,5438 | 63990,4785 | 18361,5634 | 33015,9296 | 29205,6656 | 11441,04 | 20449,7973 | 7861,54079 | 9573,10715 | 33734,3248 | 15194,0402 | 37535,0189 |
| PC 16:0;0_18:3;0 | 0 | 0 | 0 | 0 | 0 | 0 | 0 | 0 | 0 | 0 | 0 | 0 | 0 | 0 | 0 | 0 | 126,414394 | 0 | 0 | 0 |
| PC 16:0;0_20:3;0 | 1002,60749 | 368,251105 | 551,053335 | 522,889992 | 214,782675 | 552,349985 | 0 | 791,032463 | 807,982534 | 755,44998 | 415,363352 | 699,124294 | 449,884981 | 142,781337 | 296,574434 | 146,9035 | 424,281597 | 538,978328 | 294,788234 | 787,907655 |
| PC 16:0;0_20:4;0 | 26353,3568 | 11224,9332 | 17705,0734 | 13229,4599 | 6404,76926 | 17272,5169 | 11482,4724 | 19018,8599 | 22390,5253 | 28795,5189 | 5073,02411 | 9626,10585 | 6323,8732 | 3362,72526 | 5572,19558 | 2619,8051 | 4628,73688 | 9954,94684 | 4915,49436 | 11040,8658 |
| PC 16:0;0_22:5;0 | 3867,95759 | 0 | 0 | 2191,42159 | 1080,25201 | 0 | 0 | 0 | 0 | 4868,35986 | 1050,83402 | 2152,15138 | 1660,13002 | 876,879295 | 0 | 0 | 803,529644 | 2203,39 | 0 | 0 |
| PC 16:0;0_22:6;0 | 21303,5417 | 10318,487 | 14935,6937 | 10617,0019 | 4917,24809 | 13412,8332 | 10942,7321 | 15764,1554 | 18686,6903 | 24898,2835 | 6882,89134 | 10857,9513 | 9737,51148 | 3953,60851 | 6904,31542 | 3374,36104 | 3426,33667 | 11259,0265 | 6864,19655 | 11256,0265 |
| PC 16:1;0_18:1;0 | 1214,88497 | 0 | 0 | 0 | 0 | 0 | 0 | 0 | 1391,15751 | 0 | 0 | 752,028567 | 0 | 0 | 282,709996 | 0 | 418,351603 | 614,418338 | 0 | 1013,70463 |
| PC 16:1;0_18:2;0 | 0 | 0 | 0 | 0 | 0 | 0 | 0 | 0 | 0 | 0 | 0 | 0 | 0 | 0 | 0 | 0 | 601,37682 | 0 | 0 | 0 |
| PC 16:1;0_20:4;0 | 0 | 0 | 0 | 0 | 0 | 0 | 0 | 0 | 0 | 0 | 0 | 0 | 0 | 0 | 0 | 0 | 423,915195 | 0 | 0 | 0 |
| PC 18:0;0_18:1;0 | 0 | 0 | 0 | 0 | 0 | 0 | 0 | 0 | 0 | 0 | 722,409964 | 1159,79288 | 1275,41165 | 371,702671 | 762,177772 | 351,705015 | 335,309601 | 1259,42667 | 0 | 1315,47387 |
| PC 18:0;0_18:2;0 | 13470,7069 | 5312,66901 | 8772,91361 | 6740,33801 | 3521,24468 | 8817,83009 | 7115,39745 | 13009,9702 | 13696,6801 | 19993,0501 | 7052,75154 | 12030,7527 | 12347,5965 | 4441,1513 | 8671,56453 | 3709,59306 | 2114,19201 | 13542,455 | 5295,05898 | 12320,2911 |
| PC 18:0;0_20:3;0 | 0 | 0 | 0 | 0 | 0 | 0 | 0 | 0 | 0 | 0 | 0 | 0 | 0 | 0 | 0 | 0 | 320,608807 | 0 | 0 | 0 |
| PC 18:0;0_20:4;0 | 14265,3775 | 6165,13093 | 9499,32337 | 7280,78683 | 3944,55147 | 9431,57355 | 7528,62275 | 10335,927 | 13176,0502 | 20233,9721 | 4619,82187 | 7863,61422 | 6725,55129 | 3147,39672 | 5124,92233 | 2464,7696 | 2188,83209 | 8582,58168 | 4360,08005 | 7558,45877 |
| PC 18:0;0_22:6;0 | 4991,87261 | 2667,35554 | 3925,25315 | 2951,26994 | 1614,78202 | 4004,16334 | 2473,33243 | 3957,99011 | 5080,55985 | 7849,63489 | 3244,03731 | 4301,67573 | 5012,30995 | 1765,172 | 3166,77888 | 1761,747 | 997,105598 | 5182,46849 | 2785,32589 | 4105,69081 |
| PC 18:1;0_18:1;0 | 1333,25502 | 483,882229 | 869,629979 | 597,356657 | 234,94333 | 568,010012 | 489,427485 | 993,089974 | 747,004971 | 1338,69752 | 471,106013 | 955,957174 | 398,031672 | 304,638664 | 342,307786 | 189,953506 | 346,742797 | 752,441684 | 161,957653 | 1014,24465 |
| PC 18:1;0_18:2;0 | 6973,99259 | 2980,62007 | 4297,00017 | 4163,24496 | 1801,87861 | 5179,53992 | 4725,88241 | 9192,52276 | 9417,28234 | 10968,1356 | 2217,67728 | 4593,00143 | 3789,08674 | 1688,07205 | 2381,46226 | 1050,72451 | 1843,45913 | 4920,89152 | 1495,11758 | 5175,13385 |
| PC 18:1;0_20:3;0 | 0 | 0 | 0 | 0 | 0 | 0 | 0 | 0 | 0 | 0 | 0 | 0 | 0 | 0 | 0 | 0 | 160,570002 | 0 | 0 | 0 |
| PC 18:1;0_20:4;0 | 2271,39756 | 2743,60445 | 3979,15324 | 1224,32003 | 642,653982 | 5104,12335 | 3168,01757 | 5609,25007 | 6447,48986 | 3543,26248 | 483,096004 | 918,96568 | 250,418335 | 319,048663 | 1568,46338 | 698,208511 | 874,09277 | 787,749986 | 1417,15765 | 2945,06623 |
| PC 18:1;0_22:6;0 | 2868,36237 | 0 | 2187,29675 | 1591,11838 | 730,319977 | 0 | 1624,12256 | 0 | 0 | 0 | 952,493985 | 1706,43858 | 0 | 0 | 957,189997 | 0 | 867,012787 | 1488,08171 | 0 | 0 |
| PC 18:2;0_18:2;0 | 4337,48513 | 1760,06887 | 2344,59658 | 2203,04827 | 1094,36933 | 2950,01328 | 2296,7875 | 3218,26488 | 6546,81981 | 7401,81029 | 836,325328 | 2021,15434 | 1729,70335 | 750,79004 | 1299,69994 | 418,666989 | 830,121613 | 2132,76168 | 782,329405 | 2479,40614 |
| PC 18:2;0_20:4;0 | 2172,2725 | 0 | 1654,08333 | 1159,89496 | 610,655983 | 1398,43335 | 0 | 0 | 2013,47247 | 3002,52736 | 0 | 1089,8086 | 0 | 0 | 0 | 0 | 528,116417 | 0 | 0 | 0 |
| PE 16:0;0_18:1;0 | 0 | 0 | 0 | 0 | 0 | 0 | 0 | 898,937508 | 0 | 0 | 265,866001 | 660,857133 | 0 | 249,529998 | 337,546666 | 0 | 266,873193 | 522,753348 | 0 | 802,572324 |
| PE 16:0;0_18:2;0 | 9419,90972 | 3327,82216 | 5284,31018 | 4209,85659 | 1768,3533 | 6954,25034 | 4882,79492 | 8362,28728 | 10248,8446 | 10136,3778 | 2838,18531 | 5365,42586 | 4891,63518 | 1787,02005 | 3294,56568 | 1189,07595 | 1364,42003 | 5423,85658 | 2654,38108 | 6408,88287 |
| PE 16:0;0_20:3;0 | 0 | 0 | 0 | 0 | 0 | 0 | 0 | 0 | 0 | 0 | 0 | 144,592854 | 0 | 0 | 0 | 0 | 82,6860011 | 0 | 0 | 0 |
| PE 16:0;0_20:4;0 | 8247,10011 | 2650,54888 | 4851,20654 | 3219,95497 | 1557,00461 | 4947,34327 | 3190,265 | 6259,23753 | 7815,02485 | 8173,54977 | 1603,29803 | 3175,91565 | 2141,76993 | 998,549302 | 1687,64551 | 764,562011 | 1568,03236 | 3349,62845 | 1515,97177 | 3546,79071 |
| PE 16:0;0_22:5;0 | 1238,59502 | 308,575564 | 629,056642 | 375,301664 | 214,478672 | 589,50002 | 667,295009 | 896,285027 | 999,957472 | 1747,96745 | 336,026669 | 595,661444 | 257,920002 | 134,657999 | 170,227786 | 104,7865 | 176,261997 | 518,400023 | 0 | 648,536911 |
| PE 16:0;0_22:6;0 | 31251,6403 | 9870,09366 | 17248,826 | 12974,515 | 5406,01985 | 16834,4164 | 11863,2948 | 17632,4522 | 23743,1574 | 32320,7164 | 9029,77308 | 15823,1218 | 12361,7482 | 4728,68188 | 8195,13427 | 4049,65019 | 4018,62297 | 13945,4254 | 7196,74615 | 16982,906 |
| PE 16:1;0_20:4;0 | 0 | 0 | 0 | 0 | 0 | 0 | 0 | 0 | 0 | 0 | 0 | 0 | 0 | 0 | 0 | 0 | 192,630005 | 0 | 0 | 0 |
| PE 16:1;0_22:6;0 | 0 | 0 | 0 | 0 | 0 | 0 | 0 | 0 | 0 | 0 | 0 | 0 | 0 | 0 | 0 | 0 | 512,849188 | 0 | 0 | 0 |
| PE 18:0;0_18:1;0 | 0 | 0 | 0 | 0 | 0 | 0 | 0 | 0 | 0 | 0 | 261,89067 | 436,189984 | 0 | 223,997335 | 0 | 0 | 0 | 0 | 0 | 0 |
| PE 18:0;0_18:2;0 | 8304,48985 | 2197,6822 | 4549,88996 | 3286,74992 | 1552,15136 | 4251,39983 | 3989,06738 | 6726,24767 | 8081,85279 | 9357,51021 | 2489,72861 | 4535,38418 | 4519,67518 | 1698,05861 | 2984,63901 | 1047,95897 | 778,345203 | 5008,20518 | 1803,6517 | 5132,62162 |
| PE 18:0;0_18:3;0 | 0 | 0 | 0 | 0 | 0 | 0 | 0 | 0 | 0 | 0 | 0 | 184,447148 | 0 | 0 | 0 | 0 | 0 | 0 | 0 | 0 |
| PE 18:0;0_20:3;0 | 0 | 0 | 0 | 0 | 0 | 0 | 0 | 0 | 0 | 0 | 276,32467 | 512,564267 | 0 | 0 | 0 | 0 | 200,833201 | 552,406659 | 0 | 605,199979 |
| PE 18:0;0_20:4;0 | 17684,4728 | 6097,5043 | 9799,33977 | 7137,6737 | 3658,46411 | 8765,5735 | 6843,37735 | 11444,0501 | 14916,0349 | 18222,4655 | 4269,01277 | 7478,29437 | 6613,64834 | 2855,33396 | 4857,98094 | 2148,19789 | 2418,77766 | 8054,70705 | 3272,6142 | 7623,4326 |
| PE 18:0;0_22:5;0 | 0 | 0 | 0 | 0 | 256,673992 | 0 | 0 | 0 | 0 | 1350,49 | 289,575338 | 501,608551 | 0 | 0 | 0 | 0 | 181,367195 | 639,328311 | 0 | 571,933847 |
| PE 18:0;0_22:6;0 | 12108,4344 | 4181,9689 | 7255,71314 | 5698,03516 | 2520,25604 | 6085,30005 | 4916,28259 | 6690,31024 | 10321,3024 | 13315,0947 | 4066,49081 | 6747,93175 | 6438,1667 | 2226,9934 | 3980,49434 | 1823,16804 | 1101,10159 | 7216,92483 | 3210,27307 | 6564,68905 |
| PE 18:1;0_18:1;0 | 371,754989 | 0 | 133,783333 | 76,053335 | 29,2226672 | 147,506669 | 0 | 463,124998 | 271,834992 | 258,495007 | 70,091332 | 169,055717 | 128,046672 | 75,1426667 | 88,9999999 | 32,6845013 | 109,095204 | 174,408332 | 0 | 289,530777 |
| PE 18:1;0_18:2;0 | 5954,15771 | 1461,24893 | 3111,83333 | 2351,11336 | 1131,84071 | 3575,49985 | 2943,43501 | 6251,22011 | 7350,00014 | 7451,14028 | 1070,59137 | 2197,86423 | 1719,27492 | 958,402634 | 1414,33451 | 482,831001 | 862,403202 | 2710,09326 | 687,896504 | 3581,7293 |
| PE 18:1;0_20:3;0 | 0 | 0 | 0 | 0 | 0 | 0 | 0 | 0 | 0 | 0 | 0 | 0 | 0 | 0 | 0 | 0 | 61,0391974 | 0 | 0 | 0 |
| PE 18:1;0_20:4;0 | 6767,78495 | 1543,82891 | 3454,78018 | 2801,06505 | 1375,096 | 3783,04998 | 2607,74761 | 5554,23737 | 7347,34476 | 8494,81225 | 949,208657 | 2101,88287 | 883,733332 | 659,316659 | 880,444447 | 366,219014 | 1728,10116 | 1866,83993 | 0 | 2505,05851 |
| PE 18:1;0_22:5;0 | 0 | 0 | 0 | 0 | 0 | 0 | 0 | 0 | 0 | 0 | 0 | 0 | 0 | 0 | 0 | 0 | 171,9944 | 0 | 0 | 0 |
| PE 18:1;0_22:6;0 | 7162,64486 | 2645,58209 | 4475,70324 | 3649,59995 | 1488,07732 | 4396,58324 | 2894,57262 | 5109,43472 | 6636,2524 | 8273,50497 | 1450,49397 | 2969,45299 | 1700,94828 | 896,733999 | 1355,16445 | 652,727485 | 1196,29841 | 2660,9234 | 991,270612 | 3402,89373 |
| PE 18:2;0_18:2;0 | 1456,62993 | 475,742221 | 817,866673 | 565,909992 | 357,905984 | 910,506646 | 817,504972 | 1127,06251 | 2199,09996 | 2119,33255 | 296,854675 | 755,345736 | 557,608356 | 253,937324 | 356,046657 | 156,600505 | 354,616809 | 757,446686 | 247,878818 | 1123,75076 |
| PE O-22:0;0/16:0;0 | 0 | 0 | 0 | 0 | 0 | 0 | 0 | 0 | 0 | 0 | 0 | 0 | 0 | 718,072653 | 0 | 0 | 0 | 0 | 0 | 0 |
| PI 16:0;0_18:2;0 | 0 | 0 | 0 | 0 | 0 | 0 | 0 | 0 | 0 | 0 | 0 | 553,092871 | 0 | 0 | 0 | 0 | 258,304811 | 0 | 0 | 768,21075 |
| PI 16:0;0_20:4;0 | 3323,02511 | 1231,68005 | 2296,98658 | 1492,50666 | 883,877357 | 2542,29327 | 1803,79003 | 2626,89739 | 2976,00001 | 3587,75765 | 964,079301 | 2031,83293 | 1249,76496 | 672,626654 | 1085,41224 | 456,573993 | 894,061184 | 1872,74337 | 808,149436 | 2455,77225 |
| PI 18:0;0_18:2;0 | 1094,90499 | 0 | 0 | 385,388335 | 0 | 0 | 0 | 0 | 1340,93747 | 1376,97995 | 388,827324 | 890,851447 | 814,338326 | 341,718674 | 416,610009 | 0 | 176,001203 | 1121,7317 | 0 | 991,689242 |
| PI 18:0;0_20:3;0 | 0 | 0 | 0 | 0 | 0 | 0 | 0 | 0 | 0 | 0 | 375,371337 | 628,202856 | 0 | 169,435998 | 0 | 0 | 209,857988 | 751,456668 | 0 | 685,269237 |
| PI 18:0;0_20:4;0 | 41732,2254 | 16969,3555 | 27242,1487 | 20347,1422 | 9774,09681 | 25885,9762 | 21759,1882 | 30163,7769 | 40726,4996 | 51363,4157 | 14121,1103 | 23957,7947 | 17783,289 | 8849,88721 | 14550,0925 | 6763,29851 | 6152,08969 | 23228,3481 | 11040,692 | 26844,0217 |
| PI 18:1;0_18:1;0 | 0 | 0 | 0 | 0 | 0 | 0 | 0 | 0 | 0 | 0 | 0 | 0 | 0 | 0 | 0 | 0 | 19,7180003 | 0 | 0 | 0 |
| PI 18:1;0_18:2;0 | 0 | 0 | 0 | 0 | 0 | 0 | 0 | 878,290012 | 0 | 0 | 0 | 0 | 0 | 0 | 0 | 0 | 296,957588 | 0 | 0 | 0 |
| PI 18:1;0_20:4;0 | 2133,86744 | 734,533336 | 1193,56662 | 1082,33502 | 586,30534 | 1595,03003 | 1340,64004 | 2070,68995 | 2744,05003 | 2757,67505 | 340,219339 | 905,205693 | 0 | 0 | 0 | 0 | 1032,52802 | 875,910024 | 0 | 1185,89227 |
| PI 18:2;0_18:2;0 | 0 | 0 | 0 | 0 | 0 | 0 | 0 | 0 | 438,974984 | 0 | 0 | 0 | 0 | 0 | 0 | 0 | 60,3507996 | 0 | 0 | 0 |
| PS 18:0;0_22:6;0 | 0 | 0 | 0 | 0 | 0 | 0 | 0 | 0 | 0 | 0 | 0 | 3382,7843 | 4543,17848 | 0 | 0 | 0 | 0 | 0 | 0 | 0 |
| PS 18:1;0_18:1;0 | 0 | 0 | 0 | 0 | 0 | 0 | 0 | 0 | 0 | 0 | 0 | 0 | 292,226672 | 0 | 289,111104 | 0 | 104,983997 | 328,298335 | 0 | 0 |
| PS 18:2;0_18:2;0 | 0 | 0 | 0 | 0 | 205,612004 | 0 | 0 | 0 | 0 | 0 | 281,795343 | 746,484314 | 0 | 181,443334 | 306,232207 | 0 | 252,378011 | 0 | 0 | 0 |
| SM 36:2;3 | 0 | 0 | 0 | 0 | 1880,03937 | 0 | 0 | 0 | 0 | 0 | 1795,25407 | 0 | 0 | 2203,9566 | 0 | 1787,06002 | 1505,29604 | 0 | 0 | 0 |
| SM 41:3;2 | 4555,61757 | 994,637807 | 2813,36327 | 1620,01004 | 0 | 1838,35665 | 0 | 2387,54496 | 3001,83505 | 2759,90009 | 534,18533 | 968,401432 | 914,223293 | 0 | 670,594441 | 0 | 257,407188 | 0 | 0 | 733,584624 |
| SM 43:0;2 | 0 | 0 | 0 | 0 | 0 | 3124,06997 | 1513,89003 | 3792,79763 | 0 | 5534,80983 | 0 | 0 | 0 | 0 | 0 | 0 | 408,921194 | 0 | 727,254152 | 0 |
| TAG 42:2;0 | 0 | 0 | 0 | 0 | 0 | 0 | 0 | 0 | 8179,30758 | 8377,01261 | 0 | 0 | 0 | 0 | 0 | 0 | 735,399199 | 0 | 0 | 0 |
| TAG 44:1;0 | 11031,7695 | 0 | 5945,66981 | 0 | 0 | 5032,0967 | 0 | 9170,44759 | 16075,9699 | 12725,6322 | 0 | 0 | 0 | 0 | 0 | 0 | 672,718811 | 3416,61493 | 0 | 2745,55701 |
| TAG 44:2;0 | 20850,8253 | 0 | 0 | 0 | 0 | 10963,0863 | 0 | 20033,5145 | 33016,4099 | 28478,9109 | 0 | 0 | 0 | 0 | 0 | 0 | 0 | 7205,90512 | 0 | 0 |
| TAG 44:3;0 | 0 | 0 | 0 | 0 | 0 | 0 | 0 | 13720,1905 | 23225,4767 | 21790,4115 | 0 | 0 | 3809,97817 | 0 | 2443,56553 | 0 | 900,865555 | 0 | 0 | 3195,97244 |
| TAG 46:0;0 | 9990,21471 | 0 | 5705,37011 | 0 | 0 | 0 | 0 | 7662,56988 | 15632,2598 | 11950,773 | 0 | 2286,45563 | 0 | 0 | 0 | 0 | 0 | 3987,83843 | 0 | 2779,55697 |
| TAG 46:1;0 | 46197,0186 | 9364,8688 | 26739,0156 | 9666,26326 | 2786,15856 | 24625,0741 | 16940,918 | 40602,0975 | 69065,671 | 58929,0047 | 4831,81318 | 10550,7857 | 8594,92302 | 2170,54192 | 7233,27213 | 2038,29145 | 2748,77682 | 14889,4326 | 4512,41297 | 13253,9148 |
| TAG 46:2;0 | 88326,8261 | 17384,2292 | 49962,6923 | 18314,8797 | 6188,0525 | 49596,6689 | 36745,6436 | 84798,4982 | 138506,136 | 126280,375 | 8548,18598 | 20246,8545 | 23182,5701 | 5217,63929 | 16504,3788 | 4692,76333 | 5855,64194 | 36306,7722 | 9669,82561 | 28240,4533 |
| TAG 46:3;0 | 56448,1497 | 10737,133 | 30913,8425 | 11276,9103 | 3243,88536 | 33073,7813 | 24790,4992 | 57475,0853 | 92953,7201 | 77701,6687 | 4705,98602 | 10956,177 | 13666,5185 | 2842,17453 | 9467,31567 | 2629,72307 | 3736,59134 | 22265,7156 | 6287,81767 | 17271,3294 |
| TAG 46:4;0 | 21493,1703 | 7517,40244 | 14762,8562 | 4910,48495 | 1917,71539 | 13773,373 | 12222,9695 | 23944,6759 | 33899,6243 | 28584,3229 | 2191,33997 | 4937,78127 | 7010,40824 | 1620,23735 | 4379,61446 | 1513,58545 | 1651,94511 | 10085,9054 | 3153,31179 | 6360,6629 |
| TAG 46:5;0 | 0 | 0 | 0 | 0 | 0 | 0 | 0 | 0 | 11233,1271 | 8597,55754 | 0 | 0 | 0 | 0 | 0 | 0 | 0 | 0 | 0 | 0 |
| TAG 47:1;0 | 0 | 0 | 0 | 0 | 0 | 0 | 0 | 0 | 12001,4095 | 9375,31471 | 0 | 0 | 0 | 0 | 0 | 0 | 0 | 0 | 0 | 0 |
| TAG 47:2;0 | 9459,15222 | 0 | 0 | 0 | 0 | 5785,67664 | 5024,37472 | 11024,1079 | 18472,842 | 15349,462 | 0 | 0 | 0 | 0 | 0 | 0 | 729,206419 | 3825,1698 | 0 | 3409,08931 |
| TAG 47:3;0 | 0 | 0 | 0 | 0 | 0 | 0 | 0 | 9227,11253 | 11799,3855 | 12523,3805 | 0 | 0 | 0 | 0 | 0 | 0 | 733,782387 | 3168,4651 | 0 | 0 |
| TAG 48:0;0 | 36308,1026 | 9287,63602 | 22464,9874 | 11465,737 | 4692,85997 | 20151,4069 | 14312,1576 | 25828,2089 | 46491,3273 | 41710,8393 | 4122,00451 | 6847,66701 | 7489,85529 | 2128,72203 | 6648,44884 | 2321,63501 | 2309,81197 | 10495,7398 | 4434,63185 | 9374,13362 |
| TAG 48:1;0 | 232742,92 | 41166,4963 | 121661,568 | 48645,846 | 13080,8741 | 117147,84 | 74224,2289 | 160233,784 | 291098,824 | 254479,351 | 21573,5779 | 49795,7339 | 38275,4803 | 8777,34184 | 32938,8873 | 10380,908 | 14992,215 | 62458,299 | 23535,697 | 63729,8877 |
| TAG 48:2;0 | 414580,421 | 77589,3317 | 226038,589 | 89384,6957 | 24573,7457 | 232705,739 | 155928,23 | 345904,16 | 602667,465 | 514152,832 | 42986,1196 | 102426,583 | 93155,6575 | 20926,0025 | 72789,3829 | 21652,0119 | 34247,1924 | 150708,796 | 45853,7023 | 139856,638 |
| TAG 48:3;0 | 274766,846 | 55803,0319 | 147314,072 | 59086,3355 | 17113,9946 | 162563,324 | 119001,293 | 265937,271 | 440223,007 | 374733,543 | 25973,9278 | 62614,8932 | 67464,4979 | 15253,9787 | 47802,7005 | 15025,157 | 23329,8935 | 110805,753 | 31361,8649 | 91543,0803 |
| TAG 48:4;0 | 115125,713 | 26458,7339 | 63930,3271 | 28000,4946 | 9134,55518 | 67104,6829 | 57308,2542 | 129367,638 | 209733,238 | 175308,99 | 10757,0648 | 24723,4235 | 32886,4288 | 7438,82561 | 21649,3119 | 6298,08044 | 8000,0351 | 54584,9482 | 14276,849 | 38204,9737 |
| TAG 48:5;0 | 30500,0925 | 6481,58868 | 18884,8972 | 7232,81304 | 2309,27865 | 19190,5228 | 16028,527 | 36136,57 | 59401,7696 | 47884,7408 | 2511,37861 | 6336,62156 | 8620,37659 | 1850,588 | 5587,91002 | 1464,3625 | 2135,15968 | 12959,9818 | 3541,39889 | 9057,4602 |
| TAG 48:6;0 | 0 | 0 | 0 | 0 | 0 | 0 | 0 | 8074,46003 | 13739,6502 | 11404,3772 | 0 | 0 | 0 | 0 | 0 | 0 | 0 | 0 | 0 | 0 |
| TAG 49:1;0 | 20843,1482 | 4450,15324 | 11059,893 | 5450,91152 | 1654,97001 | 10523,8899 | 10102,1755 | 16811,1348 | 31113,1215 | 28923,614 | 2113,90607 | 4418,0928 | 4366,98834 | 1183,93135 | 3081,86902 | 990,406513 | 1296,02842 | 5493,565 | 2167,3483 | 5820,51094 |
| TAG 49:2;0 | 48828,6304 | 10412,1621 | 25055,1971 | 11368,6069 | 3591,32195 | 28485,082 | 22962,3556 | 46021,9336 | 77251,3819 | 71792,984 | 4416,22384 | 9572,15173 | 11011,1753 | 2873,85082 | 7838,74353 | 2347,25142 | 3708,09555 | 16253,3283 | 5978,72229 | 14052,6801 |
| TAG 49:3;0 | 30683,0931 | 5943,32006 | 15674,7039 | 6525,36988 | 2057,00731 | 18866,8633 | 14326,6404 | 29199,9125 | 50859,1652 | 45244,3218 | 2904,45741 | 6748,72262 | 7049,82201 | 1875,0747 | 5110,49906 | 1631,59096 | 2960,90088 | 11521,7066 | 4521,63304 | 10273,5938 |
| TAG 49:4;0 | 15202,7631 | 4792,32205 | 8607,76027 | 3531,13015 | 1647,50862 | 9383,8803 | 7924,00002 | 15543,3524 | 21552,2957 | 20120,2822 | 1694,74602 | 3923,62424 | 5424,94496 | 1393,61127 | 3447,5112 | 1229,07948 | 1193,85557 | 6744,80677 | 2631,39893 | 4333,83538 |
| TAG 50:1;0 | 857302,78 | 196666,768 | 475459,29 | 245587,819 | 73866,9535 | 438069,356 | 295810,604 | 539415,703 | 947998,734 | 908720,779 | 90895,4773 | 168520,05 | 149254,761 | 40667,249 | 131440,625 | 48655,1933 | 45038,4888 | 198375,079 | 88486,6961 | 203398,649 |
| TAG 50:2;0 | 2665767,21 | 583606,432 | 1472363,99 | 681843,262 | 198839,05 | 1452732,14 | 978616,638 | 1789305,88 | 3277706,91 | 2918436,28 | 275391,724 | 570013,559 | 544057,515 | 124400,309 | 429536,404 | 143112,961 | 182819,019 | 734294,688 | 318107,91 | 703434,918 |
| TAG 50:3;0 | 2301690,22 | 443138,496 | 1189806,32 | 501526,54 | 125696,36 | 1291832,07 | 810708,16 | 1592173,61 | 2936919,56 | 2507885,74 | 209777,161 | 504324,69 | 454752,909 | 101112,64 | 346897,719 | 108303,612 | 203715,979 | 683101,044 | 291319,131 | 669822,575 |
| TAG 50:4;0 | 803209 | 177912,462 | 430796,102 | 197442,729 | 55131,5562 | 494431,763 | 332807,312 | 681608,658 | 1248659,36 | 1023607,03 | 73535,0952 | 175955,582 | 196533,254 | 42323,499 | 139002,541 | 42204,464 | 76619,1895 | 284452,515 | 108511,128 | 237476,971 |
| TAG 50:5;0 | 232039,776 | 55467,9574 | 132187,182 | 59009,1769 | 17245,2609 | 147366,257 | 105847,712 | 243947,048 | 403142,052 | 335871,62 | 19859,7972 | 45122,3864 | 59344,279 | 12081,646 | 38091,9351 | 11387,454 | 16966,6946 | 87768,6246 | 28393,8038 | 65896,3952 |
| TAG 50:6;0 | 61870,0695 | 15854,8027 | 38751,0745 | 16532,3957 | 4580,50474 | 40699,7331 | 30117,1517 | 67473,0062 | 110176,201 | 96473,7225 | 5492,34708 | 12919,6181 | 16782,3855 | 3478,82875 | 10569,1338 | 3415,84945 | 4217,15775 | 24227,3935 | 7197,12594 | 17086,547 |
| TAG 50:7;0 | 13281,9998 | 0 | 7602,35627 | 0 | 0 | 8502,05342 | 6046,29278 | 14821,3303 | 24175,4603 | 20437,5505 | 0 | 0 | 3414,21843 | 0 | 0 | 0 | 922,537613 | 6219,42997 | 0 | 3632,80296 |
| TAG 51:1;0 | 15197,1674 | 0 | 8536,72345 | 4465,37018 | 1664,23941 | 7489,1297 | 7192,28745 | 12454,7696 | 18837,9896 | 22251,1554 | 2023,68402 | 0 | 3488,52674 | 1328,00666 | 3046,56108 | 1035,63905 | 0 | 4555,00841 | 0 | 4026,86156 |
| TAG 51:2;0 | 86776,3615 | 21257,0911 | 48185,3167 | 25651,7855 | 7096,92446 | 51051,1112 | 35476,7036 | 66298,1272 | 115401,592 | 115979,576 | 9532,37724 | 18611,3616 | 18488,9364 | 5931,97823 | 14615,1808 | 5414,59751 | 5811,78169 | 23954,6426 | 12294,4248 | 22637,5991 |
| TAG 51:3;0 | 136951,284 | 32810,2239 | 77092,3042 | 35885,9793 | 10881,0501 | 83737,4179 | 63618,8412 | 113015,165 | 197730,694 | 186906,643 | 14359,1957 | 29480,0895 | 34398,0535 | 9736,33194 | 24059,6008 | 8741,38355 | 10626,6655 | 43285,7164 | 23176,8933 | 38083,546 |
| TAG 51:4;0 | 71988,6255 | 18243,4972 | 37750,5207 | 19141,0081 | 6360,08898 | 45666,221 | 33412,4827 | 65039,2389 | 106770,086 | 98198,8621 | 6834,53496 | 15203,8874 | 19334,5594 | 5065,82673 | 12985,1352 | 4276,65806 | 5750,38223 | 23776,083 | 14026,6598 | 19376,5303 |
| TAG 51:5;0 | 14578,495 | 0 | 7349,2233 | 0 | 0 | 8283,20662 | 0 | 14351,7947 | 23235,8122 | 18310,0379 | 0 | 0 | 0 | 0 | 0 | 0 | 0 | 4983,93496 | 2193,80112 | 0 |
| TAG 52:1;0 | 0 | 18096,9705 | 0 | 0 | 10382,8405 | 0 | 0 | 0 | 0 | 73198,1516 | 13576,0969 | 20199,9651 | 19800,3928 | 6389,81692 | 16764,0474 | 7055,86863 | 0 | 17054,294 | 7939,74708 | 18139,2963 |
| TAG 52:2;0 | 2736758,42 | 810943,739 | 1749635,82 | 885736,287 | 249784,871 | 1507839,76 | 1025301,28 | 1687311,4 | 2785181,58 | 2852654,42 | 336963,888 | 623978,01 | 554912,364 | 184991,679 | 485160,082 | 216223,45 | 156468,75 | 655149,333 | 386424,417 | 703690,279 |
| TAG 52:3;0 | 10944514,2 | 2413567,17 | 5658572,18 | 2842413,33 | 768826,986 | 5223519,69 | 3361843,87 | 6255403,44 | 11563682,9 | 11466093,8 | 993580,648 | 1800196,53 | 2025443,93 | 563877,97 | 1622851,7 | 650008,545 | 490907,373 | 2337726,03 | 1460530,36 | 2206350,47 |
| TAG 52:4;0 | 7671558,23 | 2046647 | 4276055,09 | 2212887,57 | 655286,092 | 4328883,87 | 3083952,64 | 5104848,02 | 9407339,48 | 8763322,14 | 752103,271 | 1424007,31 | 1947849,32 | 462223,958 | 1394939,64 | 519783,936 | 436853,174 | 2180304,77 | 1440641,52 | 1782632,77 |
| TAG 52:5;0 | 2296756,59 | 595057,373 | 1279262,8 | 651621,043 | 172801,575 | 1390928,34 | 931151,657 | 1689352,42 | 3023634,64 | 2639225,16 | 189624,98 | 421554,347 | 516558,431 | 109420,797 | 353976,101 | 121728,302 | 153027,344 | 627775,116 | 351753,073 | 515737,492 |
| TAG 52:6;0 | 546425,591 | 137771,216 | 316254,4 | 153871,638 | 40777,3844 | 340356,649 | 217070,58 | 443553,696 | 775568,466 | 651293,259 | 38099,3423 | 87709,3506 | 108943,291 | 22271,2657 | 75508,5246 | 24339,4279 | 34551,9653 | 146562,03 | 66166,9922 | 111174,645 |
| TAG 52:7;0 | 105190,201 | 27894,037 | 61096,9607 | 29765,1704 | 8458,81208 | 67031,8604 | 46383,0996 | 104140,568 | 171619,148 | 142616,615 | 8896,93387 | 19030,3448 | 24320,0239 | 5439,75321 | 15876,468 | 5296,09156 | 6713,0394 | 35462,8372 | 12925,7763 | 24844,7653 |
| TAG 52:8;0 | 17767,7727 | 5103,47578 | 10512,9266 | 5474,64013 | 1949,42602 | 11866,8731 | 9176,47004 | 20046,5631 | 30663,7764 | 27477,6673 | 2020,63338 | 4085,28703 | 6229,03983 | 1257,11799 | 3656,0856 | 1156,26752 | 1431,00681 | 8546,78869 | 2570,45409 | 5167,57525 |
| TAG 53:2;0 | 36269,8817 | 10115,26 | 19864,7738 | 13412,1577 | 4632,47871 | 21696,2767 | 16497,3521 | 29391,6559 | 45597,6629 | 53105,6309 | 3852,69928 | 7534,14563 | 7490,48869 | 3299,65591 | 6640,38552 | 2727,93794 | 2144,48242 | 9649,63357 | 5153,35981 | 9642,14472 |
| TAG 53:3;0 | 97383,194 | 27174,1676 | 60284,7226 | 33548,7811 | 11252,0561 | 56848,4179 | 49819,3169 | 85728,5118 | 129575,901 | 146605,005 | 10942,969 | 19674,6513 | 23425,1293 | 8770,32026 | 18471,027 | 7528,23496 | 6124,37363 | 28982,4613 | 16017,4168 | 25099,1029 |
| TAG 53:4;0 | 107253,437 | 30665,3404 | 64325,1928 | 34494,8959 | 10825,9748 | 69277,6362 | 52888,1836 | 89976,9592 | 147526,674 | 148872,414 | 11249,8436 | 22126,0834 | 28574,8545 | 9083,62071 | 19647,2041 | 8022,41516 | 8133,13293 | 32648,7255 | 18991,0058 | 28794,1184 |
| TAG 53:5;0 | 53822,6891 | 15792,7916 | 32331,3363 | 18208,2812 | 5691,2276 | 37220,0235 | 27057,507 | 48205,8239 | 84364,8338 | 79268,4603 | 5648,84377 | 11990,4995 | 15331,9979 | 4329,4061 | 10022,4908 | 3521,13342 | 4692,01393 | 20073,6602 | 11459,9138 | 16196,3947 |
| TAG 53:6;0 | 12477,9069 | 0 | 8220,1767 | 4020,77675 | 0 | 10156,103 | 7549,19767 | 14865,5796 | 26197,4621 | 30113,6518 | 0 | 2940,1442 | 4111,10322 | 0 | 0 | 0 | 1265,32364 | 5759,10846 | 2882,32719 | 3910,97986 |
| TAG 54:1;0 | 0 | 0 | 0 | 0 | 1483,53863 | 0 | 0 | 0 | 0 | 0 | 2082,40668 | 4144,97137 | 0 | 1383,4947 | 0 | 1036,82649 | 0 | 0 | 0 | 4158,47375 |
| TAG 54:2;0 | 138356,953 | 42946,3026 | 81839,8221 | 63489,1129 | 23748,4665 | 66813,6978 | 60644,4883 | 90577,3449 | 162753,735 | 204960,155 | 23776,0239 | 43466,1947 | 41899,2074 | 16430,2508 | 33953,8087 | 15199,4476 | 8192,23328 | 41181,1352 | 23207,559 | 49383,093 |
| TAG 54:3;0 | 1394986,72 | 420656,569 | 842394,765 | 555599,264 | 166282,328 | 739777,425 | 562033,806 | 958621,368 | 1607189,33 | 1738737,95 | 155313,283 | 293985,443 | 280652,847 | 103833,099 | 226774,462 | 102069,069 | 80384,9854 | 326902,796 | 167784,693 | 365179,608 |
| TAG 54:4;0 | 3664606,93 | 1133101,81 | 2207914,63 | 1441835,94 | 444189,575 | 2117220,46 | 1645116,73 | 2890021,36 | 4894065,86 | 4914118,04 | 388349,365 | 748035,54 | 801646,932 | 267657,389 | 609458,143 | 259684,57 | 223623,486 | 936114,502 | 488292,488 | 962771,747 |
| TAG 54:5;0 | 3906180,11 | 1331351,18 | 2374692,18 | 1583480,12 | 549578,288 | 2447790,93 | 2039264,37 | 3340127,87 | 5887589,72 | 5709254,15 | 440018,026 | 835025,809 | 1084465,33 | 316680,094 | 749881,931 | 298198,578 | 260745,703 | 1257430,83 | 703596,12 | 1097598,13 |
| TAG 54:6;0 | 2374995,73 | 827281,562 | 1464058,13 | 896767,069 | 334468,058 | 1544985,45 | 1292862,55 | 2047830,35 | 3635882,87 | 3518857,42 | 282744,161 | 527758,048 | 746695,913 | 194464,437 | 501656,799 | 182107,468 | 180513,208 | 870572,917 | 508193,862 | 683214,017 |
| TAG 54:7;0 | 769631,195 | 232201,843 | 453106,944 | 251942,571 | 77414,093 | 493871,002 | 349684,372 | 613883,171 | 1081551,67 | 1016052,17 | 83638,799 | 173411,56 | 214213,308 | 48437,7085 | 146184,167 | 50909,1873 | 67876,5442 | 259081,446 | 140839,35 | 211534,025 |
| TAG 54:8;0 | 137700,148 | 42506,0654 | 85888,2078 | 46246,0105 | 13995,9602 | 95127,4808 | 66213,8367 | 117638,521 | 207036,228 | 183237,419 | 15490,6578 | 30377,0556 | 39075,0154 | 9230,22715 | 27486,1442 | 8884,47094 | 13639,3555 | 49272,4737 | 24783,9804 | 39274,4504 |
| TAG 54:9;0 | 22202,3273 | 4786,25086 | 13193,6598 | 7192,45036 | 2117,39461 | 14214,7859 | 9797,33229 | 20553,4649 | 34393,7016 | 32988,1525 | 2031,6987 | 4006,56428 | 5233,75154 | 1249,21068 | 4260,99433 | 1443,10606 | 1540,89956 | 8030,74837 | 3387,78243 | 5794,83839 |
| TAG 55:3;0 | 17041,6129 | 5825,84911 | 11510,2935 | 8023,23182 | 3482,84276 | 11451,207 | 10192,5445 | 18482,7626 | 28009,1262 | 34892,6663 | 1705,44863 | 3362,37294 | 4130,80494 | 1844,10731 | 3266,86435 | 1269,73295 | 949,822807 | 4990,15013 | 0 | 4517,77678 |
| TAG 55:4;0 | 33851,8214 | 11692,1817 | 19374,0066 | 13824,5233 | 5642,72372 | 23190,1932 | 20702,1904 | 33669,6243 | 47721,7007 | 56555,2998 | 3616,48814 | 6074,03415 | 8258,48818 | 3421,74784 | 6572,55332 | 2579,47993 | 2045,26272 | 9965,66693 | 5381,77546 | 9242,14143 |
| TAG 55:5;0 | 23420,1097 | 7767,95758 | 15256,9962 | 10479,7482 | 4116,27134 | 15915,4272 | 15664,0327 | 24776,9594 | 37681,6463 | 42028,3031 | 2935,44324 | 5213,85704 | 6791,67191 | 2746,05052 | 4691,92346 | 2145,85447 | 1956,6412 | 8443,35318 | 4754,04459 | 7983,92589 |
| TAG 55:6;0 | 14406,8575 | 4859,77756 | 8747,99649 | 4991,77496 | 2178,81473 | 10787,727 | 8092,29732 | 16105,3956 | 22748,9471 | 26783,3185 | 2226,66995 | 4245,83435 | 4910,42654 | 1793,62996 | 3491,98659 | 1281,51596 | 1651,68285 | 6520,00984 | 3492,62939 | 5848,25552 |
| TAG 55:7;0 | 13709,0802 | 0 | 8497,42015 | 4132,70672 | 1633,11132 | 9328,69673 | 7340,37995 | 12279,8324 | 19937,88 | 25292,9807 | 2066,07469 | 4540,81706 | 5026,89679 | 1459,79929 | 3815,05224 | 0 | 1820,13721 | 6296,18009 | 3218,9142 | 5821,49689 |
| TAG 56:10;0 | 18821,8129 | 6810,25346 | 11748,6135 | 7644,31636 | 2912,76932 | 12224,2069 | 8792,75739 | 17953,3827 | 30727,9992 | 31012,0082 | 2563,35274 | 5190,2427 | 7399,02814 | 1736,70673 | 4694,75322 | 1802,441 | 2666,18767 | 9846,44016 | 4512,94703 | 6499,14742 |
| TAG 56:1;0 | 0 | 0 | 0 | 0 | 0 | 0 | 14669,075 | 0 | 0 | 0 | 0 | 0 | 0 | 0 | 0 | 0 | 0 | 0 | 0 | 0 |
| TAG 56:2;0 | 0 | 0 | 0 | 4482,11511 | 2428,68725 | 0 | 7133,1501 | 8707,84521 | 13245,7149 | 16427,232 | 2975,11323 | 5913,23308 | 4908,59826 | 2148,33927 | 3388,64671 | 1283,99348 | 0 | 4167,83492 | 2569,65525 | 5949,96306 |
| TAG 56:3;0 | 62920,9852 | 17320,9286 | 36129,7989 | 28550,6376 | 12908,4638 | 37306,6902 | 33765,3494 | 60464,4728 | 102206,86 | 107899,694 | 12910,3343 | 26639,4724 | 21163,1823 | 10111,2862 | 16097,1419 | 6893,84556 | 5451,23711 | 23898,9782 | 12755,2952 | 31295,3568 |
| TAG 56:4;0 | 133530,884 | 40715,5397 | 81015,3198 | 68004,4492 | 30037,5824 | 89321,963 | 86243,8583 | 150507,488 | 239537,926 | 254625,797 | 25749,6872 | 48991,4322 | 52272,7839 | 22731,7963 | 37006,518 | 15399,5628 | 13741,6916 | 59666,9451 | 30372,7453 | 66252,8111 |
| TAG 56:5;0 | 254429,989 | 79687,9154 | 158094,915 | 105838,362 | 47787,0178 | 157570,356 | 129990,74 | 239198,99 | 351225,433 | 390893,555 | 54171,9106 | 91273,0517 | 96345,8697 | 39507,1284 | 71693,3441 | 32392,4904 | 31231,5033 | 108050,181 | 64544,1886 | 120132,998 |
| TAG 56:6;0 | 512349,319 | 175724,216 | 330532,608 | 196621,539 | 85120,0816 | 321147,766 | 259713,898 | 458805,618 | 663648,376 | 740469,742 | 103620,382 | 176755,927 | 173208,262 | 67877,5787 | 130445,574 | 57622,6158 | 65785,022 | 216656,621 | 131605,566 | 222351,685 |
| TAG 56:7;0 | 1018976,29 | 379325,697 | 683097,534 | 400641,861 | 160980,916 | 660968,018 | 541555,061 | 852210,083 | 1352999,11 | 1559031,37 | 197006,877 | 328737,575 | 358881,887 | 123958,669 | 266393,907 | 116920,486 | 121716,833 | 457690,379 | 254659,639 | 414347,159 |
| TAG 56:8;0 | 785479,431 | 309385,579 | 535618,54 | 316409,887 | 145518,127 | 533918,457 | 438299,904 | 626109,276 | 1069795,23 | 1159410,78 | 163788,94 | 270035,008 | 377336,578 | 117593,933 | 267319,421 | 110939,369 | 118694,36 | 445086,161 | 267210,245 | 352706,909 |
| TAG 56:9;0 | 165250,549 | 59756,0586 | 107453,601 | 65996,7995 | 24671,9666 | 118803,927 | 88017,7784 | 136560,583 | 234467,926 | 237171,574 | 29790,2374 | 54947,7659 | 73942,7503 | 19847,187 | 48307,9402 | 18720,4609 | 29481,311 | 88105,8884 | 49660,1015 | 73627,5189 |
| TAG 57:7;0 | 0 | 0 | 0 | 0 | 1398,62665 | 0 | 0 | 0 | 0 | 0 | 1561,64598 | 0 | 0 | 1194,78997 | 0 | 0 | 973,492432 | 0 | 0 | 0 |
| TAG 57:8;0 | 0 | 0 | 0 | 0 | 0 | 0 | 0 | 0 | 0 | 0 | 0 | 0 | 0 | 0 | 0 | 0 | 943,408775 | 0 | 0 | 0 |
| TAG 58:10;0 | 134376,659 | 70585,7086 | 100188,548 | 76191,1964 | 42931,9509 | 104229,787 | 98903,2269 | 130106,144 | 204928,169 | 226868,954 | 37442,716 | 60471,4857 | 95016,0662 | 30357,5083 | 59660,6361 | 25597,4426 | 28535,5408 | 105339,521 | 62990,3547 | 85508,0062 |
| TAG 58:11;0 | 25843,246 | 12958,5711 | 20332,9833 | 13769,9922 | 6132,80042 | 20334,5998 | 16274,7431 | 23285,5201 | 39548,6379 | 43327,6653 | 6310,99319 | 11294,6088 | 16982,5919 | 4576,93672 | 10553,0792 | 4441,8335 | 5249,62578 | 18505,1568 | 10834,9419 | 14286,254 |
| TAG 58:3;0 | 0 | 0 | 0 | 0 | 1813,84738 | 0 | 0 | 0 | 12879,0295 | 11859,6697 | 1695,86929 | 3430,76297 | 0 | 1441,94206 | 0 | 0 | 0 | 0 | 0 | 4185,70152 |
| TAG 58:4;0 | 12829,0355 | 4924,44674 | 8927,67986 | 7211,58187 | 3149,43409 | 8330,76 | 9311,64503 | 14886,4603 | 22789,0444 | 22963,3284 | 3336,02206 | 6485,8532 | 6225,56329 | 2872,66668 | 4543,31769 | 1901,08347 | 1600,21992 | 6791,08699 | 4144,454 | 8439,19974 |
| TAG 58:5;0 | 16851,4872 | 5907,35541 | 10081,95 | 8336,53688 | 4419,84876 | 10314,7697 | 9954,52523 | 18763,4671 | 25849,4711 | 26715,8055 | 6429,54254 | 11743,0769 | 10824,0652 | 5413,7338 | 7979,86348 | 3435,44841 | 3026,57642 | 12367,3948 | 6465,75759 | 14765,9874 |
| TAG 58:6;0 | 34621,7108 | 13827,0177 | 22649,0402 | 16576,4713 | 9495,91192 | 25747,4597 | 20922,9445 | 39731,5621 | 48797,2212 | 59874,4392 | 19727,4793 | 31808,9431 | 27775,631 | 13031,7548 | 19612,1513 | 8544,77692 | 9112,27264 | 32191,1271 | 16879,6551 | 41363,4139 |
| TAG 58:7;0 | 92782,2685 | 39249,4583 | 62521,0826 | 47357,7277 | 25860,8678 | 65817,0191 | 59394,7983 | 109210,787 | 139519,92 | 170283,146 | 38465,8915 | 66136,4365 | 64858,799 | 26362,3072 | 45012,5631 | 20033,3614 | 21083,6075 | 77628,0403 | 39965,0304 | 84171,6121 |
| TAG 58:8;0 | 180245,228 | 79624,0913 | 125440,114 | 95173,2763 | 51126,2004 | 131718,801 | 114502,115 | 191743,984 | 247799,53 | 292789,745 | 55288,8082 | 92271,5105 | 100496,527 | 41456,9906 | 73978,7716 | 33429,4281 | 38018,7744 | 125706,749 | 66150,3063 | 123814,885 |
| TAG 58:9;0 | 219865,665 | 104675,751 | 151462,555 | 123617,071 | 69282,9539 | 170691,757 | 151654,568 | 213260,517 | 312044,048 | 355898,857 | 61033,8898 | 101515,677 | 139072,126 | 48386,7493 | 91671,2104 | 42474,5445 | 48311,1725 | 161972,377 | 88480,6106 | 145184,655 |
| TAG 60:10;0 | 15845,238 | 8967,24171 | 12587,7897 | 7822,75359 | 5340,36191 | 13970,7963 | 12759,6331 | 20038,9385 | 21592,6051 | 28575,4848 | 8880,73222 | 13643,6149 | 15130,9109 | 6304,30984 | 10701,0534 | 4488,43813 | 6154,66194 | 17877,1257 | 10140,2496 | 17809,2047 |
| TAG 60:11;0 | 19184,0148 | 9713,94645 | 13108,6032 | 8962,18856 | 6091,82803 | 15719,4233 | 13764,137 | 19622,0028 | 22933,7525 | 34612,3767 | 7765,85007 | 12231,3023 | 12052,6687 | 5300,88075 | 9623,69177 | 4229,60997 | 5944,80515 | 17103,9041 | 9327,36565 | 14200,8018 |
| TAG 60:12;0 | 20050,1919 | 11804,8848 | 15549,0669 | 12145,1132 | 7139,59885 | 15403,2071 | 14386,6503 | 16224,0076 | 24088,0084 | 35611,1455 | 8921,49226 | 12845,2369 | 18367,7212 | 6525,96792 | 12313,8523 | 5797,34993 | 6534,56421 | 21611,956 | 11699,8852 | 15754,8611 |
| TAG 60:6;0 | 0 | 0 | 0 | 0 | 0 | 0 | 0 | 0 | 0 | 0 | 1899,43536 | 3415,60704 | 0 | 1552,67731 | 0 | 0 | 952,077198 | 0 | 0 | 4455,40758 |
| TAG 60:7;0 | 0 | 0 | 0 | 0 | 1896,77207 | 0 | 0 | 0 | 0 | 0 | 5761,91902 | 10026,7056 | 8885,45036 | 4099,45933 | 5558,92785 | 2472,09144 | 2669,40956 | 10641,1584 | 4689,72823 | 14500,91 |
| TAG 60:8;0 | 0 | 0 | 0 | 5304,71841 | 3970,4806 | 0 | 6461,39741 | 13610,5371 | 0 | 19804,2428 | 10013,3298 | 16471,7783 | 17094,5183 | 7317,61297 | 10430,0266 | 4905,9906 | 4885,20584 | 17945,9365 | 8536,06841 | 24053,3726 |
| TAG 60:9;0 | 12200,6929 | 5598,89581 | 7821,95965 | 6709,80136 | 4824,33669 | 9284,33657 | 9229,67494 | 16032,6624 | 16557,9796 | 22167,9425 | 9236,45528 | 15928,3079 | 17276,0153 | 7114,68506 | 10299,0511 | 4657,45783 | 5554,58641 | 19247,2665 | 10409,1173 | 20073,0016 |
| TAG 62:13;0 | 0 | 0 | 0 | 0 | 1484,95404 | 0 | 0 | 0 | 0 | 0 | 0 | 0 | 0 | 1331,08997 | 0 | 1074,47898 | 1378,8764 | 0 | 0 | 0 |
| TAG 62:14;0 | 0 | 0 | 0 | 0 | 0 | 0 | 0 | 0 | 0 | 0 | 0 | 0 | 0 | 1209,17336 | 0 | 0 | 1080,06325 | 0 | 0 | 0 |
| total pmol lipid | 52524724,8 | 14655339,2 | 29618491,2 | 16864714,3 | 5535253,3 | 30301094,7 | 21407569,1 | 38063644,3 | 65975249,1 | 64596942,4 | 5794495,9 | 10833163 | 12516329 | 3566673,47 | 9273054,4 | 3614737,81 | 3514731,3 | 14994508,6 | 8349240,46 | 13673314,2 |

**nmols of FAs identified in purified LDs from one liver**

| | mLDs | | | | | | | | | | dLDs | | | | | | | | | |
|---|---|---|---|---|---|---|---|---|---|---|---|---|---|---|---|---|---|---|---|---|
| FA | #1 | #2 | #3 | #4 | #5 | #6 | #7 | #8 | #9 | #10 | #1 | #2 | #3 | #4 | #5 | #6 | #7 | #8 | #9 | #10 |
| 12:0;0 | 243,922782 | 58,851294 | 140,94436 | 68,4620561 | 23,1017658 | 143,694518 | 120,579456 | 251,772003 | 424,470728 | 405,252785 | 24,6594073 | 53,1843552 | 68,0320321 | 16,8550409 | 47,768928 | 14,8034872 | 12,3180371 | 108,282886 | 28,5317508 | 71,220349 |
| 12:1;0 | 159,803016 | 31,1356959 | 91,9474619 | 19,7400294 | 7,55051459 | 73,0858541 | 69,8683445 | 171,763423 | 290,181066 | 246,186762 | 11,2215834 | 25,3037214 | 32,1839457 | 5,59747746 | 20,4669269 | 4,84354564 | 8,84530694 | 41,9884171 | 14,2381689 | 40,4464669 |
| 13:0;0 | 13,9363045 | 3,87822711 | 7,80968327 | 3,27526908 | 2,22651491 | 8,5567656 | 8,85819935 | 24,1952104 | 46,7279342 | 47,8682169 | 3,59185711 | 3,1293616 | 4,87697103 | 1,29465138 | 8,89799499 | 1,15672167 | 1,32490265 | 8,81034244 | 7,79335266 | 7,59907995 |
| 13:1;0 | 6,68650689 | 0 | 5,33663945 | 0 | 0,20714611 | 4,43429195 | 4,71091191 | 20,077102 | 18,8498393 | 21,5251418 | 0 | 1,75492696 | 0,58024527 | 0,68024461 | 0,31994721 | 0,72742406 | 1,08724601 | 9,62767107 | 0 | 2,28574538 |
| 14:0;0 | 1345,10292 | 303,244553 | 739,952077 | 350,0763 | 108,706374 | 758,865347 | 581,345979 | 1173,67317 | 2006,74455 | 1843,11768 | 163,811668 | 343,44692 | 359,649518 | 90,7945456 | 273,486074 | 88,7954687 | 94,5507921 | 530,914166 | 192,29781 | 456,816456 |
| 14:1;0 | 561,273489 | 128,116737 | 331,362503 | 138,166042 | 44,1501341 | 348,103987 | 278,123117 | 732,767249 | 991,484916 | 973,469209 | 52,7648948 | 118,81256 | 158,985701 | 37,2226956 | 105,050487 | 33,0908267 | 41,5491887 | 249,176739 | 67,175236 | 191,399972 |
| 15:0;0 | 218,342013 | 50,2698521 | 111,16861 | 58,3417691 | 19,497504 | 127,19434 | 111,664832 | 204,301837 | 336,040258 | 333,483565 | 22,1523698 | 41,8531309 | 56,344398 | 15,8348114 | 38,3913543 | 13,2165296 | 14,9332317 | 70,3224655 | 37,0493445 | 60,4281921 |
| 15:1;0 | 43,5454889 | 8,77041822 | 32,1994026 | 10,2120791 | 3,66259212 | 33,8130323 | 15,8004796 | 44,282565 | 67,0175426 | 61,7617766 | 5,2247737 | 8,51088275 | 10,6575372 | 3,29862894 | 7,86082423 | 2,40439441 | 4,06559725 | 21,0642129 | 9,49666242 | 13,8088658 |
| 16:0;0 | 33314,3871 | 8284,93533 | 18500,477 | 9258,92505 | 2787,14625 | 17751,802 | 12063,1081 | 21107,5766 | 38194,4101 | 36897,8147 | 3534,94619 | 6543,40253 | 7446,54759 | 1945,96935 | 5755,04078 | 2198,56091 | 1875,41092 | 8871,18904 | 5089,27883 | 8018,19381 |
| 16:1;0 | 9575,74761 | 1991,1859 | 5039,95943 | 2340,22557 | 565,501143 | 5759,44529 | 3503,36484 | 6897,84158 | 12709,3173 | 10739,0171 | 900,971885 | 2201,82102 | 1877,86609 | 467,995047 | 1413,83983 | 474,60325 | 1005,76341 | 2756,28684 | 1238,6647 | 2914,11853 |
| 16:2;0 | 904,268772 | 276,602347 | 530,39743 | 308,386547 | 102,179747 | 602,913717 | 440,246156 | 839,651385 | 1681,02843 | 1417,82731 | 88,4398159 | 191,171663 | 269,747987 | 60,2010582 | 181,221832 | 57,5249376 | 67,2112471 | 370,36781 | 124,463147 | 263,111838 |
| 17:0;0 | 170,80911 | 43,4255438 | 96,4980562 | 55,4052696 | 19,6601513 | 93,7155727 | 83,493821 | 143,319723 | 236,31576 | 263,080513 | 19,8484049 | 30,9991955 | 46,0031877 | 15,5656178 | 33,7212801 | 13,2904359 | 9,46551364 | 52,6665558 | 24,9408942 | 42,1452355 |
| 17:1;0 | 275,420288 | 69,6294642 | 163,447414 | 80,3038596 | 25,6963352 | 184,10147 | 124,512649 | 222,56312 | 401,579304 | 365,270744 | 30,7410671 | 66,6797216 | 67,825609 | 23,5652227 | 47,5589685 | 17,5658416 | 27,0556262 | 89,120834 | 47,9836669 | 85,3920334 |
| 17:2;0 | 46,4809115 | 10,0171408 | 27,5347317 | 14,1635379 | 3,78718355 | 26,4704239 | 18,7609764 | 34,044146 | 63,4304376 | 61,6406491 | 4,59311303 | 10,1594553 | 11,2061246 | 2,82699745 | 6,49101805 | 2,53970468 | 4,14438542 | 14,919158 | 7,74247448 | 13,8779097 |
| 18:0;0 | 1314,99965 | 439,602278 | 773,676775 | 531,763476 | 233,381948 | 615,174677 | 548,733455 | 824,969197 | 1562,17747 | 1836,62469 | 234,830844 | 378,723239 | 466,245654 | 140,124431 | 327,063811 | 134,446344 | 78,2837428 | 440,182908 | 228,712886 | 390,705673 |
| 18:1;0 | 40726,4793 | 11213,6984 | 23350,5029 | 13582,0315 | 4042,16984 | 22809,9448 | 15986,173 | 28678,7009 | 49440,9471 | 48930,743 | 4189,76937 | 8093,64698 | 8266,93254 | 2619,29888 | 6505,05017 | 2693,11952 | 2483,77091 | 10062,7863 | 5484,29064 | 10113,9091 |
| 18:2;0 | 46462,5252 | 13326,4624 | 25771,1649 | 15275,2985 | 5168,65552 | 26938,3532 | 20431,0695 | 34530,5782 | 62555,6222 | 59876,9702 | 4838,46581 | 9127,71015 | 12187,4399 | 3200,86166 | 8607,41824 | 3249,40354 | 2784,48347 | 14024,1244 | 8260,18611 | 11735,9068 |
| 18:3;0 | 5391,59346 | 1494,45946 | 3155,1078 | 1678,95677 | 460,545936 | 3213,79787 | 2099,1384 | 4209,94832 | 6653,35302 | 6132,82479 | 406,718868 | 861,838076 | 1034,42371 | 219,724693 | 740,075743 | 262,877435 | 258,405149 | 1201,66832 | 703,254883 | 982,317306 |
| 18:4;0 | 766,790517 | 176,989842 | 446,044663 | 214,627532 | 43,4790319 | 443,659508 | 239,541311 | 551,132124 | 848,192785 | 722,743251 | 23,5232473 | 58,0439406 | 69,9390938 | 10,7958486 | 53,4598132 | 18,0249414 | 17,9909113 | 82,4250089 | 48,8208344 | 61,215113 |
| 19:0;0 | 27,0433366 | 7,57306962 | 15,8581304 | 10,0876665 | 4,23452309 | 16,6362673 | 15,4824699 | 25,7575021 | 44,2110329 | 52,7164345 | 2,21189641 | 4,19290641 | 6,67974648 | 2,2379106 | 5,00522684 | 1,65115972 | 1,3804679 | 8,42642344 | 2,88134878 | 6,62474476 |
| 19:1;0 | 113,245853 | 35,501171 | 69,8383798 | 43,5315694 | 16,9662243 | 72,6856577 | 66,0780471 | 113,610092 | 167,380208 | 180,628006 | 11,7131361 | 20,5315442 | 25,947119 | 10,8899986 | 21,2491855 | 8,42502509 | 7,49483541 | 35,7929614 | 17,6054259 | 31,4019469 |
| 19:2;0 | 50,2164193 | 15,276807 | 27,1894267 | 17,6449684 | 5,92194131 | 28,5883067 | 25,2703364 | 47,98224 | 78,1437408 | 78,846565 | 4,82764365 | 8,15780925 | 9,90669677 | 3,09237485 | 7,77309699 | 3,15666028 | 2,96381121 | 12,3081085 | 6,42946195 | 11,8759812 |
| 19:3;0 | 11,7451071 | 3,0844476 | 8,92299134 | 4,33573712 | 1,33859541 | 9,4057791 | 8,7526018 | 12,8956697 | 18,3943007 | 20,3953797 | 1,29120282 | 2,63051378 | 3,33177903 | 0,72577116 | 2,32487223 | 0,58923129 | 1,05609115 | 4,01073813 | 2,39213732 | 4,02816631 |
| 19:4;0 | 4,60357285 | 1,15343192 | 4,84239827 | 2,13857955 | 0,75963366 | 4,54532687 | 2,10052734 | 5,28699475 | 7,37155785 | 10,305962 | 0,65532861 | 1,27647503 | 1,46221871 | 0,28867811 | 0,64149154 | 0,19042301 | 0,75935553 | 2,50464364 | 1,25811596 | 2,25312321 |
| 20:0;0 | 36,8221858 | 9,45764407 | 18,4495193 | 17,6315655 | 10,9034462 | 20,6465459 | 26,6909774 | 43,2290669 | 70,8778079 | 76,8950956 | 13,538495 | 28,5266882 | 24,5134644 | 9,80441646 | 16,2599398 | 5,92784639 | 3,1392927 | 22,4868733 | 14,4331248 | 28,4239775 |
| 20:1;0 | 425,008624 | 123,053283 | 248,986491 | 194,368017 | 86,4194663 | 286,404362 | 266,904081 | 463,578694 | 778,872671 | 809,220839 | 76,8637127 | 154,568179 | 163,185288 | 65,0978847 | 116,424146 | 48,5606704 | 41,6475497 | 196,613402 | 97,9399902 | 207,90936 |
| 20:2;0 | 234,816154 | 69,2843657 | 154,676462 | 97,0026366 | 42,0127297 | 152,413008 | 135,721022 | 277,311051 | 346,394983 | 406,65881 | 62,1601154 | 115,177421 | 114,297729 | 47,5128975 | 80,554328 | 33,03548 | 36,4624253 | 131,313606 | 68,4244109 | 151,539683 |
| 20:3;0 | 518,964777 | 167,97723 | 348,073711 | 206,492128 | 98,501812 | 349,200978 | 296,772732 | 558,568294 | 803,636323 | 830,102304 | 166,616114 | 291,148456 | 260,751833 | 98,7188338 | 170,178752 | 84,4451242 | 122,167054 | 334,997604 | 184,162595 | 368,675326 |
| 20:4;0 | 3418,78354 | 1341,34597 | 2512,08906 | 1251,01874 | 551,805519 | 2535,93127 | 1970,37963 | 3243,5292 | 5145,49876 | 5779,34741 | 588,926174 | 1049,01323 | 1065,18518 | 427,182319 | 811,367308 | 325,851273 | 487,069278 | 1466,84795 | 840,222768 | 1333,13262 |
| 20:5;0 | 546,626017 | 177,586809 | 362,418378 | 190,809533 | 66,831299 | 364,256196 | 231,975002 | 339,083004 | 621,448002 | 654,751202 | 81,1840265 | 139,895501 | 198,510493 | 54,7788839 | 137,367112 | 53,3456788 | 57,2368418 | 211,875549 | 139,87506 | 170,843393 |
| 21:0;0 | 0 | 0 | 0 | 0,11418906 | 0,0531862 | 0,23819349 | 0 | 1,16981291 | 2,27866381 | 1,96582253 | 0,07047711 | 0,1717688 | 0,18556357 | 0,0466461 | 0,11389874 | 0,03280001 | 0 | 0,24478145 | 0,08593789 | 0,24543759 |
| 21:1;0 | 2,19147727 | 0,64783333 | 1,37031658 | 0,81859143 | 0,23239888 | 1,86754319 | 0,88582031 | 2,03460988 | 2,74499359 | 3,66106099 | 0,32687879 | 0,5351061 | 0,68391217 | 0,20783425 | 0,41419289 | 0,14780229 | 0,11301356 | 0,92440434 | 0 | 0,67847307 |
| 21:2;0 | 1,05999585 | 0 | 0,50603497 | 0,36479995 | 0,15446932 | 0,89436537 | 0,41249948 | 0,9864329 | 1,74184664 | 1,87979581 | 0,21444254 | 0,41550813 | 0,39587414 | 0,05093843 | 0,35085703 | 0,05207353 | 0,19971353 | 0,47164683 | 0,15408687 | 0,63045544 |
| 21:3;0 | 2,37314234 | 0,68025291 | 1,65805224 | 0,8601238 | 0,59070317 | 1,65262353 | 0,53738302 | 2,00267946 | 4,36905029 | 4,12521568 | 0,73901793 | 0,91183155 | 0,76817647 | 0,38279185 | 0,64053989 | 0,18601295 | 0,58948105 | 1,29859855 | 0,6664862 | 1,51542243 |
| 21:4;0 | 2,10860199 | 0,95177134 | 1,75220677 | 0,80081614 | 0,54077396 | 1,69497637 | 1,32615611 | 1,77348384 | 2,25176824 | 3,00876221 | 0,71345646 | 0,74948842 | 0,92551615 | 0,44597242 | 0,65463744 | 0,27010639 | 0,58420858 | 1,06517833 | 0,74141218 | 1,16141927 |
| 21:5;0 | 5,72325198 | 0,64942556 | 0,90102076 | 0,65357108 | 0,83730952 | 1,2120226 | 0,62012506 | 0,98178669 | 10,1992229 | 10,2340475 | 0,85883413 | 2,98449597 | 0,46754537 | 1,30073081 | 0,35182057 | 0,12846622 | 1,6892841 | 0,56078716 | 0,36482259 | 0,65948848 |
| 21:6;0 | 0,66806161 | 0 | 0 | 0,18343166 | 0 | 0,36566956 | 0,24798921 | 0,3130173 | 1,37048225 | 0,50615215 | 0 | 0,13018725 | 0,17089672 | 0,10853566 | 0 | 0 | 0,12405791 | 0,17163265 | 0,12029797 | 0,17064772 |
| 22:0;0 | 1,5658605 | 0,49543578 | 0,87071477 | 1,17239736 | 1,14386954 | 1,05574595 | 4,42127434 | 3,41606784 | 9,66788962 | 9,1570317 | 0,91850374 | 1,79901164 | 1,43634184 | 0,68295685 | 0,85538195 | 0,25802268 | 0,09548106 | 1,05333088 | 0,72572459 | 1,96404635 |
| 22:1;0 | 17,0065984 | 4,99536294 | 10,6357979 | 8,92441361 | 5,17899829 | 9,78514106 | 51,967187 | 22,1004614 | 43,3907996 | 40,8017698 | 7,15816278 | 15,8450947 | 12,1105069 | 6,01105353 | 8,23366125 | 3,29207123 | 2,02127172 | 12,8761186 | 7,00974614 | 19,2614295 |
| 22:2;0 | 8,21134901 | 3,49438913 | 5,02838657 | 3,45099802 | 1,84227074 | 9,6386181 | 5,12914949 | 8,8272112 | 11,931398 | 12,1553102 | 3,06408054 | 5,89904743 | 6,03122162 | 2,58762341 | 3,92326622 | 1,4708218 | 1,41340603 | 5,86671823 | 2,82119542 | 8,16309626 |
| 22:3;0 | 17,0210013 | 7,10152117 | 12,3671887 | 7,87236257 | 3,72488479 | 11,5291611 | 11,0324546 | 18,7088037 | 20,7151348 | 25,7658953 | 7,01506272 | 12,2634173 | 10,1419927 | 4,43502513 | 7,25676211 | 3,49476043 | 4,03279952 | 11,897311 | 6,50031226 | 14,4058317 |
| 22:4;0 | 246,403012 | 102,061877 | 171,942728 | 107,795315 | 61,6616572 | 190,588611 | 158,109698 | 287,031061 | 341,748146 | 420,20681 | 89,1371782 | 143,410147 | 127,944593 | 63,2696554 | 100,939354 | 44,3843522 | 55,1781309 | 172,389956 | 107,747557 | 197,114017 |
| 22:5;0 | 798,178921 | 323,924727 | 545,712276 | 322,394366 | 164,843612 | 587,678572 | 493,837348 | 904,395685 | 1044,21509 | 1234,57158 | 168,134927 | 277,595774 | 304,060142 | 110,120622 | 234,903671 | 100,971039 | 118,230643 | 405,796915 | 254,201836 | 363,856848 |
| 22:6;0 | 4139,46686 | 1527,07889 | 2681,32087 | 1712,52421 | 750,844011 | 2562,73184 | 2035,71631 | 2916,86345 | 4855,63981 | 5622,6191 | 953,877981 | 1529,12313 | 1975,24579 | 624,207817 | 1407,8231 | 609,883282 | 625,894294 | 2330,10632 | 1238,8073 | 1975,04696 |
| 23:3;0 | 0,4391959 | 0,26143844 | 0,35410794 | 0,2710766 | 0,0902934 | 0,36948609 | 0,40888516 | 0,28810946 | 0,40945636 | 0,47638712 | 0,07121533 | 0,15342116 | 0,26452103 | 0,06892713 | 0,13090887 | 0,04386909 | 0,05073475 | 0,19924484 | 0 | 0,22808765 |
| 23:5;0 | 0,57800688 | 0 | 0 | 0 | 0,06374767 | 0 | 0,30671868 | 0,30216254 | 0,4825823 | 0,77470863 | 0,06336782 | 0,1196745 | 0,15798807 | 0 | 0 | 0,04909301 | 0,08615658 | 0,15960749 | 0 | 0 |
| 24:0;0 | 0 | 0 | 0 | 0 | 0,06028556 | 0 | 0 | 0 | 0,40494341 | 0,28778158 | 0 | 0 | 0 | 0 | 0 | 0 | 0 | 0 | 0 | 0 |
| 24:1;0 | 0,61301087 | 0 | 0 | 0 | 0,58812601 | 0 | 0,43123503 | 0,82518344 | 6,62305971 | 5,55409563 | 0,4446039 | 0,93086658 | 0 | 0,49521538 | 0 | 0 | 0 | 0 | 0 | 1,24999973 |
| 24:2;0 | 0,8928884 | 0,32724709 | 0,72184622 | 0,64805416 | 0,23628402 | 0,37064697 | 0,60494817 | 0,9981579 | 1,52645379 | 1,68896343 | 0,41943796 | 0,36609237 | 0,29102292 | 0,29309153 | 0,32661056 | 0,1167475 | 0 | 0,33971614 | 0,25757042 | 0,48059896 |
| 24:3;0 | 1,19211413 | 0,71633065 | 0,6020031 | 0,72725116 | 0,28319407 | 0,89215799 | 0,80031173 | 1,08190709 | 1,68216922 | 1,41333833 | 0,69817524 | 1,1694572 | 0,93060772 | 0,37064026 | 0,66698016 | 0,23914194 | 0,31844167 | 1,07979311 | 0,53648067 | 1,12068641 |
| 24:4;0 | 4,14786677 | 1,60214327 | 3,04843456 | 2,1785861 | 2,01605438 | 3,17543139 | 3,21153391 | 8,14466992 | 6,06821147 | 8,12426228 | 14,9940965 | 26,7606716 | 19,008177 | 9,8659631 | 11,0797587 | 4,60248281 | 6,29510436 | 25,0585035 | 10,3444481 | 39,20308 |
| 24:5;0 | 11,7545436 | 5,23064059 | 8,8878688 | 7,17700151 | 4,98629631 | 10,5959466 | 9,65098082 | 24,1314836 | 16,0283942 | 22,6296901 | 32,1979709 | 57,6588477 | 56,2983435 | 18,5302178 | 31,1908601 | 13,2269334 | 14,3731034 | 58,000869 | 26,2655136 | 77,0247898 |
| 24:6;0 | 20,0836982 | 7,02015382 | 14,1841421 | 9,79510001 | 6,21289559 | 15,6309183 | 14,375011 | 30,6104932 | 20,8152273 | 28,9461599 | 18,8110255 | 33,0246468 | 28,2199148 | 11,2074575 | 20,7873396 | 9,88135221 | 11,6062987 | 33,2317298 | 20,3943371 | 42,6809601 |

pmols of FAs identified in purified LDs from one liver

| type | name | carbons | mLDs #1 | mLDs #2 | mLDs #3 | mLDs #4 | mLDs #5 | dLDs #1 | dLDs #2 | dLDs #3 | dLDs #4 | dLDs #5 | dLDs #6 |
|---|---|---|---|---|---|---|---|---|---|---|---|---|---|
| SFA | Lauric ac | FA 12:0;0 | 160200,386 | 46740,73696 | 66570,40119 | 35368,47115 | 82114,68697 | 30225,52729 | 40468,30177 | 41458,69255 | 22848,21987 | 26813,11369 | 14862,67686 |
| | Myristic ac | FA 14:0;0 | 586420,2881 | 223265,152 | 465446,701 | 256567,1921 | 513960,3806 | 169655,0941 | 303195,4575 | 237135,3912 | 178831,7108 | 248491,0393 | 162749,5956 |
| | Palmitic ac | FA 16:0;0 | 11781940,92 | 5350438,232 | 9129392,7 | 4256661,682 | 9810830,078 | 2746341,248 | 4214866,028 | 3561752,319 | 1860665,131 | 3447910,767 | 2017264,252 |
| | Stearic ac | FA 18:0;0 | 1586175,995 | 1240860,596 | 1218152,313 | 1087827,377 | 1111213,913 | 1179556,885 | 1110859,451 | 1007834,091 | 989460,8307 | 931009,903 | 1254509,125 |
| | Arachidic ac | FA 20:0;0 | 75026,30234 | 33304,93927 | 33844,05136 | 30257,94506 | 38674,42131 | 39322,55507 | 52370,28599 | 35061,68365 | 33943,50052 | 32363,30986 | 28188,1237 |
| | Behenic ac | FA 22:0;0 | 31250,51498 | 12663,24878 | 10487,87594 | 6344,287395 | 12679,30865 | 23272,25924 | 10040,02571 | 9360,247254 | 9408,357143 | 7165,14647 | 11035,24089 |
| | Lignoceric ac | FA 24:0;0 | 90767,77458 | 94062,65259 | 8991,159201 | 6123,843193 | 11577,14963 | 94495,28694 | 19658,4487 | 6688,84635 | 5047,788024 | 5631,843209 | 11843,38927 |
| MUFA | Myristoleic ac | FA 14:1; n-5 | 85288,09547 | 36085,91318 | 55386,76739 | 28795,06826 | 94784,70802 | 13750,57578 | 36237,42342 | 23095,79849 | 8690,019846 | 23214,74075 | 11710,54244 |
| | Palmitoleic ac | FA 16:1;0 n-7 | 3731188,354 | 1285078,43 | 2875916,443 | 1231166,992 | 3526489,258 | 736214,447 | 1681475,83 | 1025600,51 | 563857,4219 | 1029926,91 | 506263,1989 |
| | Oleic ac | FA 18:1;0 n-9 | 16867727,05 | 8875170,898 | 14266385,5 | 8051586,914 | 15270955,81 | 3985313,416 | 6712282,715 | 5243043,213 | 3311994,019 | 5221291,504 | 2614895,02 |
| | 11-Eicosenoic ac | FA 20:1;0 n-9 | 229848,9189 | 119547,4243 | 176201,3817 | 131240,8638 | 225568,3327 | 88559,3605 | 148346,8437 | 108686,7142 | 93713,90343 | 96514,1201 | 52320,15133 |
| | Erucic ac | FA 22:1;0 n-9 | 17672,97268 | 13118,39223 | 13506,8965 | 12950,25468 | 15820,93596 | 12122,4308 | 21795,39442 | 14156,1079 | 15603,89042 | 12720,12591 | 13517,18545 |
| | Nervonic ac | FA 24:1;0 n-9 | 12176,97263 | 9238,874912 | 9195,722342 | 8524,366021 | 9794,286489 | 8997,113705 | 8839,588761 | 8599,51973 | 7707,467675 | 8048,992753 | 7646,666169 |
| PUFA w-6 | Linoleic ac | FA 18:2;0 n-6 | 19852812,5 | 9536057,739 | 14621594,24 | 8187677,002 | 16702358,4 | 4191174,316 | 6972745,972 | 7192832,031 | 3775338,745 | 6332954,712 | 2908920,593 |
| | Gamma-Linolenic ac | FA 18:3;0 n-6 | 995804,9774 | 467228,3936 | 745559,9976 | 419299,0875 | 917765,1215 | 95253,63922 | 184154,6249 | 165362,7396 | 71020,81299 | 175087,0514 | 75875,11063 |
| | Eicosadienoic ac | FA 20:2;0 n-6 | 63065,67669 | 34779,16479 | 45923,01369 | 33178,41291 | 54208,73165 | 35253,05986 | 53784,08432 | 43353,02353 | 35560,91309 | 37603,30439 | 18099,17092 |
| | Dihomo-g-linolenic ac | FA 20:3;0 n-6 | 142855,4153 | 78698,30608 | 117583,5323 | 76383,14247 | 138662,138 | 120622,1294 | 185341,301 | 132064,8003 | 90703,02963 | 107121,6297 | 55015,38754 |
| | Arachidonic ac | FA 20:4;0 n-6 | 917760,2386 | 539006,0806 | 768730,7739 | 383361,6638 | 863451,6907 | 258390,3122 | 430880,7373 | 346383,667 | 283268,4708 | 326349,7925 | 157892,2272 |
| | Docosatetraenoic ac | FA 22:4;0 n-6 | 132108,8314 | 96861,85837 | 111005,3444 | 76013,49354 | 144147,7299 | 91123,13271 | 127357,5878 | 96950,48332 | 100973,2723 | 94795,06493 | 54824,95785 |
| | Docosapentaenoic ac | FA 22:5;0 n-6 | 100311,7657 | 68156,79073 | 85627,25067 | 44110,43167 | 106169,7197 | 39047,32943 | 60931,51093 | 44628,14331 | 37875,73814 | 43329,28658 | 24685,77147 |
| PUFA w-3 | Alpha-Linolenic ac | FA 18:3;0 n-3 | 750000,1526 | 324003,9825 | 547836,1893 | 240994,7586 | 642627,9449 | 134838,5048 | 346651,535 | 284999,2943 | 107165,7848 | 225279,4456 | 98396,66367 |
| | Octadecatetraenoic ac | FA 18:4;0 n-3 | 216798,439 | 76533,71811 | 178061,2946 | 70882,41577 | 208874,5308 | 15962,28838 | 32604,57277 | 25682,90472 | 9525,867105 | 22075,14286 | 14164,36076 |
| | Eicosapentaenoic ac | FA 20:5;0 n-3 | 132046,175 | 55282,87888 | 104168,7965 | 54836,74526 | 108868,5226 | 32459,58567 | 59561,3718 | 61048,80333 | 32834,08165 | 49581,4991 | 23627,60544 |
| | Docosapentaenoic ac | FA 22:5;0 n-3 | 211845,0928 | 121436,3861 | 190170,536 | 113645,3724 | 203084,7168 | 93471,8895 | 123521,5187 | 130820,8466 | 70595,50285 | 106823,4825 | 51266,675 |
| | Docosahexaenoic ac | FA 22:6;0 n-3 | 912155,1514 | 534707,9849 | 779473,1903 | 462324,4095 | 806013,6414 | 387081,604 | 593905,9448 | 579839,4394 | 357826,5762 | 489443,7408 | 256749,7635 |
| | | total pmol FAs | 59683249,0 | 29272328,77 | 46625212,07 | 25306122,19 | 51620696,16 | 14622503,99 | 23531876,56 | 20426439,31 | 12074461,05 | 19101545,67 | 10446323,45 |

pmols of FA identified per ul mice serum

| | | | CTL | | | | | | 2h LPS | | | | | |
|---|---|---|---|---|---|---|---|---|---|---|---|---|---|---|
| | | | #1 | #2 | #3 | #4 | #5 | #6 | #1 | #2 | #3 | #4 | #5 | #6 |
| SFA | octanoic acid | FA 8:0 | 9,57 | 10,77 | 11,34 | 48,71 | 39,10 | 23,77 | 11,63 | 9,71 | 13,87 | 41,35 | 36,44 | 21,26 |
| | Decanoic acid | FA 10:0 | 1,00 | 1,01 | 0,77 | 1,90 | 1,77 | 1,50 | 2,11 | | 1,89 | 1,62 | 1,99 | 1,75 |
| | Undecanoic acid | FA 11:0 | 0,62 | 0,80 | 0,53 | 0,59 | 0,58 | 0,53 | 0,84 | 0,48 | 0,61 | 0,55 | 0,54 | 0,55 |
| | Lauric acid | FA 12:0 | 4,33 | 3,05 | 2,56 | 3,27 | 3,72 | 4,08 | 7,34 | 7,03 | 6,20 | 5,72 | 5,95 | 4,00 |
| | Tridecylic acid | FA 13:0 | | | | 0,17 | 0,24 | 0,13 | 0,07 | | | 0,30 | 0,24 | 0,25 |
| | Myristic acid | FA 14:0 | 25,07 | 20,46 | 22,42 | 14,74 | 17,46 | 21,81 | 34,83 | 28,69 | 36,06 | 27,64 | 28,92 | 18,47 |
| | Pentadecylic acid | FA 15:0 | 8,56 | 9,08 | 10,87 | 1,62 | 2,07 | 1,82 | 12,31 | 11,31 | 13,17 | 1,85 | 1,46 | 1,53 |
| | Palmitic acid | FA 16:0 | 1696,69 | 1633,94 | 1775,03 | 1387,61 | 1568,53 | 1566,35 | 1846,15 | 1744,84 | 1859,51 | 1611,65 | 1661,41 | 1509,02 |
| | Margaric acid | FA 17:0 | 16,06 | 18,87 | 20,32 | 19,73 | 22,40 | 23,42 | 20,38 | 19,09 | 20,22 | 21,88 | 17,01 | 19,28 |
| | Stearic acid | FA 18:0 | 509,82 | 523,09 | 514,53 | 476,72 | 535,93 | 551,41 | 510,13 | 465,73 | 549,05 | 505,74 | 522,60 | 509,69 |
| | Arachidic ac | FA 20:0 | 15,64 | 13,09 | 15,11 | 22,77 | 27,22 | 28,72 | 22,97 | 19,42 | 25,51 | 29,48 | 26,36 | 22,35 |
| | Behenic ac | FA 22:0 | | | 9,17 | 15,81 | 19,55 | 17,88 | 4,61 | 4,15 | 5,74 | 15,49 | 12,94 | 12,55 |
| | Lignoceric ac | FA 24:0 | 2,46 | 2,58 | 2,94 | 2,49 | 3,25 | 3,09 | 3,87 | 2,86 | 4,08 | 2,19 | 2,15 | 2,30 |
| MUFA | Myristoleic ac | FA 14:1;n-5 | 2,61 | 1,31 | 1,72 | 0,62 | 0,78 | 1,00 | 2,44 | 2,28 | 2,62 | 0,98 | 1,14 | 0,88 |
| | Palmitoleic ac | FA 16:1;n-7 | 388,09 | 286,28 | 268,97 | 180,20 | 178,48 | 275,87 | 441,06 | 339,89 | 469,83 | 344,46 | 408,40 | 269,32 |
| | Margaric acid | FA 17:1 n-7 | 16,25 | 18,40 | 18,76 | 10,96 | 11,47 | 16,41 | 24,81 | 19,73 | 21,85 | 19,31 | 15,64 | 14,29 |
| | Oleic ac | FA 18:1;n-9 | 1264,46 | 1209,01 | 1260,41 | 970,12 | 1277,86 | 1378,77 | 1576,21 | 1349,05 | 1514,07 | 1527,57 | 1620,29 | 1252,66 |
| | 11-Eicosenoic ac | FA 20:1;n-9 | 33,83 | 45,14 | 42,85 | 27,72 | 48,71 | 52,35 | 65,15 | 45,60 | 59,85 | 51,35 | 50,57 | 35,68 |
| | Erucic ac | FA 22:1;n-9 | 3,91 | 5,68 | 5,69 | 7,39 | 8,02 | 9,89 | 12,22 | 6,54 | 9,00 | 11,40 | 9,29 | 7,59 |
| | Nervonic ac | FA 24:1;0 n-9 | 1,44 | 1,46 | 1,52 | 1,64 | 2,20 | 2,31 | 2,89 | 2,43 | 3,22 | 2,31 | 1,96 | 2,13 |
| PUFA w-6 | Linoleic ac | FA 18:2;n-6 | 1891,96 | 2156,06 | 2292,91 | 2076,94 | 2573,94 | 2299,52 | 2013,89 | 2044,74 | 2127,00 | 2327,08 | 2195,13 | 2133,30 |
| | Gamma-Linolenic ac | FA 18:3;n-6 | 24,39 | 37,63 | 39,50 | 50,00 | 46,04 | 36,28 | 41,99 | 37,78 | 48,04 | 55,14 | 51,22 | 48,84 |
| | Eicosadienoic ac | FA 20:2;n-6 | 12,06 | 14,38 | 13,10 | 10,86 | 12,86 | 14,34 | 19,89 | 16,15 | 18,10 | 21,54 | 19,84 | 16,62 |
| | Dihomo-g-linolenic ac | FA 20:3;n-6 | 100,82 | 91,22 | 75,32 | 73,08 | 64,60 | 94,27 | 100,78 | 86,26 | 92,82 | 122,59 | 121,51 | 98,39 |
| | Arachidonic ac | FA 20:4;n-6 | 607,25 | 768,20 | 779,86 | 765,28 | 710,47 | 738,55 | 719,96 | 701,83 | 762,53 | 825,36 | 898,35 | 794,71 |
| | Docosapentaenoic acid | FA 22:2 n-6 | 0,51 | 0,85 | 0,88 | 0,85 | 0,79 | 0,82 | 1,81 | 1,32 | 1,67 | 1,42 | 1,14 | 0,96 |
| | Docosatetraenoic ac | FA 22:4;n-6 | 12,51 | 16,91 | 15,58 | 20,18 | 18,00 | 17,86 | 33,14 | 21,45 | 30,12 | 32,66 | 29,95 | 24,28 |
| | Docosapentaenoic ac | FA 22:5;n-6 | 30,46 | 31,14 | 29,56 | 33,97 | 34,89 | 30,55 | 39,70 | 30,85 | 32,94 | 44,79 | 52,57 | 39,03 |
| PUFA w-3 | Alpha-Linolenic ac | FA 18:3;n-3 | 27,79 | 35,79 | 42,43 | 41,36 | 93,56 | 72,15 | 38,85 | 43,72 | 45,12 | 64,94 | 53,01 | 50,25 |
| | Octadecatetraenoic ac | FA 18:4;n-3 | 2,00 | 2,01 | 2,62 | 2,02 | 2,34 | 1,44 | 4,40 | 4,20 | 4,71 | 3,36 | 1,91 | 3,09 |
| | Eicosapentaenoic ac | FA 20:5;n-3 | 26,92 | 25,82 | 23,36 | 19,53 | 19,87 | 23,33 | 31,17 | 23,87 | 36,30 | 25,69 | 21,36 | 23,47 |
| | Docosapentaenoic ac | FA 22:5;n-3 | 26,30 | 29,05 | 28,64 | 25,69 | 21,36 | 23,47 | 44,60 | 35,71 | 44,64 | 38,45 | 33,91 | 32,96 |
| | Docosahexaenoic ac | FA 22:6;n-3 | 243,03 | 289,32 | 223,29 | 268,39 | 295,12 | 280,22 | 243,13 | 248,12 | 263,20 | 332,23 | 352,33 | 309,98 |

pmols of FA identified in RAW macrophages

| | | treatment | CTL | | | | OA | | | |
|---|---|---|---|---|---|---|---|---|---|---|
| | | | #1 | #2 | #3 | #4 | #1 | #2 | #3 | #4 |
| SFA | octanoic acid | 8:00 | 160,924024 | 208,251557 | 239,38412 | 237,486572 | 69,6525281 | 99,5687537 | 216,324806 | 192,310039 |
| | Decanoic acid | 10:00 | 12,7185759 | 9,56301519 | 9,89607329 | 12,0871447 | 10,794047 | 6,54918986 | 8,56775857 | 7,11579312 |
| | Undecanoic acid | 11:00 | 3,05169282 | 2,25693345 | 2,09369332 | 2,17096097 | 2,61150451 | 2,16155631 | 2,30056863 | 1,96835981 |
| | Lauric acid | 12:00 | 24,1041325 | 18,7316931 | 18,9863997 | 20,2720989 | 18,5159958 | 17,874265 | 18,3629433 | 15,7610289 |
| | Tridecylic acid | 13:00 | 3,48540379 | 2,92520957 | 1,55638265 | 2,2154679 | 2,83314275 | 2,42453492 | 1,56683419 | 0,86831799 |
| | Myristic acid | 14:00 | 254,01036 | 321,41815 | 246,787172 | 266,960407 | 242,638989 | 365,710459 | 266,795885 | 208,704368 |
| | Pentadecylic acid | 15:00 | 7,19964407 | 7,81080355 | 6,22107035 | 6,64587522 | 6,26783139 | 9,44397917 | 6,39021873 | 5,32828069 |
| | Palmitic acid | 16:00 | 2031,55704 | 2285,81061 | 1779,12125 | 2473,67103 | 1510,9985 | 2403,89309 | 1720,97 | 1415,19712 |
| | Margaric acid | 17:00 | 45,5551375 | 52,6824288 | 40,8964684 | 44,6573497 | 42,9523174 | 65,3018099 | 45,1579287 | 40,012722 |
| | Stearic acid | 18:00 | 1528,4525 | 1714,95824 | 1444,3394 | 1781,0998 | 1357,40279 | 2112,94127 | 1552,41725 | 1219,3954 |
| | Arachidic ac | 20:00 | 17,7515218 | 19,4937283 | 16,7443301 | 22,9971744 | 10,7118176 | 19,7343355 | 14,7026343 | 12,1808257 |
| | Behenic ac | 22:00 | 23,4737662 | 23,0480547 | 21,6802789 | 24,1746081 | 12,2040175 | 24,7459347 | 13,5086153 | 11,2596824 |
| | Lignoceric ac | 24,00 | 16,9900678 | 17,4345454 | 14,874512 | 15,8674972 | 10,3556503 | 13,7119019 | 11,755314 | 8,67500216 |
| MUFA | Myristoleic ac | 14:1;n-5 | 4,62406819 | 5,07610762 | 3,98136599 | 4,25631307 | 3,47373892 | 4,27167929 | 3,94715618 | 3,17244125 |
| | Sapienic acid | 16:1;n-10 | 1333,35351 | 1747,30627 | 1438,41457 | 1353,55552 | 1051,24487 | 1667,56288 | 1548,47993 | 1075,93375 |
| | Palmitoleic ac | 16:1;n-7 | 1181,13171 | 1649,19769 | 1137,45983 | 1198,51163 | 1004,40909 | 1650,40222 | 1223,58485 | 943,03237 |
| | Margaric acid | 17:1 n-7 | 80,887269 | 105,050255 | 81,9381186 | 85,4256949 | 74,0851367 | 117,075749 | 82,4492147 | 66,658471 |
| | Oleic ac | 18:1;n-9 | 4567,63252 | 5760,44393 | 4761,44193 | 4621,35559 | 6511,28246 | 9104,28162 | 7325,5444 | 5958,08189 |
| | 11-Eicosenoic ac | 20:1;n-9 | 100,019337 | 131,34242 | 113,713551 | 106,286119 | 124,466572 | 210,698872 | 147,530205 | 117,435874 |
| | Erucic ac | 22:1;n-9 | 18,5864564 | 25,0842695 | 21,0018509 | 20,6475908 | 23,967968 | 38,930903 | 27,940334 | 22,4684722 |
| | Nervonic ac | 24:1;n-9 | 18,652588 | 22,0217099 | 20,5211987 | 19,5067368 | 17,0662047 | 26,1090657 | 20,2590994 | 15,1057552 |
| PUFA w-6 | Linoleic ac | 18:2;n-6 | 284,856395 | 340,331388 | 290,3047 | 305,369819 | 231,013485 | 360,662316 | 279,035406 | 215,686739 |
| | Gamma-Linolenic ac | 18:3;n-6 | 20,9076567 | 25,8252031 | 19,8740734 | 21,2841874 | 18,3654089 | 25,5670435 | 20,8334338 | 17,5010551 |
| | Eicosadienoic ac | 20:2;n-6 | 70,7065733 | 88,8356486 | 81,7487574 | 76,8676965 | 58,3789641 | 96,3826873 | 75,8646589 | 57,4564902 |
| | Dihomo-g-linolenic ac | 20:3;n-6 | 106,956535 | 135,360469 | 138,383763 | 137,150091 | 97,6684133 | 147,510438 | 138,9197 | 110,67186 |
| | Arachidonic ac | 20:4;n-6 | 290,315049 | 337,645949 | 370,984319 | 394,716823 | 326,838229 | 466,821583 | 431,186747 | 400,286799 |
| | Docosapentaenoic acid | 22:2 n-6 | 3,36252118 | 4,0045644 | 3,51228659 | 3,54415319 | 2,71737967 | 3,91353719 | 3,09267835 | 2,52873102 |
| | Docosatetraenoic ac | 22:4;n-6 | 34,3746394 | 39,1913691 | 48,6954219 | 52,6204855 | 43,4356362 | 57,9637022 | 62,6941779 | 60,2608467 |
| | Docosapentaenoic ac | 22:5;n-6 | 7,69024266 | 8,95319185 | 9,71069709 | 10,4448895 | 10,2281804 | 12,433706 | 10,4611769 | 10,3300748 |
| PUFA w-3 | Alpha-Linolenic ac | 18:3;n-3 | 2,02410999 | 1,82836794 | 2,80312592 | 2,82449186 | 1,03428109 | 1,6831549 | 1,36998594 | 0,63610616 |
| | Octadecatetraenoic ac | 18:4 n-3 | | | | | | | | |
| | eicosatetraenoic acid | 20:4;n-3 | 1,89406465 | 2,39174945 | 1,35384415 | 1,5298296 | 1,35293523 | 2,21817256 | 1,34330458 | 1,1592174 |
| | Eicosapentaenoic ac | 20:5;n-3 | 74,4699944 | 93,4803486 | 74,9906054 | 79,1568284 | 60,0954619 | 87,9064662 | 69,3904338 | 52,8767153 |
| | Docosapentaenoic ac | 22:5;n-3 | 187,662517 | 214,169779 | 233,285951 | 253,310753 | 213,070132 | 296,494746 | 284,213223 | 250,033027 |
| | Docosahexaenoic ac | 22:6;n-3 | 193,67226 | 236,743978 | 230,770423 | 248,464647 | 207,842952 | 283,354909 | 267,925243 | 228,802121 |

| | | treatment | FA | | | | dMIX OA | | | |
|---|---|---|---|---|---|---|---|---|---|---|
| | | | #1 | #2 | #3 | #4 | #1 | #2 | #3 | #4 |
| SFA | octanoic acid | 8:00 | 177,610802 | 166,947932 | 220,87966 | 217,005339 | 170,225728 | 162,698903 | 198,625319 | 244,919862 |
| | Decanoic acid | 10:00 | 6,32566307 | 7,83519706 | 10,4988806 | 11,0620028 | 6,5431655 | 7,42776956 | 8,00962784 | 12,3598154 |
| | Undecanoic acid | 11:00 | 2,05520443 | 2,30639004 | 2,45052958 | 2,23495037 | 2,17538992 | 2,18414869 | 2,37069918 | 2,09519929 |
| | Lauric acid | 12:00 | 16,509635 | 20,2028995 | 18,6557699 | 18,5126956 | 19,9371996 | 20,9600678 | 18,8419157 | 22,0159187 |
| | Tridecylic acid | 13:00 | 1,48796751 | 2,22869911 | 2,09181644 | 2,63508361 | 1,87175628 | 2,02260767 | 1,39435489 | 2,14064345 |
| | Myristic acid | 14:00 | 254,307228 | 371,464757 | 259,280833 | 198,042493 | 306,46611 | 371,151057 | 260,945946 | 225,56177 |
| | Pentadecylic acid | 15:00 | 6,62649356 | 9,00291405 | 5,92218545 | 5,16384497 | 7,97566401 | 9,52064329 | 6,18470951 | 5,97094429 |
| | Palmitic acid | 16:00 | 1779,03443 | 2686,80102 | 1828,97038 | 1462,17481 | 1841,15847 | 2189,19376 | 1558,82384 | 1406,6203 |
| | Margaric acid | 17:00 | 45,883057 | 63,5024514 | 45,5446685 | 36,39013 | 53,2381655 | 61,5510893 | 43,8964373 | 40,7356618 |
| | Stearic acid | 18:00 | 1425,6018 | 2122,28852 | 1505,73518 | 1109,39069 | 1566,72824 | 1849,22005 | 1338,79727 | 1190,83557 |
| | Arachidic ac | 20:00 | 11,8276949 | 19,9156011 | 13,8542769 | 12,6221386 | 13,7704139 | 17,8533092 | 13,3292223 | 12,8242736 |
| | Behenic ac | 22:00 | 18,7813916 | 26,8948634 | 19,8480368 | 17,3372728 | 15,6918335 | 25,3775274 | 15,629608 | 13,2394192 |
| | Lignoceric ac | 24,00 | 10,7599172 | 18,1968525 | 12,0029379 | 9,5178849 | 10,5280106 | 13,7744299 | 11,324052 | 8,42958119 |
| MUFA | Myristoleic ac | 14:1;n-5 | 3,78586547 | 4,40806694 | 4,02828596 | 3,46909462 | 4,23664949 | 4,71112145 | 3,81225864 | 4,0092902 |
| | Sapienic acid | 16:1;n-10 | 1064,3322 | 1610,75275 | 1401,36919 | 934,142587 | 1152,86498 | 1480,13342 | 1381,43991 | 1065,41686 |
| | Palmitoleic ac | 16:1;n-7 | 1038,24129 | 1631,98143 | 1134,78003 | 825,713008 | 1245,34084 | 1557,47295 | 1186,06476 | 980,612777 |
| | Margaric acid | 17:1 n-7 | 77,8680895 | 110,178184 | 75,4166248 | 61,3921145 | 98,8358022 | 113,21834 | 76,3947838 | 76,8596363 |
| | Oleic ac | 18:1;n-9 | 4885,41101 | 6825,98769 | 5035,70376 | 3698,89366 | 8455,03398 | 9170,33524 | 7399,74973 | 6586,75301 |
| | 11-Eicosenoic ac | 20:1;n-9 | 91,2422263 | 145,034784 | 96,2992601 | 73,8195666 | 158,993313 | 198,148113 | 151,373245 | 126,261105 |
| | Erucic ac | 22:1;n-9 | 16,3991242 | 27,0759186 | 17,1578943 | 15,194329 | 30,1192404 | 38,3689963 | 29,6010462 | 25,8058094 |
| | Nervonic ac | 24:1;n-9 | 16,2702534 | 25,3214923 | 17,1884615 | 13,4065854 | 17,4759601 | 24,3211874 | 18,528921 | 14,6345022 |
| PUFA w-6 | Linoleic ac | 18:2;n-6 | 616,378493 | 839,572204 | 654,49917 | 404,365514 | 303,594157 | 357,219213 | 301,995202 | 243,262768 |
| | Gamma-Linolenic ac | 18:3;n-6 | 68,9919005 | 77,8166205 | 69,690898 | 31,462391 | 23,2904762 | 26,5637614 | 20,7311533 | 17,6010195 |
| | Eicosadienoic ac | 20:2;n-6 | 69,3942924 | 100,126997 | 78,6823671 | 57,499142 | 76,3595103 | 94,8519189 | 76,2245709 | 61,372477 |
| | Dihomo-g-linolenic ac | 20:3;n-6 | 178,287988 | 245,127343 | 215,346564 | 137,123934 | 119,511189 | 135,68818 | 126,276487 | 108,255565 |
| | Arachidonic ac | 20:4;n-6 | 764,538827 | 983,632454 | 795,024552 | 531,486043 | 165,849598 | 199,971914 | 165,3271 | 153,89882 |
| | Docosapentaenoic acid | 22:2 n-6 | 3,43711565 | 4,62843195 | 3,69230409 | 3,13378251 | 3,33579469 | 3,58749381 | 2,9663174 | 2,42588078 |
| | Docosatetraenoic ac | 22:4;n-6 | 185,696169 | 207,39945 | 186,103789 | 104,081294 | 72,3143156 | 89,2033082 | 92,9758872 | 94,7897177 |
| | Docosapentaenoic ac | 22:5;n-6 | 45,8918791 | 47,1754252 | 34,1312586 | 24,1079776 | 10,4870732 | 11,2211374 | 11,1993125 | 11,0019722 |
| PUFA w-3 | Alpha-Linolenic ac | 18:3;n-3 | 2,76333634 | 4,45393018 | 3,80562435 | 6,12223766 | 2,5818248 | 3,42356035 | 5,89787751 | 2,6832438 |
| | Octadecatetraenoic ac | 18:4 n-3 | 1,16027078 | 1,1722637 | 1,26887318 | | 0,77902231 | 0,73610617 | | |
| | eicosatetraenoic acid | 20:4;n-3 | 2,46766747 | 3,61255758 | 2,91885013 | 1,60658847 | 2,18915559 | 2,15238881 | 1,99930672 | 1,76410095 |
| | Eicosapentaenoic ac | 20:5;n-3 | 59,3726984 | 90,4236482 | 62,9819825 | 45,7776681 | 53,9052916 | 59,5087642 | 44,1207364 | 36,4833406 |
| | Docosapentaenoic ac | 22:5;n-3 | 280,804786 | 362,29601 | 317,467968 | 239,801856 | 260,190613 | 288,705651 | 267,294557 | 257,375362 |
| | Docosahexaenoic ac | 22:6;n-3 | 211,926322 | 280,195556 | 241,455876 | 196,41616 | 236,293394 | 271,338277 | 247,816946 | 230,711003 |

| | | treatment | dMIX FA | | | | dMIX FA DGATi | | | | dMIX FA ATGLi | | | |
|---|---|---|---|---|---|---|---|---|---|---|---|---|---|---|
| | | | #1 | #2 | #3 | #4 | #1 | #2 | #3 | #4 | #1 | #2 | #3 | #4 |
| SFA | octanoic acid | 8:00 | 178,990264 | 192,598569 | 227,174244 | 214,47753 | 195,3244 | 230,443571 | 214,617324 | 199,90939 | 185,970709 | 285,118173 | 218,357207 | 229,05514 |
| | Decanoic acid | 10:00 | 8,59633134 | 8,41194251 | 10,1679515 | 11,1940977 | 9,67655689 | 8,24408971 | 9,61878824 | 10,3361726 | 10,5735959 | 11,852245 | 9,47379461 | 13,5366176 |
| | Undecanoic acid | 11:00 | 2,32371247 | 2,61493125 | 2,09578654 | 2,34649435 | 2,6516995 | 2,03628519 | 2,27098236 | 2,23253318 | 2,44692773 | 2,28987924 | 1,99557393 | 2,54853952 |
| | Lauric acid | 12:00 | 21,3427587 | 23,171121 | 22,8729371 | 24,6736384 | 23,9645302 | 21,1012248 | 17,984503 | 19,1235097 | 24,2913572 | 23,0992601 | 20,1269102 | 22,1995328 |
| | Tridecylic acid | 13:00 | 1,96016822 | 2,48360426 | 1,82699032 | 2,3271794 | 2,37973592 | 2,73696643 | 2,4211064 | 2,16205913 | 2,03582703 | 2,16916208 | 2,28924346 | 2,06371999 |
| | Myristic acid | 14:00 | 263,591594 | 415,858227 | 371,877468 | 276,868802 | 371,825068 | 381,156747 | 296,188574 | 264,100379 | 333,962371 | 437,873897 | 372,782298 | 320,296088 |
| | Pentadecylic acid | 15:00 | 7,07480064 | 9,64864591 | 8,44502465 | 6,59613963 | 8,86875439 | 9,60466547 | 7,59205246 | 6,98567837 | 7,56397653 | 9,33429008 | 8,06565393 | 7,55451206 |
| | Palmitic acid | 16:00 | 1776,78994 | 2723,15764 | 2469,72779 | 1806,85105 | 2285,77281 | 2609,40335 | 2126,39837 | 1761,0216 | 2267,67298 | 2876,40015 | 2461,52321 | 2261,95291 |
| | Margaric acid | 17:00 | 48,8708646 | 70,7710177 | 59,5478034 | 46,5435701 | 51,8260281 | 55,1247319 | 47,5560055 | 40,3951074 | 52,9173081 | 64,8909244 | 55,58307 | 52,2467133 |
| | Stearic acid | 18:00 | 1402,90842 | 2160,76136 | 1914,79787 | 1366,56294 | 1511,34216 | 1674,28577 | 1339,62037 | 1050,04554 | 1531,22108 | 1968,70408 | 1794,3362 | 1531,75673 |
| | Arachidic ac | 20:00 | 13,0919086 | 22,3814705 | 20,2634152 | 15,1317198 | 12,7047644 | 14,7375427 | 13,7921105 | 12,1873119 | 15,1378638 | 19,5051901 | 18,2582246 | 17,4430153 |
| | Behenic ac | 22:00 | 16,3776072 | 30,1766465 | 22,1457047 | 12,9732631 | 19,0670225 | 23,698624 | 15,2450264 | 9,42730613 | 20,4915876 | 25,7245207 | 20,6122567 | 17,764463 |
| | Lignoceric ac | 24,00 | 10,2263089 | 18,2819129 | 14,9139807 | 8,6424111 | 11,0656969 | 13,448413 | 12,5001178 | 8,28535461 | 10,9600233 | 14,5653439 | 14,0269177 | 11,2694909 |
| MUFA | Myristoleic ac | 14:1;n-5 | 4,27697522 | 4,80129093 | 5,44483162 | 4,12339882 | 6,18959877 | 6,14470929 | 4,29025846 | 4,76278971 | 5,39929248 | 6,23621472 | 5,32307141 | 4,92391108 |
| | Sapienic acid | 16:1;n-10 | 1005,09594 | 1544,43626 | 1879,68374 | 1205,02092 | 1301,38487 | 1404,5079 | 1632,17498 | 1144,73073 | 1142,43887 | 1563,84338 | 1734,25349 | 1307,42736 |
| | Palmitoleic ac | 16:1;n-7 | 1062,1151 | 1683,85521 | 1611,46922 | 1170,87436 | 1450,38744 | 1567,21055 | 1476,72084 | 1137,21514 | 1269,98785 | 1758,29336 | 1567,16469 | 1275,48777 |
| | Margaric acid | 17:1 n-7 | 82,065804 | 118,905786 | 104,056108 | 85,5269849 | 93,4097905 | 97,2303259 | 94,7848028 | 80,9094113 | 92,2058718 | 113,465784 | 103,590689 | 91,9907708 |
| | Oleic ac | 18:1;n-9 | 5045,66735 | 6995,12669 | 7027,51103 | 5016,2996 | 5504,82137 | 5941,88863 | 5420,62137 | 4118,32605 | 5582,95331 | 6794,86502 | 6389,76666 | 5235,13462 |
| | 11-Eicosenoic ac | 20:1;n-9 | 95,0712179 | 147,398798 | 144,572769 | 98,1380712 | 100,733542 | 102,862203 | 88,1606203 | 72,6226123 | 101,837869 | 137,550695 | 135,79861 | 106,600184 |
| | Erucic ac | 22:1;n-9 | 18,0457839 | 29,031942 | 25,4649841 | 18,1120245 | 17,5991109 | 19,0002886 | 17,2579803 | 12,9827372 | 18,7804675 | 24,5160173 | 25,8000602 | 21,6246502 |
| | Nervonic ac | 24:1;n-9 | 15,2675536 | 23,6715974 | 21,6738285 | 13,7352763 | 14,9886824 | 16,7303009 | 15,1163877 | 11,5741734 | 15,5979646 | 20,7489021 | 22,1373563 | 18,2333975 |
| PUFA w-6 | Linoleic ac | 18:2;n-6 | 899,985532 | 1159,8329 | 1388,96293 | 1151,5166 | 846,336124 | 933,161304 | 953,679459 | 590,236303 | 1047,63531 | 1182,24332 | 1257,75863 | 927,808626 |
| | Gamma-Linolenic ac | 18:3;n-6 | 100,240541 | 111,10926 | 88,3563452 | 80,6861158 | 309,318075 | 381,052457 | 281,042447 | 93,8176597 | 105,911769 | 120,030095 | 77,3335567 | 52,3529368 |
| | Eicosadienoic ac | 20:2;n-6 | 74,6790012 | 112,076051 | 122,755459 | 85,2331975 | 80,7215145 | 83,67399 | 68,997398 | 57,8102647 | 80,6527747 | 104,701051 | 118,54533 | 91,8617518 |
| | Dihomo-g-linolenic ac | 20:3;n-6 | 229,171731 | 301,741559 | 363,301288 | 262,72684 | 184,980512 | 183,247764 | 198,819427 | 168,300535 | 262,763375 | 297,745186 | 320,904293 | 251,647161 |
| | Arachidonic ac | 20:4;n-6 | 293,048714 | 395,916675 | 329,536701 | 266,014218 | 424,50828 | 350,80536 | 276,638099 | 266,357712 | 446,512463 | 553,613937 | 436,753811 | 391,241707 |
| | Docosapentaenoic acid | 22:2 n-6 | 3,64586086 | 4,94339375 | 4,98468623 | 4,03178644 | 3,69528058 | 3,72058894 | 3,23114998 | 2,91151051 | 4,15034523 | 5,04305615 | 5,0575035 | 4,63223138 |
| | Docosatetraenoic ac | 22:4;n-6 | 130,709605 | 192,249677 | 200,328112 | 168,76875 | 101,42067 | 148,007579 | 119,430225 | 196,876562 | 199,147519 | 235,788393 | 265,627918 | 241,67245 |
| | Docosapentaenoic ac | 22:5;n-6 | 16,7245397 | 22,3267666 | 18,3309331 | 15,8121134 | 10,9130913 | 12,7586789 | 11,7844887 | 14,93436 | 22,749826 | 23,7497748 | 22,4985553 | 23,6068957 |
| PUFA w-3 | Alpha-Linolenic ac | 18:3;n-3 | 11,9944893 | 15,1548993 | 20,9691478 | 26,9901569 | 9,7352649 | 10,1008226 | 12,643508 | 7,78930151 | 12,7641352 | 16,6857241 | 19,6644083 | 12,9069351 |
| | Octadecatetraenoic ac | 18:4 n-3 | 1,70087863 | 1,7889895 | 1,65064751 | 1,48543824 | 9,26755928 | 12,1420455 | 7,8308404 | 2,11773095 | 1,86716022 | 2,22624134 | 1,67232685 | 1,02892414 |
| | eicosatetraenoic acid | 20:4;n-3 | 5,7052904 | 7,01332232 | 8,64654481 | 7,55672913 | 4,04731916 | 4,00956242 | 5,959841 | 4,5506362 | 6,32940965 | 6,67554948 | 8,6354035 | 6,10403183 |
| | Eicosapentaenoic ac | 20:5;n-3 | 62,844509 | 83,5695943 | 66,5085261 | 50,3362741 | 96,234405 | 89,4648958 | 74,3497894 | 44,1131563 | 65,8001893 | 81,9453226 | 61,0106302 | 46,8081114 |
| | Docosapentaenoic ac | 22:5;n-3 | 271,229016 | 363,67243 | 437,978526 | 336,143513 | 293,186905 | 303,697135 | 295,332599 | 278,26425 | 309,145887 | 362,443845 | 395,627126 | 352,751248 |
| | Docosahexaenoic ac | 22:6;n-3 | 213,579897 | 282,257868 | 324,884357 | 251,819053 | 271,826967 | 276,754122 | 267,045653 | 231,309649 | 233,024112 | 277,310589 | 293,977531 | 261,261677 |
